# Genomic remnants of ancestral hydrogen and methane metabolism in Archaea drive anaerobic carbon cycling

**DOI:** 10.1101/2021.08.02.454722

**Authors:** Panagiotis S. Adam, George E. Kolyfetis, Till L.V. Bornemann, Constantinos E. Vorgias, Alexander J. Probst

**Author notes:** These authors contributed equally (Panagiotis S. Adam, George E. Kolyfetis).

## Abstract

Methane metabolism is among the hallmarks of Archaea, originating very early in their evolution. Other than its two main complexes, methyl-CoM reductase (Mcr) and tetrahydromethanopterin-CoM methyltransferase (Mtr), there exist other genes called “methanogenesis markers” that are believed to participate in methane metabolism. Many of them are Domains of Unknown Function. Here we show that these markers emerged together with methanogenesis. Even if Mcr is lost, the markers and Mtr can persist resulting in intermediate metabolic states related to the Wood-Ljungdahl pathway. Beyond the markers, the methanogenic ancestor was hydrogenotrophic, employing the anaplerotic hydrogenases Eha and Ehb. The selective pressures acting on Eha, Ehb, and Mtr partially depend on their subunits’ membrane association. Integrating the evolution of all these components, we propose that the ancestor of all methane metabolizers was an autotrophic H_2_/CO_2_ methanogen that could perhaps use methanol but not oxidize alkanes. Hydrogen-dependent methylotrophic methanogenesis has since emerged multiple times independently, both alongside a vertically inherited Mcr or from a patchwork of ancient transfers. Through their methanogenesis genomic remnants, Thorarchaeota and two newly reconstructed order-level lineages in Archaeoglobi and Bathyarchaeota act as metabolically versatile players in carbon cycling of anoxic environments across the globe.

## Introduction

Until recently, all known methanogens were members of the Euryarchaeota and classified into two groups: Class I (Methanopyrales, Methanobacteriales, Methanococcales) and Class II (Methanosarcinales, Methanomicrobiales)^1^. While the composition of Class I methanogens (Methanomada) has remained constant over the years, several methane metabolizing lineages are now known to be related to the Class II methanogens. Collectively they form the clade called Methanotecta^2^ or Halobacterota^3,4^ and include the Methanocellales, Methanoflorentaceae^5^, Methanonatronarchaeia^6^, Methanophagales (ANME-1), and Archaeoglobi^7–9^. The distribution of methane metabolism currently extends to most Euryarchaeota major clades and some Proteoarchaeota lineages^10,11^. Inferring methane metabolism from metagenomic data is tied to the presence of Mcr that catalyzes the reversible reduction of CoM-attached methyl to methane. The concomitant presence of the Mtr complex implies H_2_/CO_2_ methanogenesis or anaerobic methane oxidation (AMO), for instance in Nezhaarchaeota^8^ and Verstraetearchaeota^12^. Methylotrophic methanogenesis through methanol, methylamine, and methylthiol methyltransferases has been discovered in several lineages: Methanomassiliicoccales, Methanofastidiosales^13^, Nuwarchaeales (NM3)^10,14^, Verstraetearchaeota^7,8,10,15^, Korarchaeota^8,10,16^, and Thaumarchaeota^7^. The phylogeny of Mcr is only partially congruent with the archaeal species tree^11^. A divergent Mcr-like clade has been associated with anaerobic alkane oxidation (AAO) across Archaea (Syntropharchaeales^17^, Methanoliparia^10^, Methanosarcinales^10,18,19^, Bathyarchaeota^10,20^, Helarchaeota^21^, Hadesarchaea^7,8^, Archaeoglobi^8,22^). A series of other genes have been dubbed methanogenesis markers, by virtue of their taxonomic distribution matching that of methanogenesis and AMO. However, they are rarely used in metabolic annotations. It has been proposed that the presence of these genes outside methane- or alkane-metabolizing lineages could indicate that they are metabolic remnants repurposed into other pathways^10,23,24^.

While Mcr is never encountered outside of methane metabolism and AAO, there exist several archaeal and bacterial lineages that possess MtrAH, the two methyltransferase subunits. The role of these methyltransferase subunits in other types of metabolism is currently unclear. Another long-standing debate concerns how ancient methane metabolism is among Archaea^11^ and whether its original form was H_2_/CO_2_ (hydrogenotrophic, carbon dioxide reducing)^12^ or (hydrogen-dependent) methylotrophic^14^. The role of methanogenesis markers in methane/alkane metabolism and their origins are largely unstudied as well. Many of them are Domains of Unknown Function (DUFs)^25^, mirroring the large number of DUFs among the auxiliary genes of the Wood-Ljungdahl pathway (WLP)^26^. In this study, we set out to address these questions, starting from the evolution and metabolic roles of methanogenesis markers. We leveraged a combination of phylogenomic and metagenomic methods to determine the metabolic and ecological functions of methanogenesis markers as remnants of methane metabolism. Through computational biophysics methods, we explored the evolutionary mechanisms that drive the emergence and breakdown of methanogenic complexes and their related hydrogenases.

## Results & Discussion

Different sets of methanogenesis markers in the literature, such as the ones in Gao & Gupta^23^ and Borrel et al.^10^, do not always contain the same genes. Nevertheless, many markers are known to have partial taxonomic distributions among methanogens and/or consist of DUFs. We suspected that there might exist additional potential markers. Thus, we began by surveying the taxonomic distribution of archaeal DUFs, looking for co-occurrences with methane metabolism. First, we defined a DUF as “archaeal” if at least half of its distribution in Uniprot consisted of Archaea. From the distributions and phylogenies of the 155 archaeal DUFs identified (Supplementary Data 1), we distinguished two categories relevant to methane metabolism: 1) DUFs among the 38 methanogenesis markers of Borrel at al.^10^ with a generally broad distribution across methane metabolizers; 2) other DUFs distributed mainly in methane metabolizers but covering their range partially.

To date, there exist no phylogenies of most methanogenesis markers. For that reason, before considering the DUFs further, we first reconstructed the phylogenies of all 38 markers from the Borrel set (Supplementary Table 1)^10^, regardless if they were DUFs or not (Figure 1, Supplementary Figures 1-30, Supplementary Data 2). In the case of Mcr and Mtr, we created supermatrices of McrABG (Supplementary Data 3) and MtrABCDEFG (Supplementary Data 4), tested for congruence, and constructed phylogenies from the concatenated datasets. MtrFG were included despite not being part of the marker list, since their distribution matched the other subunits. MtrA comprises homologs in both Bacteria and Archaea outside of the monophyletic canonical clade of methanogens and ANME (Supplementary Figure 31). These might function as methyltransferases together with MtrH^27^. MtrH itself was omitted since it had numerous isolated homologs and interdomain transfer events that complicated its phylogeny (Supplementary Figure 32). A specific Bathyarchaeota clade (Subgroups 20&22) contains vertically inherited and complete Mtr clusters without Mcr. Many Thorarchaeota possess vertically inherited canonical MtrAH (Supplementary Figures 31, 32) without the other subunits, acting as a methyltransferase in their proposed mixotrophic lifestyle^28^. The topology of MtrA here is in agreement with other recent phylogenies^29^. Both Mcr and Mtr can be traced to the common ancestor of Euryarchaeota and Proteoarchaeota. Further tracing these complexes to the Last Archaeal Common Ancestor is problematic, since Mcr and Mtr are absent in Altiarchaeota and thus, unlike the components of the Wood-Ljungdahl pathway (WLP), we have to disregard the DPANN or assume a massive loss event. For convenience and to address the uncertainty in the earliest appearance of methane metabolism, we refer to the ancestral methane metabolizing archaeon as “Last Methane (metabolizing) Ancestor” (LMA).

**Figure 1.**
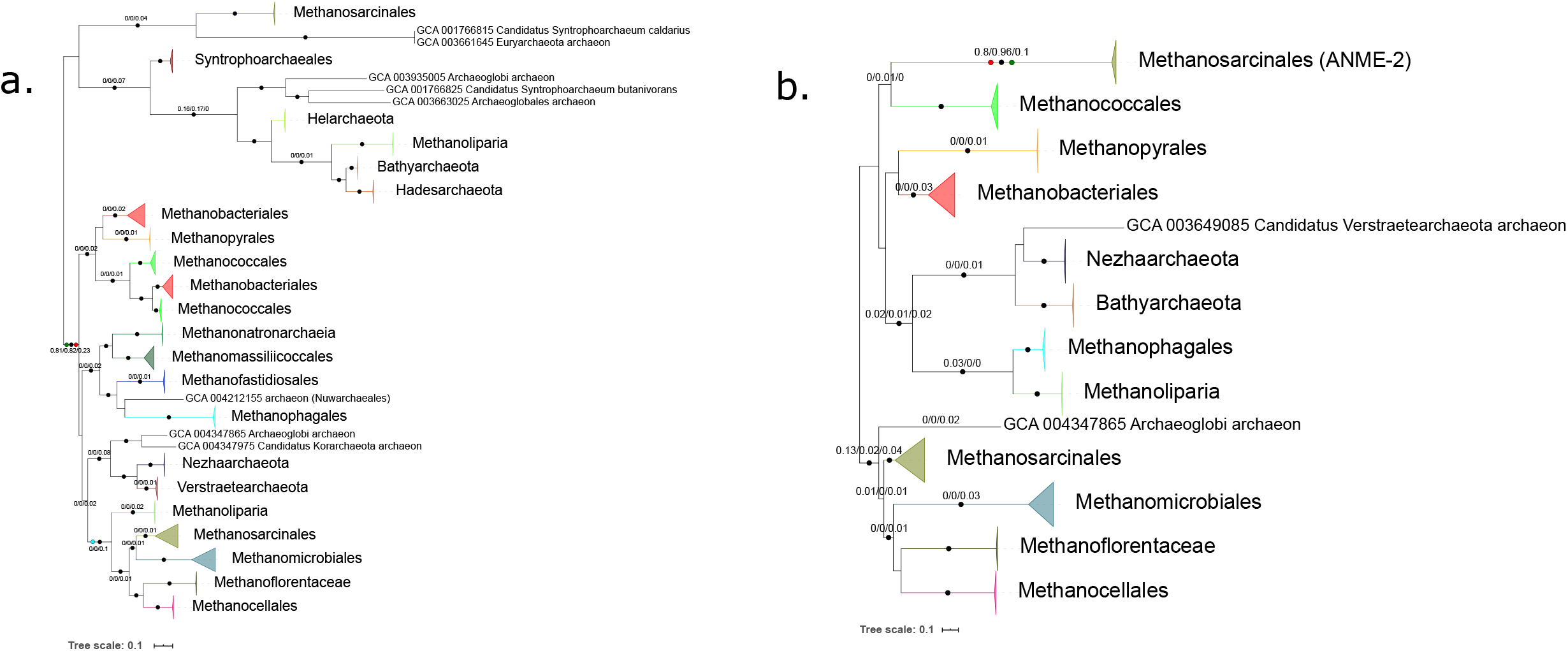
Evolution of the Mcr and Mtr complexes. Maximum Likelihood (ML) phylogenies of (a) McrABG (1141 aa positions), (b) MtrABCDEFG (1079 aa positions). Black circles indicate strongly supported branches (ultrafast bootstrap >=95, aLRT SH-like >=80), red circles correspond to the MAD root, green to MinVar, and blue to NONREV. Branch values correspond to rootstrap supports for MAD, MinVar, and NONREV respectively. In Mtr the NONREV root is within a collapsed clade. The NONREV rooting was spurious and its rootstrap supports were low, particularly for Mtr, Eha, and Ehb, probably due to the small number of positions in the supermatrices; it was mostly included to compare with the other rooting algorithms.

Eight of the remaining 30 methanogenesis markers (m15, m16, m17, m18, m25, m26, m32, m37) are present in non-methane-metabolizing lineages. Of those, only the evolution of m15, m16, and m17 had been previously examined^10^. None of our marker phylogenies are fully resolved. However, since individual lineages and even supergroups (e.g. Methanomada and Halobacterota) are monophyletic, the markers are probably as ancient as methane metabolism by proxy of Mcr. For non-methane metabolizers, the lack of resolution makes it difficult to distinguish which markers have been inherited vertically, and where lateral transfer events have occured. We established vertical inheritance by comparing the position of a lineage in the marker gene phylogeny with its position in the archaeal reference tree (Figure 2a). Such cases included the Theionarchaea in m16 (Supplementary Figure 13), Thorarchaeota in m26 (Supplementary Figure 23), non-alkanotrophic Bathyarchaeota with Mtr in m26 and m37 (Supplementary Figures 23, 29), and Hydrothermarchaeota in m32 (Supplementary Figure 24).

**Figure 2.**
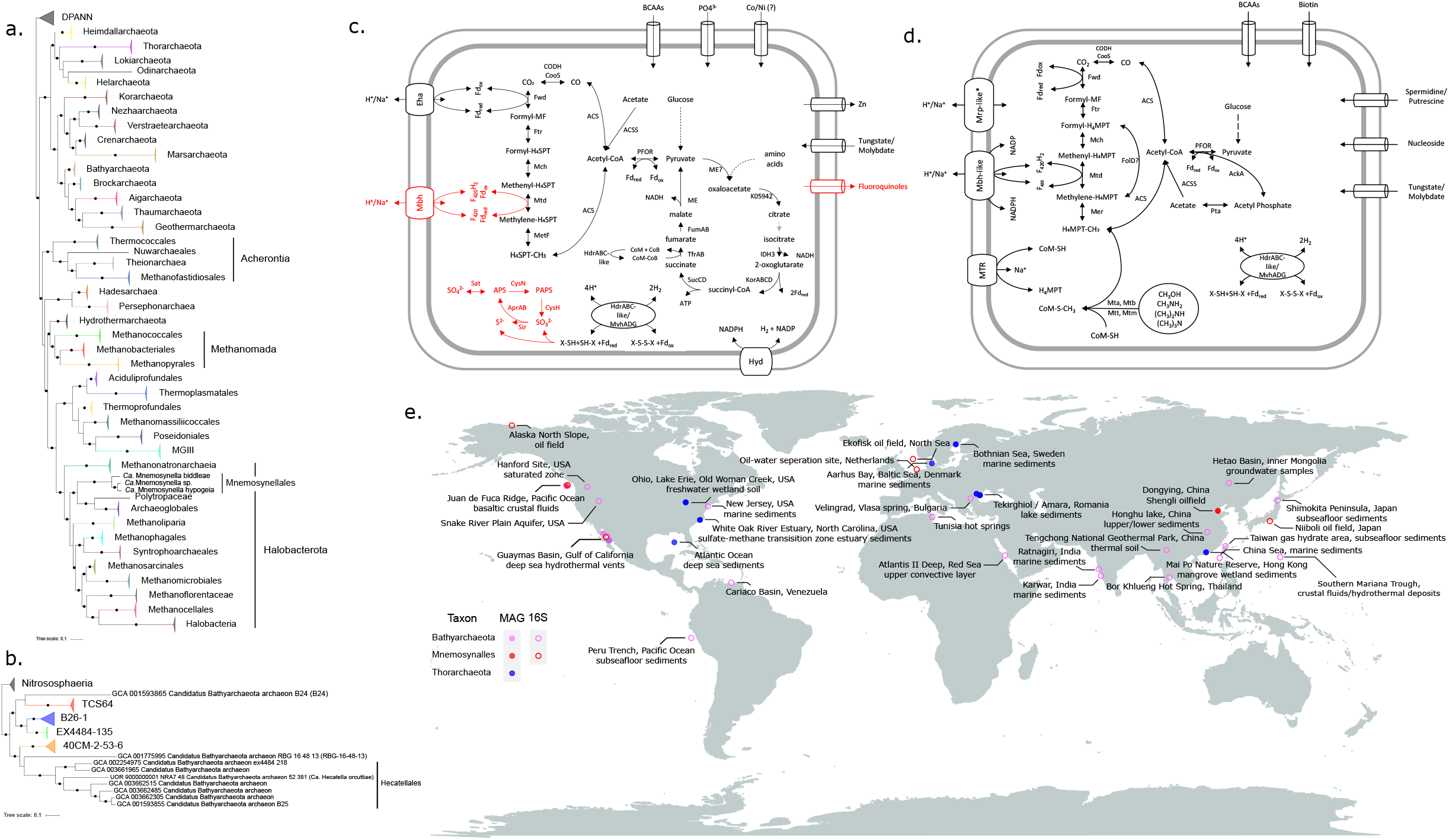
Systematics, metabolism, and biogeography of Mnemosynellales and Hecatellales. ML phylogenies of (a) Mnemosynellales within Archaea rooted at the DPANN (6021 aa positions), (b) Hecatellales in Bathyarchaeota rooted with Nitrososphaeria (7154 aa positions), based on the concatenation of 36 Phylosift markers. Black circles indicate strongly supported branches (ultrafast bootstrap >=95, aLRT SH-like >=80). Higher taxonomic groups mentioned in the text are named explicitly. Metabolic reconstructions for (c) *Ca.* Mnemosynella biddleae and *Ca.* Mnemosynella hypogeia, (d) *Ca.* Hecatella orcuttiae. Systems marked in red are found exclusively in *Ca.* M. biddleae, perhaps due to the higher quality and size of the genome. MF: methanofuran, H4MPT: tetrahydromethanopterin. (e) Biogeographic distribution of Mnemosynellales, Hecatellales, and Thorarchaeota with canonical MtrA. Coordinates/location and environment type were recovered from the respective WGS project metadata in NCBI and 16S entries in SILVA. For the reference tree of Archaea (a), we obtained in most phylogenies and usually with strong support a monophyletic clade of Hydrothermarchaeota with Methanomada. This is unlike GTDB that places Methanomada with Thermococci (here, Acherontia) in the phylum Methanobacteriota. Instead, we propose the name Phlegethonia for this superclass or superphylum that includes multiple thermophiles, following the Underworld river naming trend of Stygia and Acherontia^2^.

Our phylogenies of Mcr, Mtr, and methanogenesis markers are indicative of multiple independent losses of that metabolism, similar to previous observations about how the WLP has repeatedly been independently lost over archaeal clades^26,30^. It has recently been shown how the loss of the tetrahydromethanopterin branch and its auxiliary genes is often incomplete and tends to leave behind genomic remnants. These are usually biosynthetic genes of the tetrahydromethanopterin and methanofuran cofactors, but also genes of the main pathway (e.g., Mch in Halobacteriales)^26^. The same situation has been proposed to apply to methane metabolism^10^, and we see here that it is very pervasive across methanogenesis markers. Remnants become more common, if we relax the requirement for vertical inheritance to only one marker per lineage and attribute the rest to lack of resolution, e.g., Hydrothermarchaeota (m15, m17, m32), Theionarchaea (m16, m18), Lokiarchaeota (m15, m17, m18, m25), Subgroup 20&22 Bathyarchaeota (m17, m18, m26, m37).

Lineages with methane metabolism remnants can be regarded as former methanogens/methanotrophs, although loss of alkanotrophy could result in such patterns, too. Since the taxa listed above retain the WLP^2,26^, and anaerobic methanotrophy generally seems to be a derived trait across separate Halobacterota clades, they probably used to be H_2_/CO_2_ methanogens. The link between the WLP and Mcr through Mtr persists in progressive intermediate loss stages in the Subgroup 20&22 Bathyarchaeota and Thorarchaeota, making this inference more robust. Other than m26, which is tied to Mtr, we could neither identify reasons for the conservation discrepancies among markers and among lineages, nor confidently infer the repurposed function of uncharacterized remnant markers from genomic context or otherwise. For our observations on the evolution and functional annotation of the markers, see the Supplementary Information.

The NCBI taxon “Euryarchaeota archaeon JdFR-21”, possesses five methanogenesis markers, more than any other non-methane/alkane metabolizing archaeon. JdFR-21 is a member of the Archaeoglobi related NRA7 clade and was recovered from subsurface fluid metagenomes of the Juan de Fuca Ridge^31^, like the alkane oxidizer *Ca.* Polytropus marinifundus^22^ (formerly JdFR-42). The JdFR metagenomes also contain JdFR-11, one of the Bathyarchaeota with canonical Mtr. We found two additional NRA7 MAGs (Archaeoglobi MAG-15, Archaeoglobi MAG-16) from the Shengli oil field metagenomes^32^ that were submitted to NCBI after we created our local genomic databases, and were thus not included in other phylogenomic analyses. We downloaded the JdFR and Shengli metagenomic reads from SRA, reassembled, rebinned them, and manually curated the genomes, improving upon their NCBI counterparts. The refined bins corresponding to JdFR-21, MAG-16, and JdFR-11 fulfill the quality criteria for being classified as high-quality genomes^33^. We then determined their taxonomic placement trying to account for various sources of bias in the phylogenies, to clarify how they fit in the history of methane metabolism (Supplementary Data 5). The two JdFR MAGs are the highest-quality representatives of their respective order-level lineages (Figure 2a, 2b, Supplementary Table 2). The taxonomic delineation was confirmed by pairwise Average Nucleotide Identity (ANI) and Average Amino acid Identity (AAI) comparisons (Supplementary Figure 33). We propose the names *Ca.* Mnemosynella biddleae for JdFR-21, *Ca.* Mnemosynella hypogeia for MAG-16 (order: Mnemosynellales), and *Ca.* Hecatella orcuttiae (order: Hecatellales) for JdFR-11. Full genome statistics and proposed nomenclature for all MAGs binned in this study are given in Supplementary Table 2.

All Mnemosynellales possessed the same methanogenesis markers (Supplementary Data 2). Based on their phylogenetic position as the basal divergence of Archaeoglobi, we could infer that the markers were vertically inherited, making Mnemosynellales former H_2_/CO_2_ methanogens. Their metabolisms (Figure 2c, Supplementary Data 6) revolve around the WLP but defining whether it is oxidative or reductive is problematic. They can oxidize acetate and possess a Hyd-like hydrogenase for hydrogen evolution, but they also encode the anaplerotic Eha hydrogenase of H_2_/CO_2_ methanogens. Mnemosynellales appear capable of performing most TCA cycle reactions other than the steps from malate to oxaloacetate and citrate to isocitrate. The succinate to fumarate conversion was predicted to be catalyzed by the CoM and CoB-forming thiol:fumarate reductase that is syntenic to an Hdr-like (heterodisulfide reductase) complex and Eha. Originally characterized in Methanobacteriales^34^, such fumarate reductases have been proposed to function in Natranaeroarchaeales^35^ and some Asgard lineages^36^. The CoM-CoB heterodisulfidic bond could be regenerated by the Hdr-like complex, with a reductive WLP functioning as an electron sink. The underlying assumption here is a source of oxaloacetate, perhaps from amino acid fermentation or from pyruvate through the oxaloacetate-decarboxylating malate dehydrogenase, since pyruvate carboxylase was not found. Nonetheless, it is possible that the TCA reactions run in the opposite direction through reducing potential from hydrogen or sulfur species. Additionally, both genomes encoded an Hdr/Mvh-like complex that could function on CoM-CoB or polysulfides. The *Ca.* M. biddleae genome also contains genes for an Mbh/Mrp-like hydrogenase and both assimilatory and dissimilatory sulfur metabolism, where Hdr/Mvh could perform thiosulfate or other heterodisulfide disproportionation.

Hecatellales (Bathyarchaeota subgroups 20&22) include the B25 MAG that has been proposed to be an acetogen^37^. *Ca.* H. orcuttiae (Figure 2d, Supplementary Data 6) seems to have the capacity for acetogenesis running the WLP reductively, but also acetate assimilation and transferring methyl moieties from methanol and methylamines into an oxidative WLP through Mtr. Membrane potential is probably generated by an Mbh/Mrp-like hydrogenase regulated by an additional Mrp antiporter that is syntenic to the formylmethanofuran dehydrogenase, Fwd. *Ca.* H. orcuttiae might perform hydrogen dependent heterodisulfide disproportionation via an Hdr/Mvh-like complex, similar to Mnemosynellales. The Bathyarchaeota member CR_14 (not in our datasets) branches within order B26-1 and contains a complete canonical Mtr that has been suggested to link methylated compounds to the WLP^38^. The presence of Mtr outside Hecatellales further corroborates our inference of ancestral H_2_/CO_2_ methanogenesis in Bathyarchaeota.

In terms of biogeography (Figure 2e, Supplementary Figure 34, Supplementary Data 5), Mnemosynella is the only known genus in the order and is found globally in oil fields. It includes a divergent geothermal clade found exclusively in the Eastern Pacific but the phylogeny is not adequately resolved to determine its origin (Supplementary Figure 34a). Hecatellales MAGs have only been found in geothermal environments in the Eastern Pacific but from their 16S rRNA gene sequences we can deduce that they are present in many types of mainly high temperature environments around the world, into which metagenome sequencing efforts should be expanded (Supplementary Figure 34b). In contrast, the Thorarchaeota MAGs that utilize the canonical MtrA originate from a wide variety of anaerobic environments and localities. Due to the diversity of how methanogenesis remnants have been integrated in metabolism around the WLP, Mnemosynellales, Hecatellales, and Thorarchaeota can occupy multiple niches across diverse environments in the global carbon cycle.

Having finalized the phylogenies of the 38 methanogenesis markers, we turned our attention to the evolution of the partial markers among DUFs. We further expanded beyond DUFs searching for homologs and constructing phylogenies for all “proteins that are specific for methanogens” and “proteins that are specific to certain subgroups of methanogens” from Gao & Gupta^23^ (Supplementary Data 7). We could subdivide all these genes into two more categories: The first category comprises genes with a narrow distribution that could not confidently be inferred to be as ancient as Mcr and Mtr. Among others, in this category there were five genes found exclusively in Methanopyrales and Methanobacteriales. Three of them, based on synteny, are probably involved in pseudomurein biosynthesis (Supplementary Figure 35). Another case is the Hcg proteins in the biosynthetic pathway of the iron guanylylpyridinol cofactor of the Hmd hydrogenase in Methanomada and Desulfurobacteriales (Supplementary Figure 36, Supplementary Data 8, 9). The second category consists of genes whose origin can be traced to the LMA either under the classic root of Archaea with respectively monophyletic Proteoarchaeota and Euryarchaeota or with a root within Euryarchaeota from Raymann et al.^39^.

With few exceptions, most of these ancient genes are subunits of the Eha and Ehb anaplerotic hydrogenases that are known to provide electrons during methanogenesis and carbon fixation respectively in Methanomada^40–42^. The evolution of these hydrogenases and their relationship with methane metabolism outside Methanomada are mostly unknown. Thus, we also searched for homologs of any remaining subunits or expanded previous datasets by extrapolating the expected distribution (Supplementary Methods, Supplementary Data 8), tested for congruence, and concatenated them into supermatrices (Supplementary Data 10, 11). The Eha genes form a highly conserved cluster and they have evolved mainly vertically with some lineage-specific tinkering involving gain/loss of subunits or use of different ferredoxins (Figure 3a, Supplementary Results & Discussion). The exceptions are a possible ancient homologous recombination event affecting some Methanobacteriales and a transfer between Mnemosynellales and Persephonarchaea (MSBL1) (Figure 3a). Determining the direction of this transfer depends on the root of the phylogeny, as placed by outgroup-free rooting with Minimal Ancestor Deviation (MAD) and Minimum Variance (MinVar). Each scenario is supported by the phylogenies of a subset of WLP components (Supplementary Figures 37-41, Supplementary Data 7). A detailed analysis on the evolution of Eha is presented in the Supplementary Information. Eha can be traced to at least the ancestor of Euryarchaeota, corresponding to the LMA under the root from Raymann et al.^39^, or even earlier depending on the taxonomy of Persephonarchaea (Figure 2a).

**Figure 3.**
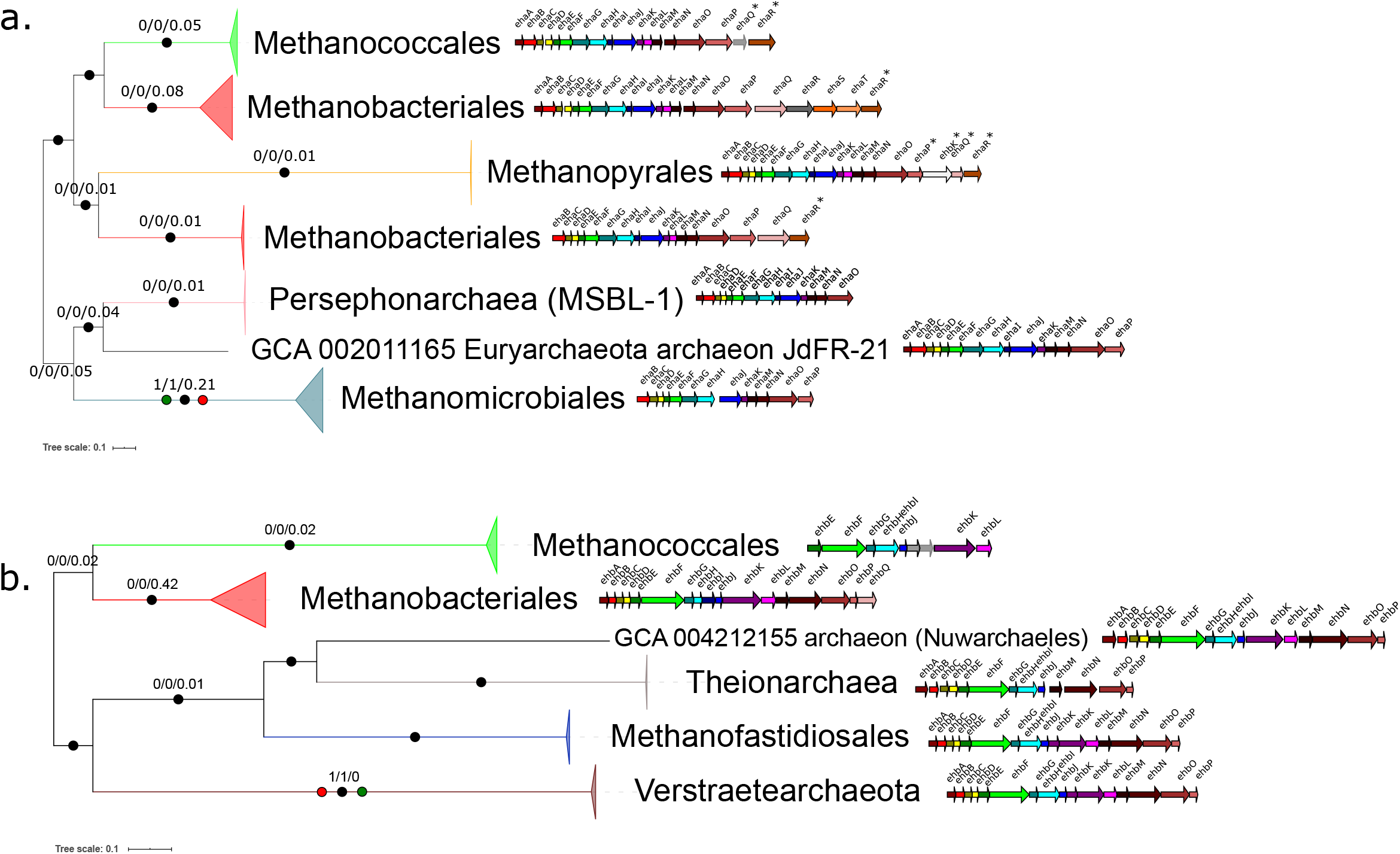
Evolution and comparative genomics of Eha and Ehb. ML phylogenies of (a) EhaBCDEFGHJLMNO (1828 aa positions), (b) EhbABCDEFGHIJKLMNOP (2770 aa positions), along with the genomic organization of the hydrogenase clusters in a representative genome for each major clade. Black circles indicate strongly supported branches (ultrafast bootstrap >=95, aLRT SH-like >=80), red circles correspond to the MAD root, green to MinVar. Branch values correspond to rootstrap supports for MAD, MinVar, and NONREV respectively. For both Eha and Ehb, the NONREV root is within a collapsed clade. Subunits marked with asterisks are problematic in terms of their homology and/or nomenclature (see Supplementary Information). The taxa used as illustrative cases for the cluster organization were: Methanothermobacter marburgensis str. Marburg (GCA_000145295; Methanobacteriales), Methanocaldococcus jannaschii DSM 2661 (GCA_000091665; Methanococcales), Methanopyrus kandleri AV19 (GCA_000007185; Methanopyrales), Methanothermobacter tenebrarum (GCA_003264935; Methanobacteriales small clade in Eha), Methanospirillum hungatei JF-1 (GCA_000013445; Methanomicrobiales), Euryarchaeota archaeon JdFR-21 (GCA_002011165; NRA7/Mnemosynellales), Candidate division MSBL1 archaeon SCGC-AAA259E19 (GCA_001549095; MSBL1/Persephonarchaea), *Candidatus* Methanomethylicus mesodigestum (GCA_001717035; Verstraetearchaeota), Arc I group archaeon ADurb1013_Bin02101 (GCA_001587595; Methanofastidiosales), Theionarchaea archaeon DG-70-1 (GCA_001595815; Theionarchaea), archaeon (GCA_004212155; Nuwarchaeales).

The evolution of Ehb (Figure 3b) is more complicated than Eha. Beyond lineage-specific tinkering, such as the loss of EhbKL in Theionarchaea, the signal among subunits is inconsistent, resulting in different topologies that are rarely strongly supported, often affecting the position of Methanococcales (Supplementary Data 11). The Ehb genes form a highly conserved cluster, except for Methanococcales where the genes encoding subunits EhbEFGHIJKL and sometimes EhbMO are co-localized and separate from the rest. Furthermore, EhbHI are fused similar to Acherontia and Verstraetearchaeota. This is probably the result of a massive homologous recombination event related to the Acherontia (Supplementary Figures 42-44, Supplementary Data 13; see Supplementary Information for a detailed description). Outgroup-free rooting (Supplementary Data 12) placed the root at Verstraetearchaeota, corresponding to a split between Euryarchaeota and Proteoarchaeota and Ehb having been present in the LMA.

In the membrane-associated complexes Eha, Ehb, and Mtr, many subunits are distinct protein families apparently emerging at the LMA. Unlike generic ion translocation and hydrogenase subunits, they are exclusively associated with these complexes. To determine how such subunits could have become established, we tested for selective pressures in the complexes. We calculated the site-specific evolutionary rates of Mcr, Mtr, Eha, and Ehb subunits as a selection proxy, following Sydykova & Wilke^43^. For both Eha and Ehb, significant differences were found within each complex (Kruskal-Wallis, p [2.3E-5 - 2E-2]). However, they were hard to pinpoint, since there were no subunits with consistently significantly different rates (Supplementary Figure 45, Supplementary Data 14). One exception was weakly significant (Dunn’s test and/or pairwise Mann-Whitney, q (false-discovery corrected p-value) <0.05) lower rates in the catalytic hydrogenase subunits EhbMN (Supplementary Figure 45j-l). Apart from a few outliers, the positions in all subunits are under neutral or weakly purifying selection, although our using trimmed alignments probably excludes some divergent positions.

We then tested whether predicted transmembrane segments undergo different selection compared to the extramembrane positions of the subunits. Our hypothesis was that the transmembrane regions would be subject to stronger purifying selection, due to being buried and/or in contact with other subunits^44^ and/or forming functional features (e.g., ion translocators in EhaHIJ^42^, EhbF^40,41^, MtrE^45^ or MtrCDE^46^). Nevertheless, there was no significant difference between transmembrane and extramembrane positions for most subunits (Figure 4). Where such a difference existed (Mann-Whitney, p [6.2E-12 - 3E-2]), it was extramembrane residues that had lower rates and were under purifying selection (exception: EhaE). The transmembrane segments were mostly under neutral selection. Any correlations between a position’s predicted transmembrane probability and rate, even if significant (Pearson correlation, p [7.8E-12 - 4.7E-2), were moderate or weak (generally |r|<=0.4, Supplementary Data 14) indicating that other structural features (solvent accessibility, flexibility, packing) and functional conservation contribute to selective pressure on these complexes, too.

**Figure 4.**
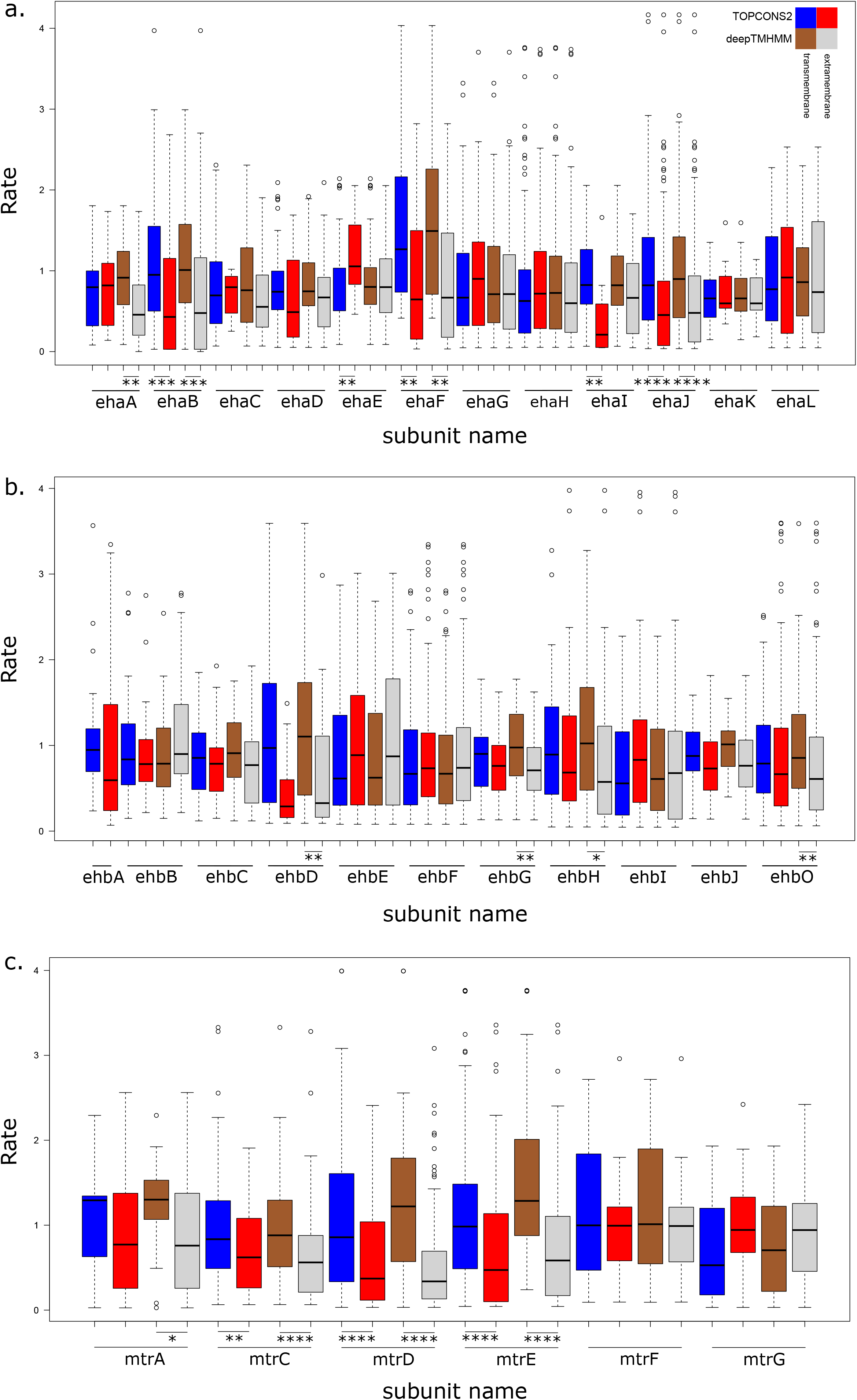
Selective pressure comparison between membrane bound and extramembrane residues of Eha, Ehb, and Mtr. Boxplots for site-specific empirical bayesian rates calculated under Poisson+G16 for each predicted transmembrane subunit of (a) Eha, (b) Ehb, (c) Mtr, split between transmembrane and extramembrane residues as predicted by TOPCONS2 and DeepTMHMM. Asterisks denote statistical significance (* <5E-2, ** <1E-2, *** <1E-3, **** <1E-4).

The metabolism of the methanogenic ancestor has been long debated with arguments in favor of both H_2_/CO_2_^12^ and hydrogen-dependent methylotrophic^14^ methanogenesis. Along with previous work placing the WLP at the common ancestor of Archaea^2,26^, we have established here the presence of Mcr, Mtr, Eha, and Ehb at the LMA. This suggests that it was at least an H_2_/CO_2_ methanogen fixing carbon by means of the WLP. Eha and Ehb form sister clades among group 3 [Ni-Fe] hydrogenases^41^. Since they both provide electrons to the initial reduction of CO_2_ to formylmethanofuran, they most probably arose from a duplication and subsequent tinkering separating carbon fixation from methanogenesis in the LMA’s lifestyle. While AMO is a possibility, the reversal of methanogenesis seems to be a derived trait emerging independently in Halobacterota clades. The origins of ANME Mcr and Mtr are often not in agreement and not inside Halobacterota, suggesting lateral transfers (Figure 1). However, it remains ambiguous how methylotrophic methanogenesis fits in the picture and how anaerobic alkane oxidation emerged. To address the first issue, we constructed phylogenies of the methyltransferase subunits MtaB (methanol), MtmB, MtbB, and MttB (mono-, di-, trimethylamine). Despite multiple putative ancient transfer events, MtaB could have been present in the ancestor of Euryarchaeota and perhaps the LMA (Supplementary Figure 46). For methylamine methyltransferases, the number of transfers, including interdomain, and lack of resolution in the phylogenies complicate any inference beyond the ancestors of specific Euryarchaeota clades (Supplementary Figures 47-49) but they were probably not found in the LMA. The combined phylogenies of Mcr, Mtr, and markers associated with them suggest that both complexes were inherited vertically by the various Proteoarchaeota lineages, including Korarchaeota and Verstraetearchaeota. In the case of Verstraetearchaeota, this vertical inheritance includes Ehb (Figure 3b). The phylogenies of the WLP components are more poorly resolved (Supplementary Figures 37-41) but in general the Verstraetearchaeota have not acquired these genes through recent transfers. Similarly, information from Mcr, Ehb, m16, and previous work^10^, indicates that the ancestor of Acherontia employed methanogenesis coupled to the WLP.

Combined with the phylogenies of methanol and methylamine methyltransferases, these observations imply that often hydrogen-dependent methylotrophic methanogenesis is a recent emergence due to a loss of the WLP. Other occurrences (Methanonatronarchaeales, Methanomassiliicoccales) could be the result of ancient transfer events. In that view, Ehb in Acherontia and many Verstraetearchaeota is actually a WLP-H_2_/CO_2_ methanogenesis remnant. Topological differences among the phylogenies of subsystems (WLP, Mcr, methyltransferases) indicate that methylotrophic methanogenesis was assembled as a patchwork of transfers. Similar outlier cases exist in the inheritance of H_2_/CO_2_ methanogenesis, too. The Archaeoglobi member *Ca.* Methanomixophus hydrogenotrophicum^9^ possesses an apparently vertically inherited Mtr. However, its Mcr and some of the associated methanogenesis markers are more recent acquisitions from within the Proteoarchaeota (Figure 1). It is uncertain whether that constitutes a homologous recombination or if ancestrally Methanomixophus behaved like Hecatella.

To determine whether the methanogenic ancestor had the capacity for alkane oxidation and further consolidate our predicted phenotype, we reconstructed ancestral sequences for Mcr, Mtr, Eha, and Ehb, for various possible roots (Supplementary Data 15). To account for bias introduced by taxa with missing subunits, we also reconstructed the supermatrix phylogenies and ancestral sequences only using taxa possessing all subunits of the respective complexes. Root placement does affect the reconstructed sequences and by extension their highest similarities but in general these consist of H_2_/CO_2_ and more rarely methylotrophic methanogens (Methanobacteriales, Methanococcales, Verstraetearchaeota in Ehb, some Methanosarcinales) but no alkane oxidizers. For McrA, we also performed homology modeling of the ancestral sequences. Both in terms of sequence conservation^10^ and upon a cursory comparison of the methyl-CoM binding cavity size between ancestral and extant sequences, the ancestral McrA did not have the capacity to accommodate larger alkyl-CoM molecules (Supplementary Figure 50). Thus, it is unlikely that the LMA had any capacity for alkanotrophy, even if the Mcr-like homolog was a basal divergence.

To summarize, the ancestor of non-DPANN Archaea and perhaps all Archaea was a hydrogenotrophic, carbon fixing methanogen that could use CO_2_ and maybe methanol as substrates but not oxidize alkanes. However, the loss of this metabolism was far from a straightforward process, creating varying degrees of intermediate metabolic states present across extant Archaea. These states are centered around the WLP and result mainly in various forms of mixotrophy. The lineages that possess them, such as Mnemosynellales, Hecatellales, and Thorarchaeota thus occupy diverse niches in anaerobic carbon cycling. Hydrogen-dependent methylotrophic methanogenesis has arisen from H_2_/CO_2_ methanogenesis multiple times in unrelated recent clades due to losses of the WLP and through patchwork acquisitions of other components. The anaplerotic H_2_/CO_2_ hydrogenases Eha and Ehb are prime examples of remnants that survive these metabolic transitions but the evolutionary pressures that have shaped the emergence of these large complexes warrant further study.

## Methods

### DUF distribution

We determined the taxonomic distribution of 4049 DUFs and Uncharacterized Protein Families (UPFs) from Pfam release 32.0 with a custom script (distributions_uniprot.py) against a local copy of Uniprot (release 2019_07). For families where no distribution was found, due to lack of cross-references to Pfam, we estimated the distribution from that family’s “Species” tab. Families with at least 50% Archaea in their distribution were retained for downstream analyses as “archaeal” DUFs.

### Homology searches

For initial homology searches we used HMMER 3.2.1^47^ with a cutoff of 1E-5 against local databases of 1808 archaeal and 25118 bacterial genomes. These genomes consist of all Archaea and Bacteria entries on NCBI as of 2019.06.01 dereplicated at species level. The HMM profiles were retrieved preferably from Pfam^48^ or, if one could not be retrieved, from eggNOG’s arCOGs^49^. For the 155 DUFs and genes from^23^, we also searched against local databases of 1611 Eukaryotes and 14494 viruses with the same parameters. Due to only getting hits of dubious quality, all eukaryotic sequences were ultimately removed.

For searches that produced too many hits (as a rule of thumb >1000), we performed a new homology search using DIAMOND^50^ v0.9.24.125 (blastp -e 1e-5 --more-sensitive -k 1000) with a single seed sequence.

### Alignment and single gene phylogenies

We aligned all datasets with MUSCLE^51^. Then we manually curated the alignments to remove distant and/or poorly aligning homologs and fuse contiguous fragmented sequences with a custom script (fuse_sequences.py) and realigned them. Finally, we trimmed the alignments with BMGE^52^ (BLOSUM30).

We reconstructed all single gene phylogenies in IQ-Tree 2^53^ under the model automatically selected by Modelfinder^54^ (-m MFP). We calculated branch supports with 1000 ultrafast bootstrap^55^ and 1000 aLRT SH-like^56^ replicates, and the approximate Bayes test^57^ (-bb 1000 -alrt 1000 -abayes). We visualized all phylogenies in iTOL^58^.

### Mcr, Mtr, Eha, Ehb, and Hcg supermatrix phylogenies

To increase the signal of Mcr, Mtr, Eha, Ehb, and Hcg sequences, we constructed a series of supermatrix phylogenies with taxa that possessed at least two proteins of the respective complex/pathway. Specifically, we concatenated McrABG (McrCD were among the 38 methanogenesis markers and generally not used in the literature), MtrABCDEFG (MtrH was problematic for reasons detailed above), and EhbABCDEFGHIJKLMNOP. For Eha we included subunits EhaBCDEFGHJLMNO. In the Hcg genes we noticed strongly supported incongruences already in the single gene trees and reflected in gene co-localization (Supplementary Figure 35, Supplementary Data 8, 9), so we created two supermatrices; HcgAEFG and HcgBC.

We inferred single gene Maximum Likelihood (ML) phylogenies in IQ-Tree 2 with the trimmed alignments (as above) of the proteins in each supermatrix under the model predicted by Modelfinder with 100 bootstrap replicates (-b 100). We collapsed nodes with support below 80% with TreeCollapseCL 4 (http://emmahodcroft.com/TreeCollapseCL.html). We tested these trees for congruence against the supermatrix tree using the internode certainty test^59^ in RaxML^60^. We removed any incongruent sequences from their respective subunits and repeated the process until no further incongruence could be detected. The only exception was the Methanococcales+Methanobacteriales clade of Mcr where despite our best efforts we could not detect the source of incongruence and ultimately disregarded it, as we did not consider it to affect the overall topology.

For Ehb, even though they did not qualify as (strongly supported) incongruences, the position of Methanococcales was inconsistent among subunits and their synteny was far less conserved than other clades. Thus, to explore potential homologous recombination events, we constructed additional phylogenies for subsets of the Ehb subunits (EhbEFGHIKLMO, EhbEGHIKLM). Detailed explanations for the rationale behind the subunit choices for the concatenations of Eha, Ehb, and Hcg are given in the Supplementary Methods.

For the final concatenated datasets, we ran phylogenies in IQ-Tree 2 under the same parameters as single gene trees above, then used these as guide trees to infer phylogenies under the LG+C60+F+G model with the PMSF approximation^61^. Branch supports were calculated with 1000 ultrafast bootstrap and 1000 aLRT SH-like replicates, and the approximate Bayes test.

For all synteny comparisons in the manuscript figures we used GeneSpy^62^.

### Targeted reconstruction of genomes from the Juan de Fuca Ridge and Shengli metagenomes

We retrieved publicly available reads of metagenomes that contained the target organisms from division NRA7 and Bathyarchaeota (assembly accessions; JdFR-20: GCA_002011155, JdFR-21: GCA_002011165, JdFR-10: GCA_002009985, JdFR-11: GCA_002011035, MAG-15: GCA_014361185, MAG-16: GCA_014361165) were from SRA (JdFR: SRR3723048, SRR3732688; Shengli: SRR11866725, SRR11866724, SRR11866717) and quality filtered them using BBDuk (https://sourceforge.net/projects/bbtools/) and Sickle^63^. We assembled the reads using metaSPADES v3.14.1^64^. The JdFR metagenomes were assembled individually, while the Shengli metagenomes were co-assembled, as in the original publication^32^. Both JdFR and Shengli metagenomes were then processed identically using the uBin helper scripts^65^. Automated binning was performed using ABAWACA^66^ with 3000/5000 and 5000/10000 as minimum/maximum scaffold size parameters, respectively. Additional automated binning was performed with MaxBin2^67^ and both available marker sets encompassing 40 or 107 marker genes, respectively, were employed. The resulting four sets of bins were consolidated in DASTool^68^. Target bins were picked through each organism’s rpS3 sequence in Genbank and then curated in uBin using GC, coverage and taxonomy^65^, supervised by 38 universal archaeal marker genes^69^. Since they possessed at least one marker, we also produced genomes of two Geothermarchaeota (JdFR-13: GCA_002011075, JdFR-14: GCA_002011085) and three Hydrothermarchaeota (JdFR-16: GCA_002010065, JdFR-17: GCA_002011115, JdFR-18/Ca. Hydrothermarchaeum profundi: GCA_002011125) in the same manner. Genome quality was estimated with CheckM^70^ and we manually picked one genome for each species based on it. All our bins were improvements on the ones already submitted to NCBI, except JdFR-13 that contained more contigs/scaffolds but a higher N50.

### NRA7 and Bathyarchaeota taxonomy and phylogenomics

As per their GTDB^3^ classification, the three Mnemosynella species (*Ca.* M. biddleae/JdFR-20,21, *Ca.* M. sp./MAG-15, *Ca.* M. hypogeia/MAG-16) and *Ca.* H. orcuttiae (JdFR-10,11) are members of order-level lineages in Archaeoglobi and Bathyarchaeia respectively. Due to their higher quality and inclusion in our local genomic databases after the dereplication, we refer to JdFR-21 and JdFR-11 throughout this text.

For their phylogenomic placement we used 36 Phylosift^71^ markers (DNGNGWU00035: porphobilinogen deaminase, was omitted, since it yielded too few hits at our default 1E-5 HMM search cutoff). We performed the homology searches, alignments, and dataset curation as described above. We added sequences of the Mnemosynella species to a set of 183 taxa covering the taxonomic range of Archaea and *Ca*. H. orcuttiae to the Bathyarchaeia representative genomes in GTDB r95. The 183 archaeal taxa included genomes from Hydrothermarchaeota and Geothermarchaeota binned here substituting their NCBI counterparts. We downloaded the representative genomes for Bathyarchaeia and Archaeoglobi from NCBI as nucleotide contigs (.fna files) and determined open reading frames for all genomes with Prokka^72^ (--kingdom Archaea --compliant), omitting JdFR-11 and JdFR-21. As an outgroup for the Bathyarchaeia phylogeny, we used the Nitrososphaeria from the set of 183 archaeal genomes, except for the Brockarchaeota^73^ whose position was unstable in this case.

In both cases, we used IQ-Tree 2 to reconstruct the following phylogenies:

1. Model automatically selected by Modelfinder (-m MFP)
2. LG+C60+F+G (PMSF approximation with (1) as the guide tree)
3. Two phylogenies for each supermatrix under the GHOST heterotachy model^74^. For the Bathyarchaeia dataset in the first phylogeny the number of categories was determined in Modelfinder (-mset LG -mrate E,H) and in the second phylogeny the maximum number of categories was set to three (-mset LG -mrate E,H -cmax 3). This corresponds to the highest number of categories for which the number of positions in the supermatrix approached or was >10x the number of free parameters to be estimated in the model. For the complete archaeal dataset, Modelfinder crashed upon reaching H4, so the respective datasets were set to H3 and H2.
4. GTR4 with SR4-recoded^75^ data (-mset GTR -mfreq F,FQ)
5. SR4C60 as in^28^ (PMSF approximation with (4) as the guide tree)
6. GTR6 with Dayhoff6-recoded^75^ data (-mset GTR -mfreq F,FQ)
7. A series of phylogenies with progressively desaturated subsets of the original supermatrix, under the model automatically selected by Modelfinder (-m MFP). The empirical Bayesian site-specific rates were calculated from the supermatrix and phylogeny in (2) with fixed branch lengths (-blfix) under the Poisson+G16 model. The Poisson (JC-like) model was selected based on the literature^76,77^ and, after a small internal benchmark (Supplementary Data 5). For the benchmark we estimated both empirical Bayesian (“random effects”) and ML (“fixed effects”) rates for the supermatrices under the Poisson, Poisson+G16, LG, LG+G16 models. Then we calculated Pearson and Spearman correlations between (i) rate estimation methods with a given model, (ii) substitution matrices, (iii) with and without rate heterogeneity. All correlations were very strong, but the Poisson model was slightly more internally consistent. G16 was chosen to imitate the behavior of Rate4Site^78^ and because +R16 in IQ-Tree does not function together with -blfix.
8. LG+C60+F+G for the progressive desaturation datasets (PMSF approximation with the respective phylogenies from (7) as guide trees)

For all runs, branch supports were calculated with 1000 ultrafast bootstrap and 1000 aLRT SH-like replicates, and the approximate Bayes test.

To corroborate the taxonomic level of the NRA7/Mnemosynellales (GTDB order JdFR-21) and the Bathyarchaota Subgroups 20&22/Hecatellales (GTDB order B25) clades, we calculated pairwise ANI and AAI values for all GTDB representative genomes in Archaeoglobi and Bathyarchaeia. JdFR-21 and JdFR-11 were substituted with the MAGs binned here. ANI values were calculated with orthoANI^79^ and AAI with CompareM (https://github.com/dparks1134/CompareM).

To assess the biogeographic and environmental distribution of Mnemosynellales and Hecatellales, we constructed their 16S phylogenies. For Mnemosynellales we used all sequences in SILVA classified under JdFR-20 (SILVA, SSU r138.1). We picked Hecatellales sequences from among Bathyarchaeia sequences (SILVA Ref NR, SSU r138.1; SILVA contained >50k sequences), aligned with MUSCLE and through a preliminary BioNJ phylogeny^80^. We used the 16S sequences from *Ca.* Polytropus marifundus (GCA_002010305) and RBG-16-48-13 (GCA_001775995) as outgroups for Mnemosynellales and Hecatellales respectively. We aligned the final datasets with MAFFT L-INS-i v7.475^81^, curated them manually, trimmed with BMGE (PAM100), and reconstructed an ML phylogeny with IQ-Tree 2 as above.

### Metabolic reconstructions

The metabolic potential of the *Ca.* M. biddleae, *Ca.* M. hypogeia, and *Ca.* H. orcuttiae was predicted with BlastKOALA^82^ using the JdFR-21 and JdFR-11 taxids from NCBI and searching against the species_prokaryotes database. Additional annotations were produced with HydDB^83^ (including supplementary annotation of Fe-Fe hydrogenases using downstream genes), dbCAN2^84^ (dbCAN meta server with all options enabled), and MEROPS^85^ (searched locally with DIAMOND blastp, cutoff 1E-5).

### Outgroup-free rooting and rootstraps

For all the Mcr, Mtr, Eha, Ehb, and Hcg supermatrices described above, we performed non-outgroup rooting with the MAD^86^ and MinVar^87^ methods on phylogenies under the LG+C60+F+G model and 100 bootstrap replicates (-b 100) (PMSF approximation as above). Rooted phylogenies were also inferred under the NONREV non-reversible protein model^53^ with 100 bootstrap replicates. The sets of rooted phylogenies and bootstrap trees were used to calculate rootstrap supports^88^. We also rooted the singlegene methyltransferase (MtaB, MtmB, MtbB, MttB) phylogenies with MAD and MinVar.

### Gene and site concordance factors (gCF, sCF)

We calculated gCF and sCF^89^ for Eha, Ehb, Mtr, and Mcr using the mixture model phylogenies as species trees, the subunit single gene phylogenies (with incongruences resolved) as gene trees, and the supermatrices as input alignments. To isolate the effect of individual subunits on the signal, we also calculated sCF with the mixture model phylogenies as species trees but in a series of separate runs with each subunit as the input alignment.

### Site rate estimation and transmembrane segment prediction

We estimated empirical Bayesian and ML site-specific rates for all Eha, Ehb, Mcr, and Mtr subunits as above, from their trimmed alignments (before congruence testing) and respective single gene phylogenies. We benchmarked the effect of model choice on such shorter alignments by calculating Pearson and Spearman correlation coefficients between ML and empirical Bayesian rates under Poisson and Poisson+G16 separately and between ML rates under the two models. While ranks did not change, short alignments created unrealistic outlier values in ML rates when rate heterogeneity was included in the model. However, the Bayesian Poisson+G16 and ML Poisson rates were almost perfectly correlated. We tested whether any subunits within each complex had significantly higher or lower rates through a Kruskal-Wallis test followed by Dunn’s test and a series of Mann-Whitney U tests for all subunit pairs of each complex.

We calculated the transmembrane per site probability for each subunit both numerically with the Python implementation (https://github.com/dansondergaard/tmhmm.py) of TMHMM2.0^90^ and on the Polyphobius server^91^, and as a structural feature on the TOPCONS2 server^92^ and with DeepTMHMM (https://biolib.com/DTU/DeepTMHMM), to account uncertainties, due to differences among algorithms and the fact that we used *Methanothermobacter marburgensis* sequences from the trimmed alignments as input. For all subunits with predicted transmembrane segments in TOPCONS2, we calculated Spearman and Pearson correlations between empirical Bayesian rates under Poisson+G16 and the transmembrane helix probability from TMHMM2.0 and Polyphobius. We also ran the Mann-Whitney test to compare the populations of rates between positions that were predicted as transmembrane helices and those that were not (i.e. extramembrane) in TOPCONS2 and DeepTMHMM.

### Ancestral sequence reconstruction

We reconstructed ancestral sequences via the empirical Bayesian method in IQ-Tree 2 (-asr) for all nodes and all concatenated subunits of Eha, Ehb, Mcr, and Mtr in two ways. First, we used the supermatrix phylogenies constructed previously for each complex under the LG+C60+F+G model but substituted the concatenation of trimmed alignments with their untrimmed equivalents for the reconstruction. We parsed the ASR output with a custom script (ASR_parser.py) that separates the sequences of individual subunits and calculates the mean posterior probability for the reconstructed sequence of each node. These reconstructed sequences consist of the residue with the highest probability for each site. The mean posterior probabilities are gross underestimates, since IQ-Tree does not reconstruct indels, and thus the probability for sites with many gaps ends up being very low. Our second approach to ASR was almost identical. However, this time we reduced the datasets for each complex to only include taxa that possessed a complete complex to avoid including large gaps in the supermatrix that could affect the reconstruction. If a subunit was missing in entire clades of the phylogeny, we either omitted that subunit (EhaL, EhbKLN) or these taxa in the case of Methanopyrales in Mcr where we had only three subunits. We then inferred phylogenies with automatic model selection (-m MFP) and used them as guide trees for LG+C60+F+G phylogenies (PMSF approximation), reconstructing ancestral sequences in tandem.

Finally, we retroactively added indels to the reconstructed sequences by a consensus-like approach. For each subunit, the reconstructed sequences corresponding to potential LMA nodes from both approaches were added to their respective datasets of complete complex taxa. These were realigned and trimmed with Clipkit^93^ (-m gappy - g 0.5) to remove positions with at least 50% gaps. Due to their missing clades, EhaL and EhbKLN were omitted from indel inference.

### McrA homology modeling

We performed all homology modeling on the Phyre2 server^94^ with the intensive mode. For all visualization and structural alignments, we used Pymol v2.4^95^ and its alignment plugin, aligning each homology model to the best template picked by Phyre2 (all to one, defaults). All RMSDs were <0.4 Å.

### Statistical analyses

We performed all statistical tests in base R^96^, except Dunn’s test for which we used the dunn.test package^97^. We visualized results using base R or ggplot2^98^.

## Data availability

Custom scripts mentioned in the Methods section can be found in the GitHub repository: https://github.com/ProbstLab/Adam_Kolyfetis_2021_methanogenesis.git All Supplementary Data files have been uploaded to Figshare under https://doi.org/10.6084/m9.figshare.15088110.v1.

## Author contributions

Roles defined according to the CRediT system. For each role, name order corresponds to size of contribution. Brackets denote equal contribution in the author list order.

Conceptualization: PSA; Data curation: GEK, PSA, (TLVB, AJP); Formal analysis: PSA, GEK, (TLVB, AJP); Funding acquisition: (PSA, AJP); Investigation: GEK, PSA, (TLVB, AJP); Methodology: PSA, AJP; Project administration: PSA; Resources: AJP, CEV; Supervision: PSA, AJP, CEV; Software: GEK, PSA, TLVB; Validation: (PSA, GEK); Visualization: GEK, PSA, TLVB; Writing-original draft: PSA, GEK; Writing-reviewing & editing: (PSA, GEK, TLVB, CEV, AJP).

The authors have agreed that PSA and GEK contributed equally to the manuscript and both may put their name first in the author order for the purposes of including this article in their CV publication list.

## Acknowledgments

PSA is supported by a postdoctoral fellowship from the Alexander von Humboldt Foundation. TLVB and AJP are supported by funding from the Ministerium für Kultur und Wissenschaft des Landes Nordrhein Westfalen (“Nachwuchsgruppe Dr. Alexander Probst”). The authors would like to thank (1) Michael Rappé, Sean Jungbluth, Bo-Zhong Mu, and Yi-Fan Liu for permissions to use and assistance with their metagenomic data; (2) Jennifer Biddle and Beth Orcutt for allowing us to name species after them; (3) Suha Nasser-Khdour, Robert Lanfear, and Bui Quang Minh for helpful discussions and advice on many of the analyses involving IQ-Tree; (4) Aharon Oren for advice and corrections regarding microbial nomenclature.

