## Supplementary Information for "Genomic remnants of ancestral hydrogen and methane metabolism in Archaea drive anaerobic carbon cycling"

### Descriptions of proposed taxa

Description of *Candidatus* Mnemosynella biddleae (gen. nov., sp. nov.)

*Candidatus* Mnemosynella biddleae (Mne.mo.sy.nel'la. N.L. fem. dim. n. *Mnemosynella*, little Mnemosyne, after the Greek mythology Titaness, goddess of memory, and mother of the Muses; a reference to the species containing multiple methanogenesis markers and "remembering" its methanogenic ancestry; bidd'le.ae. N.L. gen. fem. n. *biddleae*, named after Jennifer Biddle, in honor of her contributions to microbial ecology).

Description of *Candidatus* Mnemosynella hypogeia (gen. nov., sp. nov.)

*Candidatus* Mnemosynella hypogeia (Description as above for the genus Mnemosynella. hy.po.gei'a. N.L. fem. adj. *hypogeia*, subterraneous, earth-born )

Description of Mnemosynellaceae (fam. nov.)

(Mne.mo.sy.nel.la.ce'ae. N.L. fem. n. *Mnemosynella*, a Candidatus genus name; -*aceae*, ending to denote a family; N.L. fem. pl. n. *Mnemosynellaceae*, the *Mnemosynella* family)

Description of Mnemosynellales (ord. nov.)

(Mne.mo.sy.nel.la'les. N.L. fem. n. *Mnemosynella*, a Candidatus genus name; -*ales*, ending to denote a family; N.L. fem. pl. n. *Mnemosynellales*, the *Mnemosynella* order)

Description of *Candidatus* Hecatella orcuttiae (gen. nov., sp. nov.)

*Candidatus* Hecatella orcuttiae (He.ca.tel'la. N.L. fem. dim. n. *Hecatella*, little Hecate, after the Greek mythology goddess of witchcraft and crossroads; a reference to the species' metabolism being at the "crossroads" of methanogenesis and the Wood-Ljungdahl pathway by means of its Mtr complex; or.cut'ti.ae. N.L. gen. fem. n. *orcuttiae*, named after Beth Orcutt, in honor of her contributions to microbial ecology, including studies on the Juan de Fuca Ridge).

Description of Hecatellaceae (fam. nov.)

(He.ca.tel.la.ce'ae. N.L. fem. n. *Hecatella*, a Candidatus genus name; -*aceae*, ending to denote a family; N.L. fem. pl. n. *Hecatellaceae*, the *Hecatella* family)

Description of Hecatellales (ord. nov.)

(He.ca.tel.la'les. N.L. fem. n. *Hecatella*, a Candidatus genus name; -*ales*, ending to denote a family; N.L. fem. pl. n. *Hecatellales*, the *Hecatella* order)

Description of *Candidatus* Geothermarchaeum rappei (gen. nov., sp. nov.)

*Candidatus* Geothermarchaeum rappei (Ge.o.therm.ar.chae'um. Gr. fem. n. *gê*, the earth; Gr. masc. adj. *thermos*, hot; N.L. neut. n. *archaeum*, ancient one, archaeon, from Gr. masc. adj. *archaîos*, ancient; N.L. neut. n. *Geothermarchaeum*, an archaeon from hot earth; rap.pe'i. N.L. gen. masc. n. *rappei*, named after Michael Rappé, in honor of his contributions to microbial ecology, including obtaining the original samples from the Juan de Fuca Ridge used in this study).

Description of *Candidatus* Scotarchaeum ottlingeri (gen. nov., sp. nov.)

*Candidatus* Scotarchaeum ottlingerii (Scot.ar.chae'um. Gr. masc. n. *skotos*, darkness; *archaeum*, ancient one, archaeon, from Gr. masc. adj. *archaîos*, ancient; N.L. neut. n. *Scotarchaeum*, an archaeon living in darkness; ott.lin'ge.ri. N.L. gen. masc. n. *ottlingeri*, named after Markus Ottlinger, German visual artist depicting hydrothermal settings and biofilms).

Description of Geothermarchaeaceae (fam. nov.)

(Ge.o.therm.ar.chae.a.ce'ae. N.L. neut. n. *Geothermarchaeum*, a Candidatus genus name; -*aceae*, ending to denote a family; N.L. fem. pl. n. *Geothermarchaeaceae*, the *Geothermarchaeum* family)

Description of Geothermarchaeales (ord. nov.)

(Ge.o.therm.ar.chae.a'les. N.L. fem. n. *Geothermarchaeum*, a Candidatus genus name; -*ales*, ending to denote a family; N.L. fem. pl. n. *Geothermarchaeales*, the *Geothermarchaeum* order)

Description of *Candidatus* Pyrohabitans jungbluthii (gen. nov., sp. nov.)

*Candidatus* Pyrohabitans jungbluthii (Py.ro.ha'bi.tans. Gr. neut. n. *pŷr*, fire; L. pres. part. *habitans*, inhabiting; N.L. masc. n. *Pyrohabitans*, an inhabitant of fire; jung.blu'thi.i. N.L. gen. masc. n. *jungbluthii*, named after Sean Jungbluth, in honor of his contributions to microbial ecology, including obtaining the original samples from the Juan de Fuca Ridge used in this study).

### Supplementary Methods

#### Detailed description of homology search methods for Eha, Ehb, Hcg

Since several of the proteins in the Eha, Ehb, and Hcg sets were either DUFs or part of the lists from Gao & Gupta[1], we could compare the single gene distributions and phylogenies to determine potential issues in the homology searches for the more problematic (poorly annotated, fast evolving etc.) proteins in each complex or pathway. HMM searches often produced >1000 hits for most subunits and thus a lot of computational power, time, and manual work would be required to isolate our homologs of interest. For that reason, we instead searched for homologs using DIAMOND blastp with two seeds on the taxonomic extremes of each complex or pathway. For Eha they were *Methanothermobacter marburgensis* (Methanobacteriales) and *Methanolacinia petrolearia* (Methanomicrobiales), for Ehb *M. marburgensis* and *Ca.* Methanosuratus petracarbonis (Verstraetearchaeota), and for Hcg *M. marburgensis* and *Desulfurobacterium thermolithotrophum* (Desulfurobacteriales). *M. marburgensis* was used as a reference for picking the first seed using information from [2]. Other known homologs from the literature in *Methanococcus maripaludis, Methanocaldococcus jannaschii, Methanothermobacter thermoautotrophicus, Methanopyrus kandleri* were used as (additional) seeds in some cases. When obtaining fewer homologs than expected (i.e. entire clades missing partially or entirely), the HMM profiles or DIAMOND seeds were expanded by using all hits of the previous search round to create a new HMM profile or collection of seeds and rerun the search. To clean up the datasets and retain our homologs of interest, we first aligned the initial pool of hits with MUSCLE[3]. Then in Seaview[4] we manually inspected the alignments, constructed preliminary phylogenies with BioNJ[5] (Poisson or Observed distances) and/or PhyML[6] (default options, no topology optimization), and isolated (monophyletic) clades containing our homologs of interest. None of these initial phylogenies were retained, since they were made on a trial and error basis, but they can be reproduced through the datasets in the Supplementary Data. We confirmed suspected gene losses by manually comparing the hits of the homology searches with the synteny in each taxon. Below we list for each gene whether we used the base seeds for a DIAMOND search and if it was found in this study as a DUF, or in Gao & Gupta[1] with the corresponding Pfam or arCOG accession for the HMM profile. We also detail any deviations in the homology search methodology.

##### Eha

EhaA: (Gao & Gupta, 2007), PF17367, custom HMM from all the hits of the original

search
EhaB: (Gao & Gupta, 2007), arCOG04828
EhaC: DUF2109
EhaD: DUF2108
EhaE: DUF2107
EhaF: DUF2106
EhaG: DUF2105
EhaH: PF10125, found from synteny
EhaI: (Gao & Gupta, 2007) contained only arCOG05034. From Uniprot cross-references for multiple sequences, we found HMM profiles for arCOG05034 and arCOG06464 and pooled the hits together. Mnemosynellales, Persephonarchaea, and *M. kandleri* were not found through HMM searches, so we added their sequences manually from expected synteny and BLAST searches against NCBI. We pooled all these sequences for recursive DIAMOND searches, using the hits from the previous search as seeds. We stopped after 3 rounds of searches, since the number of hits decreased afterwards.
EhaJ: base seeds
EhaK: (Gao & Gupta, 2007) contained only arCOG08277. From Uniprot cross-refs, found HMM profiles for arCOG06676, arCOG08277, arCOG10247, arCOG60928, and proceeded as with EhaI. Only 1 round of DIAMOND searches was necessary, as we found no further hits afterwards.
EhaL: DUF2104, custom HMM from all the hits of the original search.
EhaM: DUF1959, custom HMM from all the hits of the original search.
EhaN: base seeds
EhaO: base seeds
EhaP & EhaQ: In *M. kandleri*, the gene called EhaP in Uniprot (AAM01675.1) is an unrelated ferredoxin, distantly related to RnfB homologs in Firmicutes but also to its own EhaR* (AAM01674.1). The real EhaP homolog in *M. kandleri* is called EhaQ (AAM01677.1). However, when running BLASTp against NCBI non-redundant with the *M. marburgensis* EhaQ, the first *M. kandleri* hit at twilight zone homology is AAM01676.1 listed in Uniprot as EhbK (marked with an asterisk in Figure 3a). Using *M. maripaludis* as the query, the first *M. kandleri* hit is the ferredoxin AAM01674.1. We did not find either of these sequences in our local DIAMOND searches, even if they could be considered as EhaQ-like. Many Methanococcales also carry an EhaQ-like homolog that was not picked up in our homology searches but is part of the Eha cluster (AAB98510.1 in *M. jannaschii*, marked with an asterisk in Figure 3a).
EhaP: base seeds, plus MSBL1 archaeon SCGC-AAA382A20
EhaQ: *M. marburgensis* and *M. maripaludis*
EhaR: The gene called EhaR in *M. marburgensis* (ADL58396.1) was annotated as a ribokinase and only picked hits in Methanobacteriales. The one called EhaR in *M. maripaludis* based on Uniprot and synteny (ABO34430.1) and its closest homolog in *M. kandleri* (AAM01674.1) did have some hits in Methanobacteriales. These included SCG85576.1 in *Methanobacterium congolense*, and when NCBI non-redundant was queried, ADL59095.1 (MvhB) in *M. marburgensis*. However, the syntenic polyferredoxin ADL58399.1, as well as SCG85576.1, and ADL59095.1, when used in a BLASTp search in the same way, pick up AAM01674.1 as their first hit in *M. kandleri*. The final seeds for EhaR were *M. maripaludis* (ABO34430.1), *M. kandleri* (AAM01674.1), and *M. marburgensis* (ADL58399.1).
EhaS: base seeds, DIAMOND with all hits of the first search.
EhaT: base seeds, DIAMOND with all hits of the first search

Ehb

EhbA: base seeds
EhbB: arCOG04878
EhbC: (Gao & Gupta, 2007), arCOG04877
EhbD: base seeds
EhbE: base seeds
EhbF: base seeds
EhbG: (Gao & Gupta, 2007), arCOG05076
EhbH: base seeds
EhbI: base seeds
EhbJ: custom HMM profile from pooled hits of arCOG06490 and arCOG06683
EhbK: base seeds, plus *M. maripaludis*
EhbL: base seeds
EhbM: base seeds
EhbN: base seeds
EhbO: base seeds
EhbP: (Gao & Gupta, 2007), PF10622

Hcg

HcgA: base seeds
HcgB: DUF3236
HcgC: DUF1188
HcgD: absent in *D. thermolithotrophum,* seeds were *M. marburgensis* and *M. kandleri*
HcgE: (Gao & Gupta, 2007), arCOG01677
HcgF: UPF0254
HcgG: (Gao & Gupta, 2007), PF10113

Subunit omissions from Eha, Ehb, and Hcg supermatrices

As mentioned in the Methods section, certain subunits were omitted from the supermatrices of Eha, Ehb, and Hcg. The specific reasons for each subunit are summarized below.

EhaA: Missing in Methanomicrobiales; the phylogeny does not even recover the monophyly of the other clades (Persephonarchaea and Methanopyrales inside Methanobacteriales at different branches, Mnemosynellales inside Methanococcales).
EhaI: Issues in the homology searches. Very short gene with strange topology in its phylogeny (Persephonarchaea and Methanococcales inside Methanobacteriales).
EhaK: Issues in the homology searches. Very short gene with strange topology in its phylogeny (Methanopyrales inside Methanobacteriales, Methanococcales as grades at the base of Methanomada, Persephonarchaea and Mnemosynellales as grades at the base of Methanomicrobiales).
EhaP: Ferredoxin with very low conservation among clades and poorly aligning. Issues with the annotations and homology searches (i.e. establishing orthology).
EhaQ: Ferredoxin with very limited distribution (Methanobacteriales, Methanococcales). Issues with the annotations and homology searches (i.e. establishing orthology).
EhaR: Limited distribution (Methanobacteriales, Methanococcales, Methanopyrales). Issues with the annotations and homology searches (i.e. establishing orthology).
EhaS: Paralog of Ftr whose distribution does not match the other subunits[7]. Only part of the Eha cluster in Methanobacteriales.
EhaT: Very limited distribution (only Methanobacteriales).

For EhaAIK their short length makes their omission probably inconsequential. In general, we did not wish to risk the overall supermatrix signal by introducing multiple poorly conserved subunits or with dubious orthology. Similar to EhaA, EhaC also has a slightly problematic topology but is not as pronounced (Methanococcales and Methanopyrales respectively monophyletic but inside Methanobacteriales).

EhbJ: Only omitted in the Methanococcales cluster supermatrices. Strange topology of the single gene phylogeny (Methanofastidiosales and Theionarchaea at two separate branches within Methanobacteriales). A similar monophyly issue exists in EhbH (either Verstraetearchaeota or Methanococcales+Acherontia within Methanobacteriales, depending on the root position, but monophyletic among themselves) but it is less pronounced in our opinion, so that subunit was kept in the supermatrices.

HcgD: Absent in Desulfurobacteriales; there was no monophyletic clade for the other taxa in the preliminary phylogenies.

### Supplementary Results & Discussion

#### Functional annotation and evolution of the methanogenesis markers

m4-m9: A cluster of six genes that are co-localized in many genomes and are predicted to be co-transcribed in *Methanolobus psychrophilus* R15[8] but do not co-purify with Mcr[9]. Some of them are absent from various alkane oxidizers but it is impossible to determine whether this is a result of genome incompleteness or variation in the processes controlled by these genes. There exist very few cases of putative remnants among them, for example the hot spring Thaumarchaeota SAG E04 (GCA_000405745) that contains genes of the WLP[10]. M9 is a homolog of McrC.

m10: AtwA (also called McrA2) is an ATP-binding protein[11] that is necessary for Mcr activation[9]. We found that there exist two homologs that we call here canonical AtwA and AtwA-like, assuming that the canonical AtwA has a wider taxonomic distribution. The presence of multiple homologs (AtwA1, AtwA2) can be found in the annotations of publicly available sequences but these do not correspond to the two clades we recovered i.e. there are both so-called AtwA1 and AtwA2 sequences within both canonical AtwA and AtwA-like. AtwA-like is absent in most Halobacterota methane metabolizers. This distribution does not seem to be linked to the presence of multiple Mcr homologs in Methanomada members, as the duplication of the *atwA* gene most probably occurred at the LMA, while the aforementioned Mcr complexes arose from more recent duplications (Figure 1a). The topology of AtwA-like with Methanosarcinales at the base and non-monophyletic Methanomada might have been affected by long branch attraction. However, the canonical AtwA side recovers the monophyly of most lineages as expected from McrABG (Figure 1a) and corroborates the poorly supported NONREV root at Halobacterota. This in turn supports that anaerobic alkane oxidation (AAO) is a more recent invention but Mcr-like complexes still require AtwA.

m11: McrC probably participates in Mcr activation, since it is part of a reductase complex capable of reducing coenzyme $F_{430}$ to the $Ni^{1+}$ form[9] and corresponds to part of the McrA3 component[11]. Other than McrC and m9, we identified a third distant homolog only present in Methanobacteriales (Supplementary Data 2; Supplementary Figure 8 only shows the canonical McrC).

m12-m14: McrD (m12) has been posited to be a chaperone delivering $F_{430}$ from its production after the reaction catalyzed by CfbE to Mcr[12]. However, we think that this association might not be universal, since McrD is present in essentially all methane metabolizers and alkane oxidizers, except for Methanophagales. CfbD (m13) and CfbE (m14) have a different distribution, with one or both being absent in many AAO and even some methanogens. CfbE is encoded by one Methanophagales MAG (GCA_004212135, hot spring). These discrepancies indicate that several Archaea might contain $F_{430}$ biosynthesis variants or even a modified cofactor, such as the methylthio modified $F_{430}$ found in the Mcr in marine Methanophagales[13,14].

m15: MtxX is either a phosphotransacetylase or methyltransferase, part of the MtxXAH operon in *M. barkeri*[15,16]. It is impossible to determine its function from its distribution and phylogeny, since it does not consistently coexist with some other methanogenesis or WLP subsystem but the association with Mtr is probably a rarity.

m16: M16 has sequence similarity to the AIR synthase but its function is unknown. The *M. maripaludis* homolog (MMP0291) is annotated as “hydrogenase expression/formation protein related” and in proximity to some *hyp* genes[17].

m17: DUF1743 includes a series of homologs that contain one of the domains of TiaS, the $tRNA^{Ile}$-agmatinylcytidine synthetase (that homolog was removed from the final phylogeny shown in Supplementary Figure 14) and is very ancient among Archaea, independent of methane metabolism. It is predicted to function as an ATPase[18] but its exact function in methane metabolism or as a remnant remains indeterminate.

m18-m19: No information in the literature, function cannot be predicted.

m20: M20 contains homologs of YcaO that functions as a glycine thioamidase for the post-translation modification of an Mcr glycine (Gly465 in *M. acetivorans*) into thioglycine[19]. The topology of our phylogeny is roughly in agreement with the one published in[19]. This modification is absent in alkane oxidizers.

m21: Rossmann fold protein with a cysteinesulfonic acid modification (PDB ID: 2R47). No further information could be determined.

m22: No useful information.

m23: Annotated as P-type ATPase or HAD hydrolase. Absent in Methanopyrales.

m24: No useful information.

m25: Sparse distribution: absent in Methanopyrales and many Halobacterota lineages but widely present in Lokiarchaeota. Related to dihydrouridine synthase (PF0127). It is co-localized with formylmethanofuran--tetrahydromethanopterin formyltransferase/$F_{420}$-dependent NADP oxidoreductase in Lokiarchaeota and $F_{420}H_2$ oxidase in Mnemosynella, thus a role in energy/hydrogenase-type reactions involving $F_{420}$ is possible.

m26: Based on its distribution, phylogeny, and synteny, m26 is associated with Mtr, even including the Thorarchaeota clade of the canonical MtrA. Its distribution also includes Persephonarchaea sequences (Supplementary Data 2) at twilight zone homology that aligned poorly and ended up on a long branch, thus we removed them. The presence of a divergent m26 in Persephonarchaea suggests that they are former methane metabolizers ($H_2/CO_2$ or methanotrophs) that have already lost all traces of Mtr and are currently also losing its related marker.

m32: UPF0285 is not found outside of Euryarchaeota. Its function cannot be deduced. We recover a strong monophyly of Hydrothermarchaeota (where it is syntenic with Ftr) with Methanomada. This agreement with the species tree (Figure 2a) and synteny supports the case for Hydrothermarchaeota also being former $H_2/CO_2$ methanogens. Absent in Methanopyrales; the *M. kandleri* gene annotated as UPF0285 (MK0078) is a member of the unrelated DUF1464.

m33: MamA catalyzes the methylation of Arg285 (*M. acetivorans*)[20,21]. Its distribution is limited to Methanomada and a subset of Halobacterota. Together with the

distribution and phylogeny of McmA[22] and our phylogeny of m20, it suggests that the *M. acetivorans* pattern of Mcr posttranslational modifications is not universal among methane metabolizers and entirely absent in alkane oxidizers without hampering the complex's functionality.
m34: Limited to Methanomada and a subset of Halobacterota. Its function cannot be deduced.
m35: Limited to Methanomada and a subset of Halobacterota. Its function cannot be deduced. There exists one solved structure (PDB ID: 1KJN) of the *M. thermautotrophicus* homolog that is a selenomethionine-containing homodimer.
m36: Sparse distribution in Methanomada and a subset of Halobacterota. Phylogeny is very poorly resolved.
m37: Its distribution includes Persephonarchaea where it is syntenic with Fwd and Ftr. The synteny could be related to Persephonarchaea being former methane metabolizers (see below about Eha). We cannot infer anything about its function.
m38: Limited to Methanomada, Methanosarcinales, Methanobacteriales. Its function cannot be deduced.

Based on their distributions and phylogenies, many of the 38 methanogenesis markers should probably not be considered as such, instead being "demoted" to partial markers like the various Eha and Ehb subunits.

A few markers (m15, m17, m18, m19, m21, m24, m36) have undergone interdomain horizontal transfers from Archaea to Bacteria, suggesting that they might be incorporated into other metabolic pathways without them necessarily being remnants of methane metabolism.

For some markers whose function is impossible to infer (e.g. m15-m19) we can at least hypothesize a lack of relationship with Mcr, due to their inconsistent presence in alkane oxidizers, while they are found in Archaea and Bacteria often without the Wood-Ljungdahl pathway (WLP).

M15, m16, and m17 each contain a strongly monophyletic Archaeoglobales cluster (including *Ca.* Methanomixophus hydrogenotrophicum) with *Ca.* Methanodesulfokores washburnensis (GCA_004347975, Korarchaeota) inside it. This observation contributed to our inference that Mcr was vertically inherited in Methanodesulfokores and other methylotrophic methanogenesis components were acquired horizontally. Nonetheless, due to the existence of such transfers and the proximity of the Proteoarchaeota Mcr and Mtr sequences to Halobacterota (Figure 1), we have to consider the possibility that Proteoarchaeota acquired both complexes as ancient transfers from within Halobacterota. We do not consider this scenario as likely a scenario further, as it is not parsimonious and clashes with information from other markers (particularly m26 that includes Thorarchaeota).

Evolution of the Hcg proteins

We observed that in Methanococcales the genomic location of the *hcg* genes is not stable. *hcg*BC are clustered together in *M. jannaschii* and distant from *hmd* but *hcgAG* and *hmd* are syntenic (Supplementary Figure 36), corroborating a previous synteny analysis in *M. maripaludis*[23]. This is reflected in the evolutionary histories of the single gene phylogenies and supermatrices where HcgBC and HcgAEFG give different positions for Methanococcales that are further supported by the gene and site concordance factors for the two supermatrices (Supplementary Data 13). For HcgBC, there exists a monophyletic clade of Methanobacteriales and Methanococcales similar to Hmd and Hmd-like. However, in HcgAEFG, Methanobacteriales are monophyletic with Methanopyrales, similar to the archaeal species tree (Figure 2a), albeit not strongly supported. Other than a lack of evolutionary pressure to maintain a tight cluster, these data imply the involvement of homologous recombination events in the evolution of the Hcg pathway and Hmd, barring other sources of bias in the phylogenies. Assuming Hmd and HcgBC represent the correct history, then the most parsimonious scenario involves an ancient lateral acquisition of HcgAEFG by Methanobacteriales or by Methanococcales. The latter case assumes a subsequent dissolution of the cluster in Methanococcales and the lack of resolution in the HcgAEFG phylogeny not conflicting with the Methanopyrales-Desulfurobacteriales monophyly in the HcgBC tree. Nevertheless, if we accept that the branching in Methanomada should reflect the archaeal reference tree, it is HcgBC-Hmd that were transferred horizontally to the ancestral Methanobacteriales or Methanococcales. In any case, an acquisition by the Desulfurobacteriales in a hydrothermal environment is a plausible assumption. Note that in all cases the Methanobacteriales clade includes a recent transfer to Methanomicrobiales as previously noted for Hmd/Hmd-like[7].

For the HcgBC supermatrix, both MAD and MinVar root the tree at the strongly monophyletic group of Methanopyrales-Desulfurobacteriales (with relatively high rootstrap support) while NONREV roots it at the strongly monophyletic group of Methanobacteriales (with low rootstrap support). Neither root agrees with what we would expect from the reference tree i.e. a basal separation of Methanococcales (Supplementary Figure 36b, as displayed). Regarding the HcgAEFG supermatrix, MAD and Minvar disagree in the root placement. MAD roots at Methanopyrales and MinVar at Desulfurobacteriales. The rootstrap values strongly support the latter but both are roughly consistent with what we have observed previously for Hmd/Hmd-like[7]. In any case, this biosynthetic pathway does not appear to predate the ancestor of Methanomada. Finally, as mentioned in the Supplementary Methods, the Desulfurobacteriales do not seem to have a proper HcgD but we did not even recover a proper HcgD monophyletic clade in the preliminary phylogenies either. Thus, it is possible that various paralogs (perhaps with some promiscuity) catalyze the HcgD reaction in different taxa.

Evolution of Eha

Our main interest in the evolution of Eha was the direction of the transfer between Mnemosynellales and Persephonarchaea. With a root between Methanomada and Halobacterota (Figure 3a, as displayed), the Persephonarchaea are not in their expected position (Figure 2a) and have acquired Eha laterally. This scenario is consistent with the high number of methanogenesis markers in Mnemosynellales and the phylogenies of WLP genes from both the carbonyl and methyl branches (CdhB, Mch, Mtd; Supplementary Figures 37-39, Supplementary Data 7). MAD and MinVar outgroup-free rooting placed the root between Methanomicrobiales and the remaining clades reversing the inferred direction of the transfer (Figure 3a, Supplementary Data 12). We recovered this scenario in the phylogenies of the WLP carbonyl methyltransferase module genes (CdhDE) (Supplementary Figures 40, 41, Supplementary Data 7); Eha could have been part of the same transfer event. This inference also suggests that Persephonarchaea could be former methanogens, given that they possess some markers but their position in these phylogenies is unresolved.

A second issue is the second (smaller) Methanobacteriales clade. It contains genomes from a variety of genera and often genomes of partially undefined taxonomy

(Supplementary Data 10). There is no overlap of taxa between the two Methanobacteriales clades, indicating that there were no gene duplications involved but rather ancient homologous recombination events. As with Hcg, we cannot define the original Methanobacteriales clade and by extension a putative source of the transfer although following the species tree (Figure 2a), the smaller Methanopyrales-affiliated clade is the original. Determining the position of the Methanopyrales themselves in reference phylogenies has been long plagued by artifacts[24,25] and it is likely that such issues carry over to the Hcg and Eha phylogenies. The issue is exemplified by the disagreements in the phylogenetic signal among subunits by proxy of their sCF (Supplementary Figure 44a, Supplementary Data 13) where Node 232 (Methanopyrales + small Methanobacteriales) and Node 153 (Methanococcales + large Methanobacteriales) have the lowest values among deeper nodes in the tree. Other low sCF values for Node 151 (Mnemosynellales + Persephonarchaea) could be related to divergence in their sequences and poor signal. Within Methanobacteriales, we observed the absence of Eha in *Methanosphaera*, despite it possessing the WLP[7] and Ehb (this study). Since *Methanosphaera* is a methylotrophic methanogen, it would appear that at least for Methanobacteriales there exists a tight association between $H_2/CO_2$ methanogenesis and Eha that does not persist as a methanogenesis remnant. Judging from its absence in other methanogens (e.g. Nezhaarchaeota, Verstraetearchaeota), Eha is not always indispensable for $H_2/CO_2$ methanogenesis. Depending on whether the Euryarchaeota are monophyletic or not[26] and the direction of the Mnemosynellales-Persephonarchaea transfer, Eha became lost among other Archaea either at least once (before the divergence of Acherontia) or at least twice (Acherontia-Stygia, Proteoarchaeota). That is in addition to additional losses throughout the Halobacterota and at the base of Diaforarchaea.

Although the identity and homology of ferredoxin homologs is fraught with ambiguity, there seems to have been a lot of tinkering in Methanomada involving the EhaPQR subunits that are MvhB-like ferredoxins. The Eha operon in many Methanobacteriales includes an extra gene compared to what was characterized in *Methanothermobacter thermautotrophicus* in [27]. Beyond EhaQRST homologs in Methanomada clades, additional tinkering can be traced to the origin of specific clades through subunit gains and losses. EhaP, while almost certainly present at the LMA, has been lost by Persephonarchaea. EhaI was similarly lost at the origin of Methanomicrobiales. EhaL is absent in Methanomicrobiales, Persephonarchaea, and Mnemosynellales. Together with the uncertainty of the Eha phylogeny root, it is impossible to determine whether EhaL was lost by these lineages, or invented at the origin of Methanomada, and by extension how ancient it is. Even more complicated is the case of EhaA that is missing from Methanomicrobiales and the smaller Methanobacteriales clade. The simplest scenario posits two individual loss events but the poorly resolved EhaA phylogeny (Supplementary Data 10) is also compatible with a more complex history: EhaA emerged at the base of Methanomada (or lost by Methanomicrobiales, Persephonarchaea, and Mnemosynellales in one or more events) and then transferred twice separately to Persephonarchaea and Mnemosynellales. The patchy distribution of EhaAILP could mean that they are non-essential for the function of the hydrogenase. Alternatively, the missing subunits in Methanomicrobiales, Persephonarchaea, and Mnemosynellales might be related to lack of coupling between $H_2/CO_2$ methanogenesis (with the exception of some Methanomicrobiales) and Fwd through Eha.

Evolution of Ehb

As mentioned in the Results & Discussion section, the most extraordinary event in the evolution of Ehb is the homologous recombination in the Methanococclaes. Even though we suspected its existence from the synteny and single-gene phylogenies (Supplementary Data 11), only rarely was the position of Methanococcales strongly supported. To investigate the signal difference among subunits and its origin, we reconstructed phylogenies for three different Ehb supermatrices: one with all subunits (Figure 3b), one corresponding to the Methanococcales syntenic cluster (EhbEFGHIKLMO, EhbJ was omitted due to its small size and inconsistent topology; Supplementary Figure 42), and one for the Methanococcales cluster without subunits that recover the monophyly of Methanomada (EhbEGHIKLM; Supplementary Figure 43). The Methanococcales cluster originated from a massive homologous recombination event related to the Acherontia, perhaps followed by further transfers or fast evolution of certain subunits that ended up determining the phylogenetic signal of the supermatrices.

We further tested the putative homologous recombination of Ehb in Methanococcales through our variant application of sCF (see Methods). In the sCF heatmap for the supermatrix of all Ehb subunits (Supplementary Figure 44b), the nodes with low sCF values in multiple subunits are 208 and 113. Node 208 simply reflects the uncertainty in the internal branching of Acherontia, although transfer events cannot be discounted. Node 113 corresponds to Methanomada; here it is subunits EhbBEGH that have lower values. With the exception of EhbB, all of them are part of the Methanococcales Ehb cluster. Most subunits of the cluster are also found together in the Euclidean clustering of the heatmap. This situation persists for the EhbEFGHIKLMO supermatrix (Supplementary Figure 43c) and only partially resolves for EhbEGHIKLM (Supplementary Figure 43d) where Methanococcales group with Acherontia, as for some subunits (especially EhbEG) their position remains inconsistent due to pervasive differences in the signal or further recombination, as described above. We observed the same behavior for gCF. For the complete Ehb supermatrix the gCF for Acherontia is 81.25 and for Methanomada 31.25 demonstrating the signal clash among subunits. For EhbEFGHIKLMO they drop to 77.78 and 22.22 respectively. Most of the signal for a monophyletic Methanomada has almost vanished apart from EhbFMO. Finally, in EhbEGHIKLM Methanococcales and Acherontia are monophyletic at 71.43 gCF.

The reasons for Ehb being retained by methylotrophic methanogens in the absence of the WLP is not entirely clear. In the case of Methanofastidiosales and Nuwarchaeales it seems like a last-ditch solution, since they lack any other hydrogenases[28]. The Acherontia ancestor had already lost Eha despite our inference that it was an $H_2/CO_2$ methanogen, given the presence of the WLP in Theionarchaea[7]. In the case of Verstraetearchaeota, Ehb has persisted in methylotrophic and $H_2/CO_2$ even though Eha had already been lost and is not found in other TACK methanogens either.

Notes on the evolution of WLP

Owing to the higher number of available genomes since two recent in-depth analyses of the WLP[7,29], we made two noteworthy observations. In the carbonyl branch article[30], an additional set of carbonyl branch enzymes in Syntropharchaeales was ignored, since they had been deemed to be too divergent and forming long branches in preliminary phylogenies. In this study, we identified a close relative of that set in a basal alkanotrophic Methanophagales[31]. The sequences from Syntropharchaeales and Methanophagales form a monophyletic clade but their branching position is not consistent among subunits. In contrast, the main Syntropharchaeales carbonyl branch sequences are found consistently within Halobacterota (Supplementary Figures 37, 40, 41). We hypothesize a participation of this carbonyl branch version in AAO but its exact function cannot be determined; it could function on propionyl-CoA or butyryl-CoA. The evolution and comparative genomics of the carbonyl branch will eventually need to be reexamined as more genomes become available. Concerning the $H_4$MPT branch, in the Mtd tree we found three Proteobacteria sequences forming a monophyletic branch inside Thorarchaeota, a clear interdomain transfer and the first such case for Mtd (Supplementary Figure 39). As the clade contained multiple (and non-identical) sequences, we could discount the possibility of binning errors. These Proteobacteria MAGs need to be studied in depth to determine the role of Mtd. Our preliminary hypothesis is that they are part of either a hybrid $H_4$MPT branch with Mtd instead of bMtd in methylotrophy, or that Mtd is substituting one of the functions of FolD in the $H_4$F branch.

| Borrel et al., 2019 | EggNOG accession number | Gao & Gupta, 2007 |
|---|---|---|
| m1_mcrA | arCOG04857 | MMP1559 |
| m2_mcrB | arCOG04860 | MMP1555 |
| m3_mcrG | arCOG04858 | MMP1558 |
| m4_Predicted rotamase | arCOG04900 | MMP0154 |
| m5_DUF2102 | arCOG04901 | MMP0337 |
| m6_DUF2112 | arCOG04903 | MMP0312 |
| m7_YjiL-like | arCOG02679 | MMP0608 |
| m8_DUF2113 | arCOG04904 | MMP0656 |
| m9_arCOG03226 | arCOG03226 | MMP0421 |
| m10_atwA | arCOG00185 | MMP0620 |
| m11_mcrC | arCOG03225 | MMP1557 |
| m12_mcrD | arCOG04859 | MMP1556 |
| m13_cfbD | arCOG04888 | MMP0428 |
| m14_cfbE | arCOG02822 | MMP0173 |
| m15_mtxX | arCOG00854 | MMP1346 |
| m16_AIR-synthase-related | arCOG00640 | MMP1531 |
| m17_Zn-ribbon protein | arCOG01116 | MMP1593 |
| m18_DUF2099 | arCOG04893 | MMP1644 |
| m19_DUF2117 | arCOG03231 | MMP0143 |
| m20_Gly-thioamidase | arCOG02882 | MMP1056 |
| m21_DUF2124 | arCOG04847 | MMP0563 |
| m22_DUF2098 | arCOG04846 | MMP0001 |
| m23_SolubleP-type ATPase | arCOG01579 | MMP1641 |
| m24_DUF2111 | arCOG04902 | MMP0311 |
| m25_arCOG04853 | arCOG04853 | MMP1704 |
| m26_DUF2114 | arCOG04866 | MMP1223 |
| m27_mtrA | arCOG03221 | MMP1564 |
| m28_mtrB | arCOG04867 | MMP1563 |
| m29_mtrC | arCOG04868 | MMP1562 |
| m30_mtrD | arCOG04869 | MMP1561 |
| m31_mtrE | arCOG04870 | MMP1560 |
| m32_arCOG04885 | arCOG04885 | MMP0642 |
| m33_Arg-methyltransferase | arCOG00950 | MMP1554 |
| m34_DUF1894 | arCOG04844 | MMP0698 |
| m35_DUF1890 | arCOG04845 | MMP0701 |
| m36_DUF2115 | arCOG03215 | MMP0665 |
| m37_DUF2119 | arCOG04894 | MMP1309 |
| m38_DUF2121 | arCOG03213 | MMP0021 |

**Supplementary Table 1.** Table of all 38 (m1-m38) methanogenesis marker genes following the set and numbering in Borrel et al.[28], their respective EggNOG accession numbers and *Methanococcus maripaludis* gene names as mentioned in [1]. Markers noted in red are not included in Gao & Gupta[1].

| Genome | Proposed Name | NCBI assembly | Genome Size (Mb) | Completeness (%) | Contamination (%) | GC (%) | Scaffolds (number) | Longest Scaffold (bp) | N50 (scaffolds) | Contigs (number) | Longest contig (bp) | N50 (contigs) | Coding density (%) | Predicted genes (number) |
|---|---|---|---|---|---|---|---|---|---|---|---|---|---|---|
| NRA7_88_Euryarchaeota_41_62 | *Ca.* Mnemosynella biddleae | GCA_002011165 | 1.50 | 96.08 | 0.65 | 41.18 | 11 | 341116 | 235752 | 11 | 341116 | 235752 | 91.98 | 1730 |
| NRA7_48_Euryarchaeota_41_21 | | GCA_002011155 | 1.47 | 96.08 | 0.65 | 41.14 | 21 | 238144 | 187849 | 21 | 238144 | 187849 | 92.84 | 1711 |
| nra7_reassver2_Euryarchaeota_45_14 | *Ca.* Mnemosynella hypogeia | GCA_014361165 | 1.29 | 93.46 | 1.31 | 44.82 | 38 | 106407 | 54584 | 41 | 106407 | 44894 | 93.21 | 1505 |
| nra7_reassver2_Euryarchaeota_47_10 | *Ca.* Mnemosynella sp. | GCA_014361185 | 0.90 | 81.05 | 0.65 | 46.69 | 73 | 68921 | 19803 | 88 | 68921 | 17146 | 95.50 | 1106 |
| NRA7_48_Candidatus_Bathyarchaeota_archaeon_52_381 | *Ca.* Hecatella orcuttiae | GCA_002011035 | 1.75 | 100 | 0.93 | 51.85 | 3 | 731052 | 535545 | 3 | 731052 | 535545 | 89.25 | 1881 |
| NRA7_88_Candidatus_Bathyarchaeota_52_49 | | GCA_002009985 | 1.74 | 100 | 0.93 | 51.93 | 9 | 1267205 | 1267205 | 13 | 769127 | 497978 | 89.37 | 1890 |
| NRA7_88_JdFR-13_38_55 | *Ca.* Scotarchaeum ottlingeri | GCA_002011075 | 1.70 | 93.20 | 0.97 | 37.67 | 7 | 1676113 | 1676113 | 11 | 691515 | 495125 | 90.26 | 1750 |
| NRA7_48_JdFR-14_42_64 | *Ca.* Geothermarchaeum rappei | GCA_002011085 | 1.58 | 94.17 | 0.97 | 41.83 | 1 | 1583546 | 1583546 | 1 | 1583546 | 1583546 | 92.28 | 1613 |
| NRA7_88_JdFR-14_42_10 | | GCA_002011085 | 1.44 | 93.20 | 1.29 | 42.03 | 204 | 69808 | 11815 | 243 | 40089 | 9032 | 92.17 | 1644 |
| NRA7_88_JdFR-16_51_899 | *Ca.* Pyrohabitans jungbluthii | GCA_002010065 | 1.99 | 98.13 | 2.80 | 51.20 | 15 | 620739 | 394757 | 15 | 620739 | 394757 | 92.48 | 2246 |
| NRA7_48_JdFR-16_51_665 | | GCA_002010065 | 1.17 | 51.04 | 1.87 | 51.06 | 82 | 64021 | 22921 | 82 | 64021 | 22921 | 92.30 | 1370 |
| NRA7_48_JdFR-17_50_1185 | | GCA_002011115 | 1.62 | 74.22 | 2.88 | 50.46 | 101 | 89324 | 20372 | 124 | 89324 | 19456 | 91.32 | 1880 |
| NRA7_48_JdFR-18_39_115 | | GCA_002011125 | 2.19 | 98.13 | 1.87 | 39.11 | 19 | 1917514 | 1917514 | 24 | 1201297 | 1201297 | 91.69 | 2424 |

**Supplementary Table 2.** Summary statistics of genomes binned in this study and their proposed names (see Descriptions of proposed taxa). Bins named NRA7_48 and NRA7_88 correspond to SRR3723048/U1362A and SRR3732688/U1362B from [32], while the "nra7_reassver2" bins originate to a co-assembly of the three Shengli metagenomes[33].

| taxon | assembly | coordinates | location | isolation_source | metagenome_source |
|---|---|---|---|---|---|
| Euryarchaeota archaeon JdFR-21 * | GCA_002011165 | 47.76 N 127.76 W | Pacific Ocean | basaltic crustal fluids | subsurface metagenome |
| Archaeoglobi archaeon * | GCA_014361165 | 38.15 N 118.5 E | China: Dongying | Shengli oilfield at Shandong province | oil production facility metagenome |
| Archaeoglobi archaeon * | GCA_014361185 | 38.15 N 118.5 E | China: Dongying | Shengli oilfield at Shandong province | oil production facility metagenome |
| Candidatus Bathyarchaeota archaeon ex4484 218 | GCA_002254975 | 27.0388 N 111.2456 W | USA: Guaymas Basin, Gulf of California | deep-sea hydrothermal vent sediments from dive 4484 | marine sediment metagenome |
| Candidatus Bathyarchaeota archaeon | GCA_003661965 | 29.52416667 N 113.57000000 W | Mexico: Guaymas Basin, Gulf of California | deep-sea hydrothermal vent sediments from dive 4569_2 depth 12-15 cm | marine sediment metagenome |
| Candidatus Bathyarchaeota archaeon JdFR-11 * | GCA_002011035 | 47.76 N 127.76 W | Pacific Ocean | basaltic crustal fluids | subsurface metagenome |
| Candidatus Bathyarchaeota archaeon | GCA_003662515 | 29.52416667 N 113.57000000 W | Mexico: Guaymas Basin, Gulf of California | deep-sea hydrothermal vent sediments | marine sediment metagenome |
| Candidatus Bathyarchaeota archaeon | GCA_003662485 | 29.52416667 N 113.57000000 W | Mexico: Guaymas Basin, Gulf of California | deep-sea hydrothermal vent sediments from dive 4571_4 depth 12-15 cm | marine sediment metagenome |
| Candidatus Bathyarchaeota archaeon | GCA_003662305 | 29.52416667 N 113.57000000 W | Mexico: Guaymas Basin, Gulf of California | deep-sea hydrothermal vent sediments from dive 4569_2 depth 21-24 cm | marine sediment metagenome |
| Candidatus Bathyarchaeota archaeon B25 | GCA_001593855 | 27.0119 N 111.4054 W | Mexico: Guaymas Basin, Gulf of California | marine sediment sample collected by push cores at a deep sea vent site during cruise AT15-25 on Alvin dive 4358 | hydrothermal vent metagenome |
| Candidatus Thorarchaeota archaeon | GCA_004525055 | 44.0552806 N 28.6053 E | Romania: Tekirghiol | lake sediment | sediment metagenome |
| Candidatus Thorarchaeota archaeon | GCA_004524305 | 44.0552806 N 28.6053 E | Romania: Tekirghiol | lake sediment | sediment metagenome |
| Candidatus Thorarchaeota archaeon | GCA_003345595 | 41.3778 N 82.5108 W | USA: Ohio, Lake Erie, Old Woman Creek | freshwater wetland soil | wetland metagenome |
| Candidatus Thorarchaeota archaeon | GCA_003345555 | 41.3778 N 82.5108 W | USA: Ohio, Lake Erie, Old Woman Creek | freshwater wetland soil | wetland metagenome |
| Candidatus Thorarchaeota archaeon | GCA_004524445 | 44.0552806 N 28.6053 E | Romania: Tekirghiol | lake sediment | sediment metagenome |
| Candidatus Thorarchaeota archaeon | GCA_003662765 | 29.52416667 N 113.57000000 W | Mexico: Guaymas Basin, Gulf of California | deep-sea hydrothermal vent sediments from dive 4571_4 depth 0-3 cm | marine sediment metagenome |
| Candidatus Thorarchaeota archaeon SMTZ1-83 | GCA_001563325 | 34.7414 N 77.1234 W | USA: White Oak River Estuary, North Carolina | Sulfate-methane transisition zone estuary sediments 16-26 cm | sediment metagenome |
| Candidatus Thorarchaeota archaeon | GCA_003662775 | 27.015 N 111.379 W | Mexico: Guaymas Basin, Gulf of California | deep-sea hydrothermal vent sediments from dive 4569_9 depth 0-3 cm | marine sediment metagenome |
| Candidatus Thorarchaeota archaeon | GCA_004376265 | 26.28 N 86.81 W | Atlantic Ocean | deep sea sediments associated with petroleum seepage | marine sediment metagenome |
| Candidatus Thorarchaeota archaeon AB_25 | GCA_001940705 | 56.103333 N 10.45783299 E | Denmark: Aarhus Bay, Baltic Sea | Marine sediment | marine sediment metagenome |
| Candidatus Thorarchaeota archaeon MP11T_1 | GCA_002825515 | 22.4992 N 114.0276 E | Hong Kong: Mai Po Nature Reserve | mangrove wetland sediments | sediment metagenome |
| Candidatus Thorarchaeota archaeon SMTZ1-45 | GCA_001563335 | 34.7414 N 77.1234 W | USA: White Oak River Estuary, North Carolina | Sulfate-methane transisition zone estuary sediments 16-26 cm | sediment metagenome |
| Candidatus Thorarchaeota archaeon MP9T_1 | GCA_002825535 | 22.4979 N 114.0295 E | Hong Kong: Mai Po Nature Reserve | sediment (20-25 cm) from mangrove covering field | sediment metagenome |
| Candidatus Thorarchaeota archaeon | GCA_003345545 | 41.3778 N 82.5108 W | USA: Ohio, Lake Erie, Old Woman Creek | freshwater wetland soil | wetland metagenome |
| Candidatus Thorarchaeota archaeon | GCA_004524595 | 44.6085139 N 27.331375 E | Romania: Amara | lake sediment | sediment metagenome |
| Candidatus Thorarchaeota archaeon | GCA_004525375 | 63.29 N 19.49 E | Sweden: Bothnian Sea | Bothnian Sea sediment, site NB8 | marine sediment metagenome |
| Candidatus Thorarchaeota archaeon | GCA_004525645 | 63.29 N 19.49 E | Sweden: Bothnian Sea | Bothnian Sea sediment, site NB8 | marine sediment metagenome |
| Candidatus Thorarchaeota archaeon | GCA_004524565 | 44.6085139 N 27.331375 E | Romania: Amara | lake sediment | sediment metagenome |
| Candidatus Thorarchaeota archaeon | GCA_004524435 | 44.0552806 N 28.6053 E | Romania: Tekirghiol | lake sediment | sediment metagenome |

* genomes from metagenomes reassembled and rebinned in this study

**Supplementary Table 3.** Detailed metadata (accessions, location, environment) for the Mnemosynellales, Hecatellales, and Thorarchaeota MAGs in Figure 2e. For convenience, since there are too many entries, the metadata for Mnemosynellales and Hecatellales 16S sequences are presented in Supplementary Data 5.

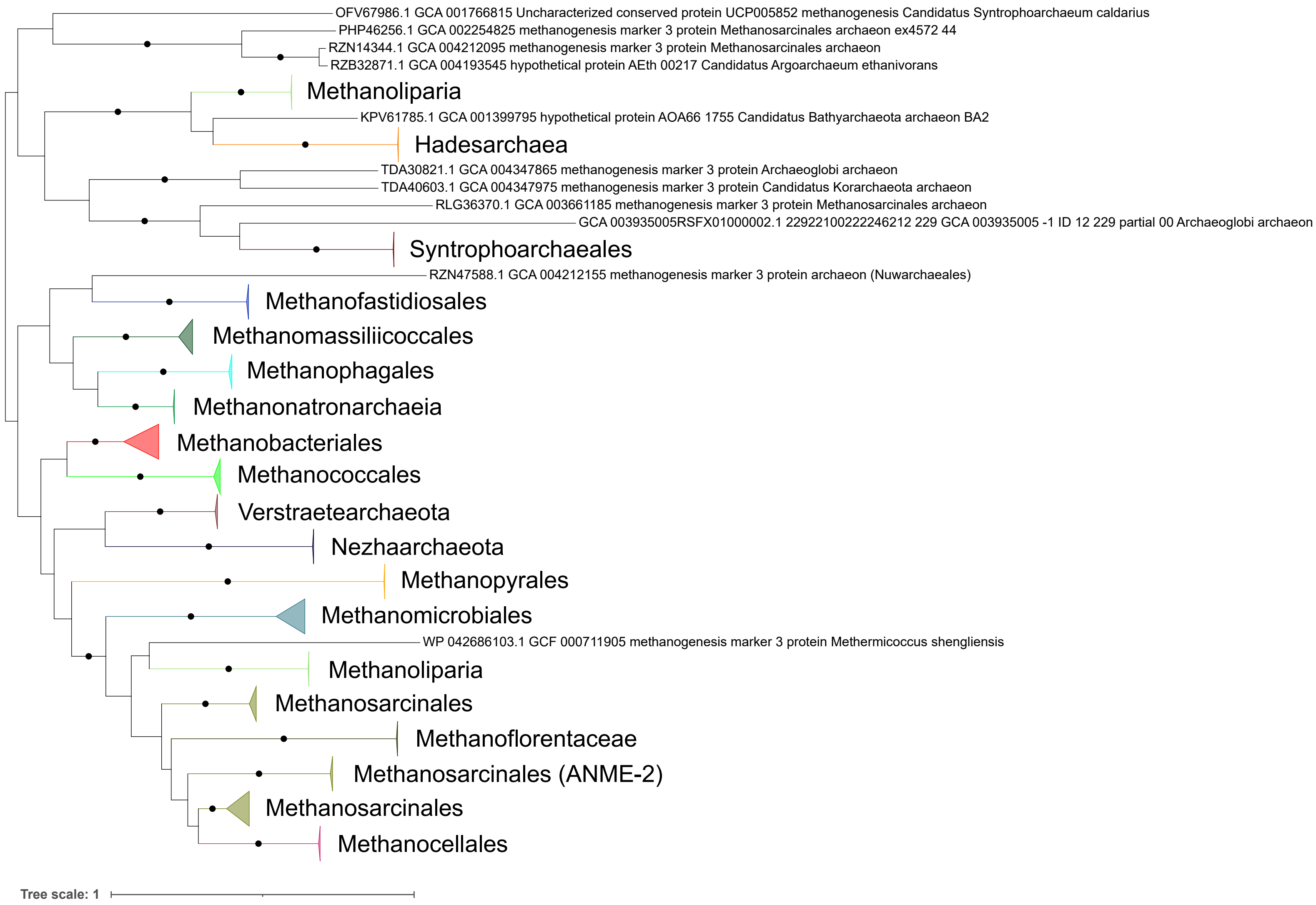
OFV67986.1 GCA 001766815 Uncharacterized conserved protein UCP005852 methanogenesis Candidatus Syntrophoarchaeum caldarius
PHP46256.1 GCA 002254825 methanogenesis marker 3 protein Methanosarcinales archaeon ex4572 44
RZN14344.1 GCA 004212095 methanogenesis marker 3 protein Methanosarcinales archaeon
RZB32871.1 GCA 004193545 hypothetical protein AEth 00217 Candidatus Argoarchaeum ethanivorans
Methanoliparia
KPV61785.1 GCA 001399795 hypothetical protein AOA66 1755 Candidatus Bathyarchaeota archaeon BA2
Hadesarchaea
TDA30821.1 GCA 004347865 methanogenesis marker 3 protein Archaeoglobi archaeon
TDA40603.1 GCA 004347975 methanogenesis marker 3 protein Candidatus Korarchaeota archaeon
RLG36370.1 GCA 003661185 methanogenesis marker 3 protein Methanosarcinales archaeon
GCA 003935005RSFX01000002.1 22922100222246212 229 GCA 003935005 -1 ID 12 229 partial 00 Archaeoglobi archaeon
Syntrophoarchaeales
RZN47588.1 GCA 004212155 methanogenesis marker 3 protein archaeon (Nuwarchaeales)
Methanofastidiosales
Methanomassiliicoccales
Methanophagales
Methanonatronarchaeia
Methanobacteriales
Methanococcales
Verstraetearchaeota
Nezhaarchaeota
Methanopyrales
Methanomicrobiales
WP 042686103.1 GCF 000711905 methanogenesis marker 3 protein Methermicoccus shengliensis
Methanoliparia
Methanosarcinales
Methanoflorentaceae
Methanosarcinales (ANME-2)
Methanosarcinales
Methanocellales
Tree scale: 1

**Supplementary Figure 1.** ML phylogeny of methanogenesis marker m4. Black circles indicate strongly supported branches (ultrafast bootstrap >=95, aLRT SH-like >=80).

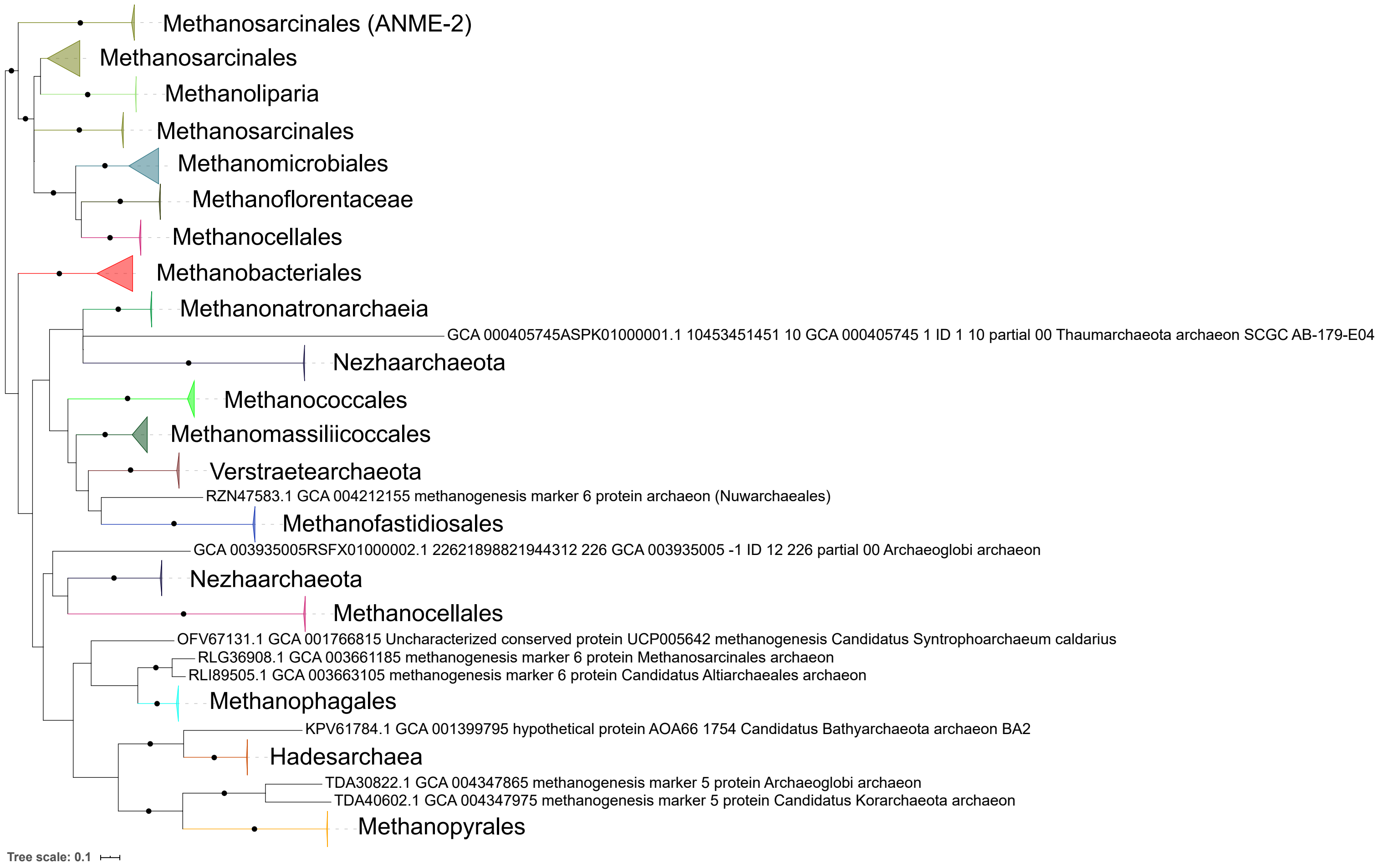

Methanosarcinales (ANME-2)
Methanosarcinales
Methanoliparia
Methanosarcinales
Methanomicrobiales
Methanoflorentaceae
Methanocellales
Methanobacteriales
Methanonatronarchaeia
GCA 000405745ASPK01000001.1 10453451451 10 GCA 000405745 1 ID 1 10 partial 00 Thaumarchaeota archaeon SCGC AB-179-E04
Nezhaarchaeota
Methanococcales
Methanomassiliicoccales
Verstraetearchaeota
RZN47583.1 GCA 004212155 methanogenesis marker 6 protein archaeon (Nuwarchaeales)
Methanofastidiosales
GCA 003935005RSFX01000002.1 22621898821944312 226 GCA 003935005 -1 ID 12 226 partial 00 Archaeoglobi archaeon
Nezhaarchaeota
Methanocellales
OFV67131.1 GCA 001766815 Uncharacterized conserved protein UCP005642 methanogenesis Candidatus Syntrophoarchaeum caldarius
RLG36908.1 GCA 003661185 methanogenesis marker 6 protein Methanosarcinales archaeon
RLI89505.1 GCA 003663105 methanogenesis marker 6 protein Candidatus Altiarchaeales archaeon
Methanophagales
KPV61784.1 GCA 001399795 hypothetical protein AOA66 1754 Candidatus Bathyarchaeota archaeon BA2
Hadesarchaea
TDA30822.1 GCA 004347865 methanogenesis marker 5 protein Archaeoglobi archaeon
TDA40602.1 GCA 004347975 methanogenesis marker 5 protein Candidatus Korarchaeota archaeon
Methanopyrales
Tree scale: 0.1

**Supplementary Figure 2.** ML phylogeny of methanogenesis marker m5. Black circles indicate strongly supported branches (ultrafast bootstrap >=95, aLRT SH-like >=80).

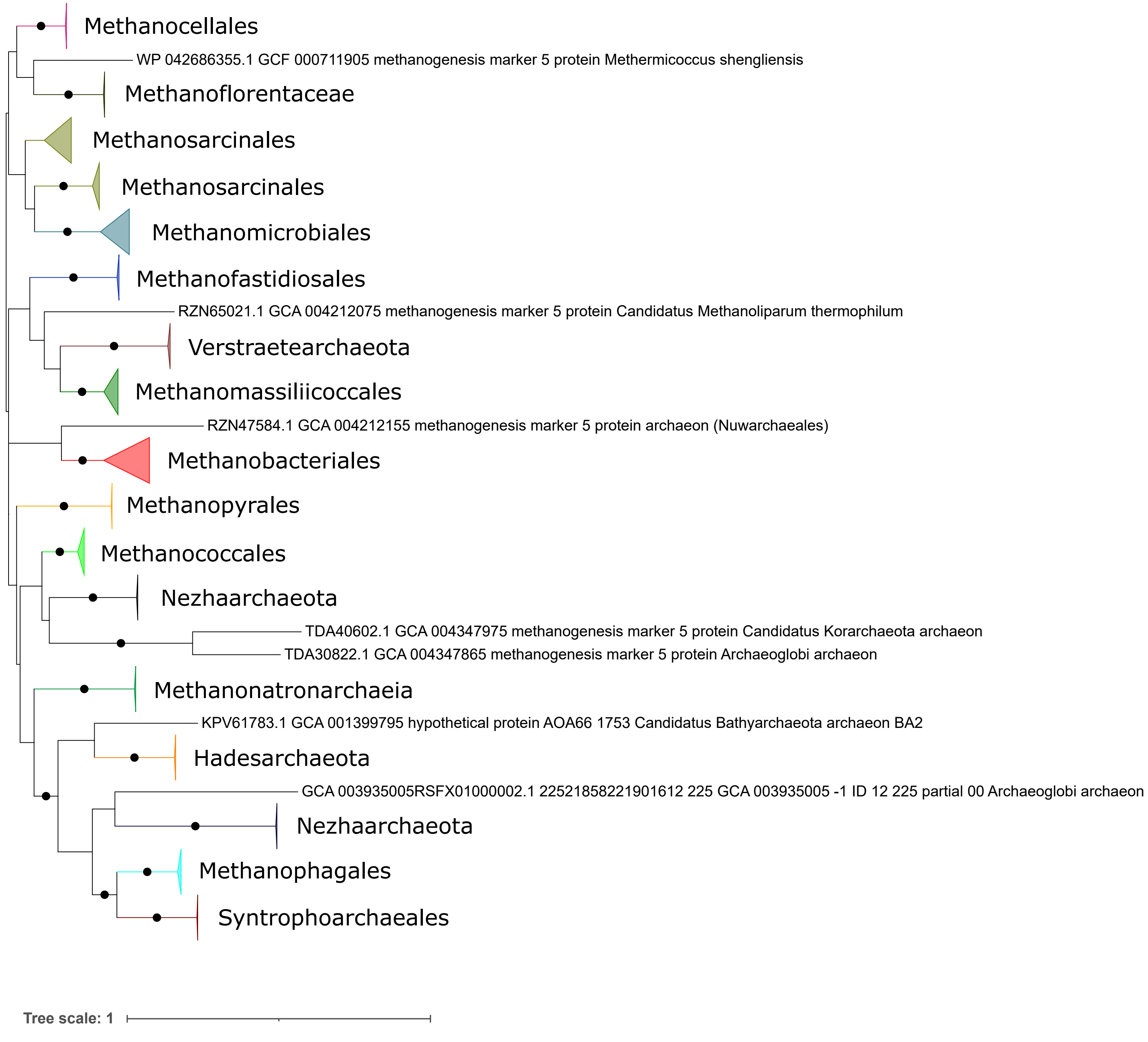
Methanocellales
WP 042686355.1 GCF 000711905 methanogenesis marker 5 protein Methermicoccus shengliensis
Methanoflorentaceae
Methanosarcinales
Methanosarcinales
Methanomicrobiales
Methanofastidiosales
RZN65021.1 GCA 004212075 methanogenesis marker 5 protein Candidatus Methanoliparum thermophilum
Verstraetearchaeota
Methanomassiliicoccales
RZN47584.1 GCA 004212155 methanogenesis marker 5 protein archaeon (Nuwarchaeales)
Methanobacteriales
Methanopyrales
Methanococcales
Nezhaarchaeota
TDA40602.1 GCA 004347975 methanogenesis marker 5 protein Candidatus Korarchaeota archaeon
TDA30822.1 GCA 004347865 methanogenesis marker 5 protein Archaeoglobi archaeon
Methanonatronarchaeia
KPV61783.1 GCA 001399795 hypothetical protein AOA66 1753 Candidatus Bathyarchaeota archaeon BA2
Hadesarchaeota
GCA 003935005RSFX01000002.1 22521858221901612 225 GCA 003935005 -1 ID 12 225 partial 00 Archaeoglobi archaeon
Nezhaarchaeota
Methanophagales
Syntrophoarchaeales
Tree scale: 1

**Supplementary Figure 3.** ML phylogeny of methanogenesis marker m6. Black circles indicate strongly supported branches (ultrafast bootstrap >=95, aLRT SH-like >=80).

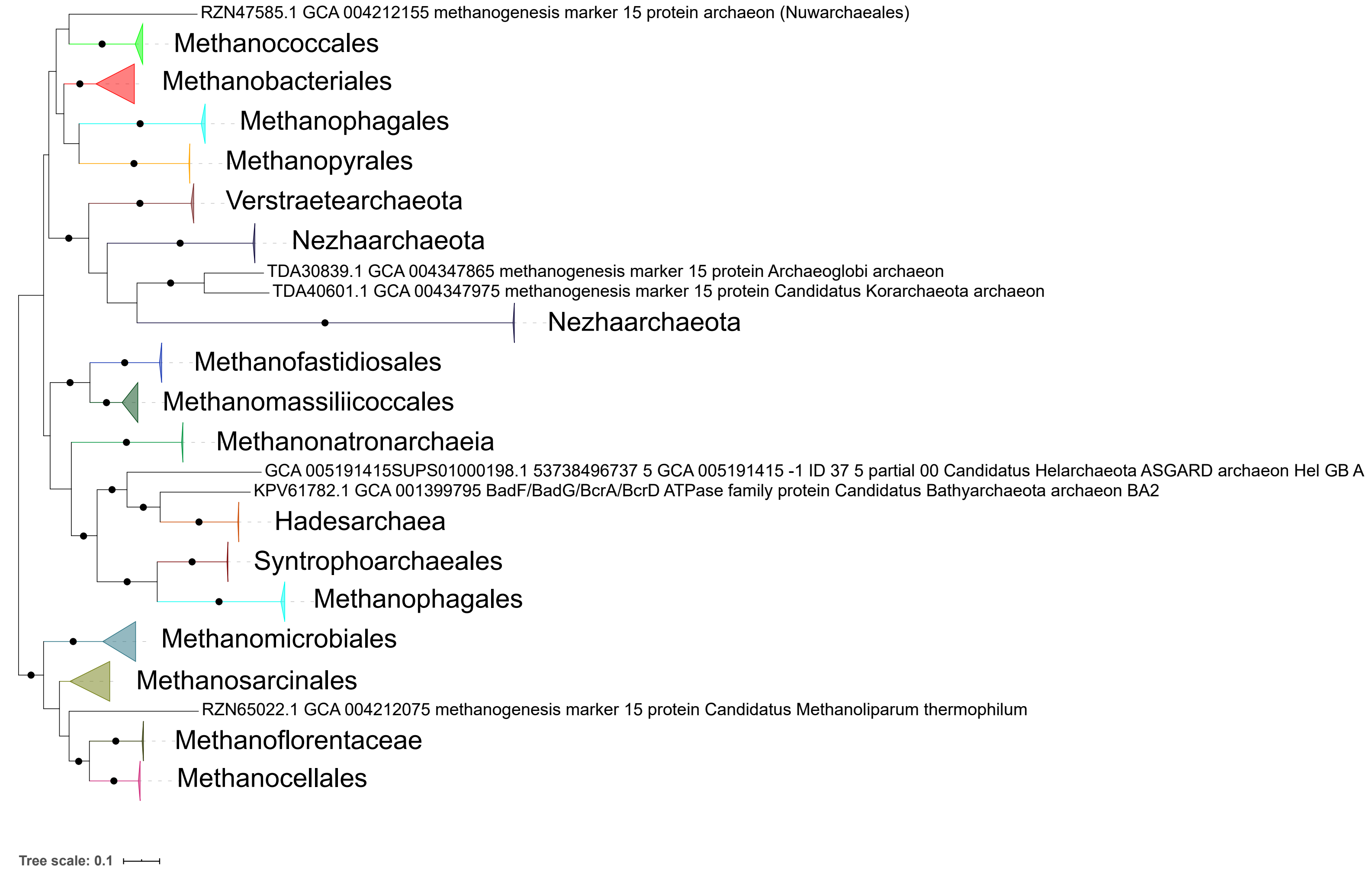

RZN47585.1 GCA 004212155 methanogenesis marker 15 protein archaeon (Nuwarchaeales)
Methanococcales
Methanobacteriales
Methanophagales
Methanopyrales
Verstraetearchaeota
Nezhaarchaeota
TDA30839.1 GCA 004347865 methanogenesis marker 15 protein Archaeoglobi archaeon
TDA40601.1 GCA 004347975 methanogenesis marker 15 protein Candidatus Korarchaeota archaeon
Nezhaarchaeota
Methanofastidiosales
Methanomassiliicoccales
Methanonatronarchaeia
GCA 005191415SUPS01000198.1 53738496737 5 GCA 005191415 -1 ID 37 5 partial 00 Candidatus Helarchaeota ASGARD archaeon Hel GB A
KPV61782.1 GCA 001399795 BadF/BadG/BcrA/BcrD ATPase family protein Candidatus Bathyarchaeota archaeon BA2
Hadesarchaea
Syntrophoarchaeales
Methanophagales
Methanomicrobiales
Methanosarcinales
RZN65022.1 GCA 004212075 methanogenesis marker 15 protein Candidatus Methanoliparum thermophilum
Methanoflorentaceae
Methanocellales
Tree scale: 0.1

**Supplementary Figure 4.** ML phylogeny of methanogenesis marker m7. Black circles indicate strongly supported branches (ultrafast bootstrap >=95, aLRT SH-like >=80).

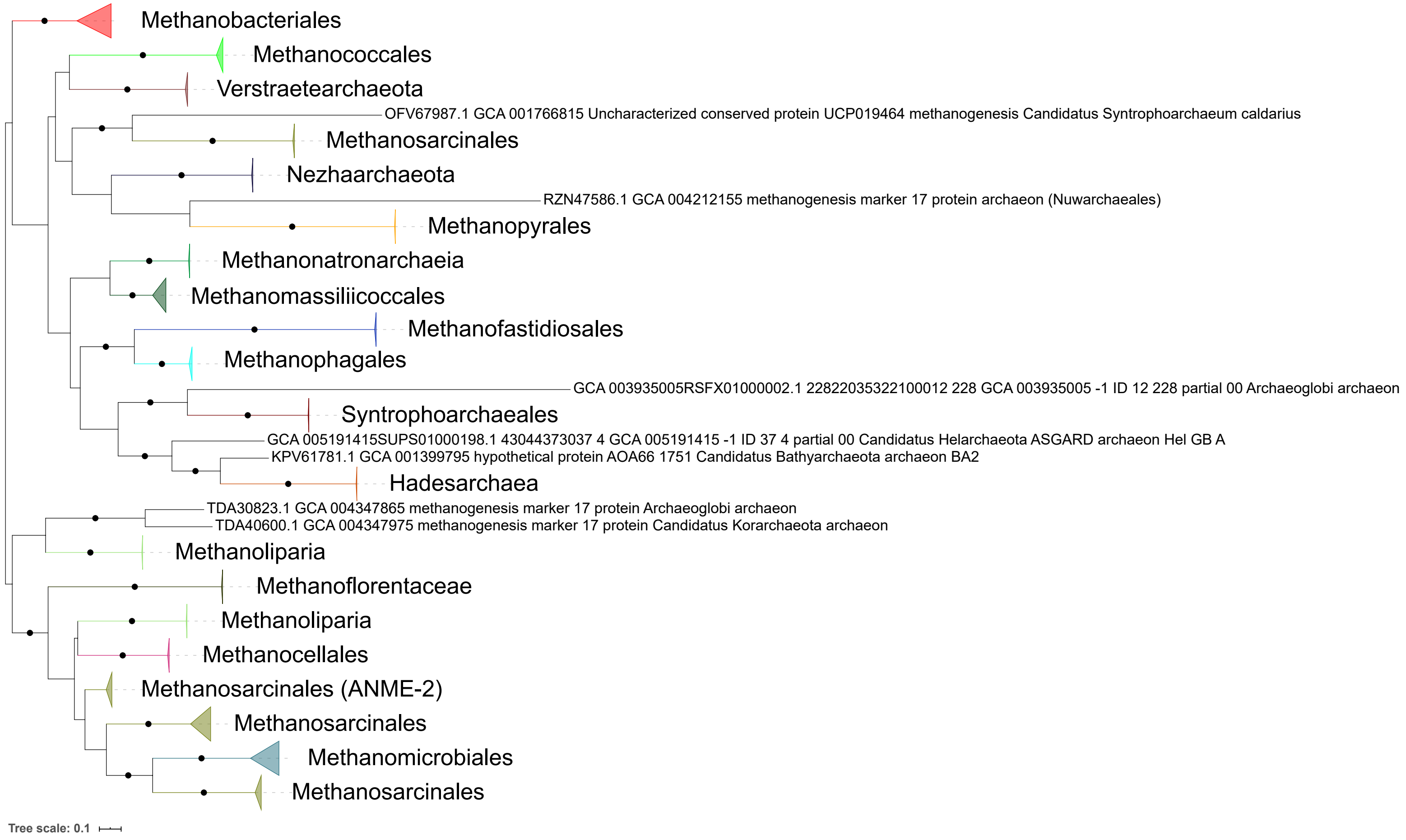

Methanobacteriales
Methanococcales
Verstraetearchaeota
OFV67987.1 GCA 001766815 Uncharacterized conserved protein UCP019464 methanogenesis Candidatus Syntrophoarchaeum caldarius
Methanosarcinales
Nezhaarchaeota
RZN47586.1 GCA 004212155 methanogenesis marker 17 protein archaeon (Nuwarchaeales)
Methanopyrales
Methanonatronarchaeia
Methanomassiliicoccales
Methanofastidiosales
Methanophagales
GCA 003935005RSFX01000002.1 22822035322100012 228 GCA 003935005 -1 ID 12 228 partial 00 Archaeoglobi archaeon
Syntrophoarchaeales
GCA 005191415SUPS01000198.1 43044373037 4 GCA 005191415 -1 ID 37 4 partial 00 Candidatus Helarchaeota ASGARD archaeon Hel GB A
KPV61781.1 GCA 001399795 hypothetical protein AOA66 1751 Candidatus Bathyarchaeota archaeon BA2
Hadesarchaea
TDA30823.1 GCA 004347865 methanogenesis marker 17 protein Archaeoglobi archaeon
TDA40600.1 GCA 004347975 methanogenesis marker 17 protein Candidatus Korarchaeota archaeon
Methanoliparia
Methanoflorentaceae
Methanoliparia
Methanocellales
Methanosarcinales (ANME-2)
Methanosarcinales
Methanomicrobiales
Methanosarcinales
Tree scale: 0.1

**Supplementary Figure 5.** ML phylogeny of methanogenesis marker m8. Black circles indicate strongly supported branches (ultrafast bootstrap >=95, aLRT SH-like >=80).

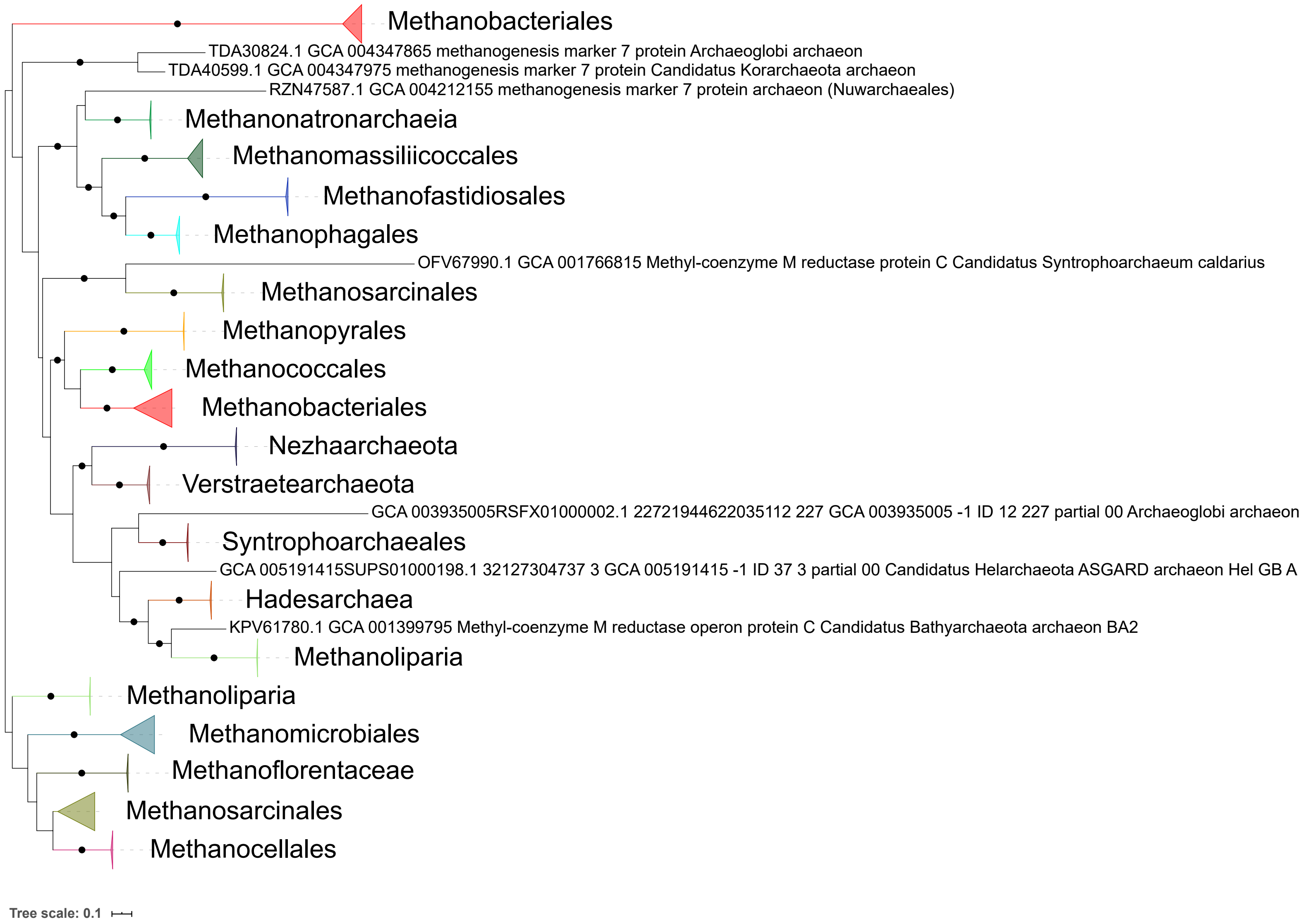

Methanobacteriales
TDA30824.1 GCA 004347865 methanogenesis marker 7 protein Archaeoglobi archaeon
TDA40599.1 GCA 004347975 methanogenesis marker 7 protein Candidatus Korarchaeota archaeon
RZN47587.1 GCA 004212155 methanogenesis marker 7 protein archaeon (Nuwarchaeales)
Methanonatronarchaeia
Methanomassiliicoccales
Methanofastidiosales
Methanophagales
OFV67990.1 GCA 001766815 Methyl-coenzyme M reductase protein C Candidatus Syntrophoarchaeum caldarius
Methanosarcinales
Methanopyrales
Methanococcales
Methanobacteriales
Nezhaarchaeota
Verstraetearchaeota
GCA 003935005RSFX01000002.1 22721944622035112 227 GCA 003935005 -1 ID 12 227 partial 00 Archaeoglobi archaeon
Syntrophoarchaeales
GCA 005191415SUPS01000198.1 32127304737 3 GCA 005191415 -1 ID 37 3 partial 00 Candidatus Helarchaeota ASGARD archaeon Hel GB A
Hadesarchaea
KPV61780.1 GCA 001399795 Methyl-coenzyme M reductase operon protein C Candidatus Bathyarchaeota archaeon BA2
Methanoliparia
Methanoliparia
Methanomicrobiales
Methanoflorentaceae
Methanosarcinales
Methanocellales
Tree scale: 0.1

**Supplementary Figure 6.** ML phylogeny of methanogenesis marker m9. Black circles indicate strongly supported branches (ultrafast bootstrap >=95, aLRT SH-like >=80).

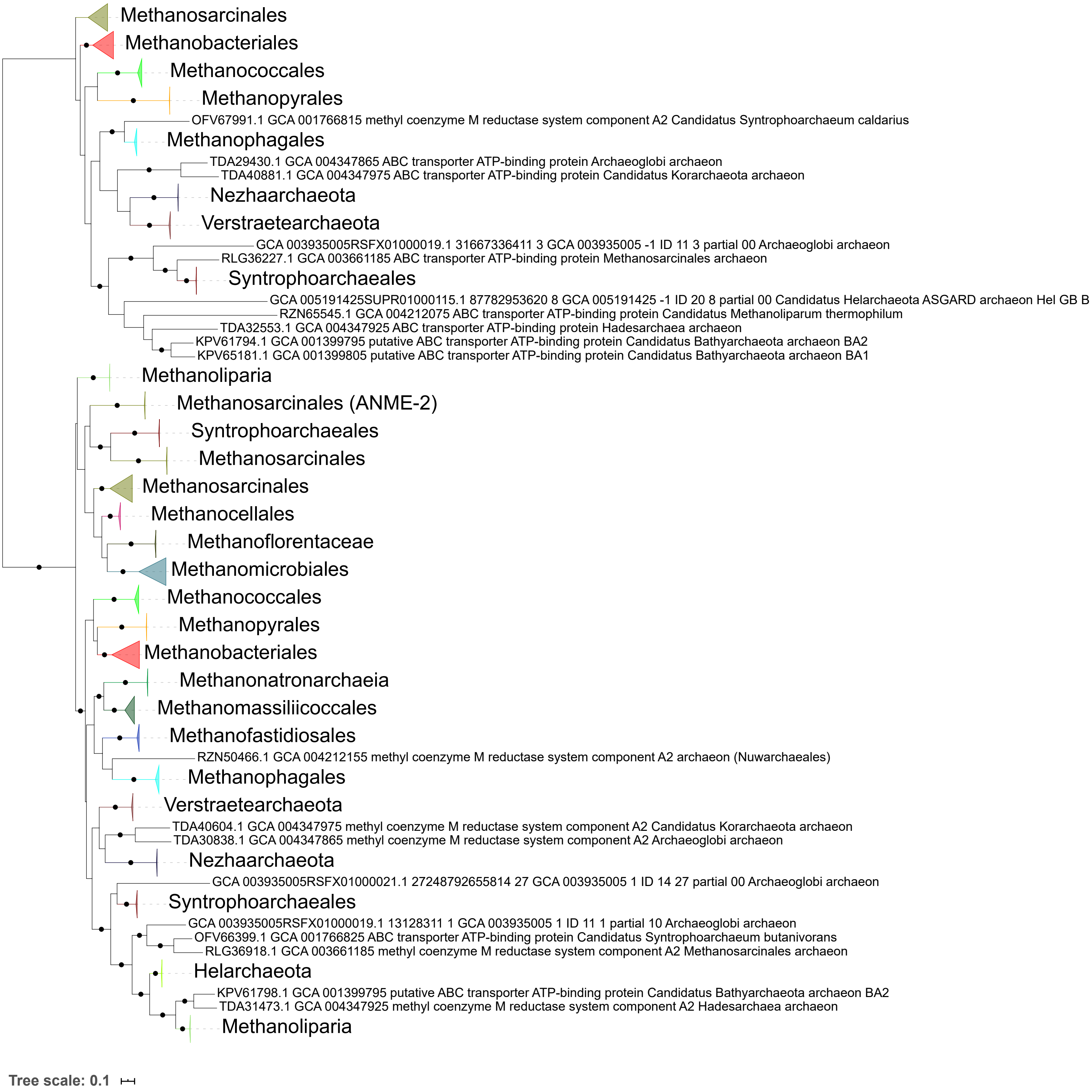

Methanosarcinales
Methanobacteriales
Methanococcales
Methanopyrales
OFV67991.1 GCA 001766815 methyl coenzyme M reductase system component A2 Candidatus Syntrophoarchaeum caldarius
Methanophagales
TDA29430.1 GCA 004347865 ABC transporter ATP-binding protein Archaeoglobi archaeon
TDA40881.1 GCA 004347975 ABC transporter ATP-binding protein Candidatus Korarchaeota archaeon
Nezhaarchaeota
Verstraetearchaeota
GCA 003935005RSFX01000019.1 31667336411 3 GCA 003935005 -1 ID 11 3 partial 00 Archaeoglobi archaeon
RLG36227.1 GCA 003661185 ABC transporter ATP-binding protein Methanosarcinales archaeon
Syntrophoarchaeales
GCA 005191425SUPR01000115.1 87782953620 8 GCA 005191425 -1 ID 20 8 partial 00 Candidatus Helarchaeota ASGARD archaeon Hel GB B
RZN65545.1 GCA 004212075 ABC transporter ATP-binding protein Candidatus Methanoliparum thermophilum
TDA32553.1 GCA 004347925 ABC transporter ATP-binding protein Hadesarchaea archaeon
KPV61794.1 GCA 001399795 putative ABC transporter ATP-binding protein Candidatus Bathyarchaeota archaeon BA2
KPV65181.1 GCA 001399805 putative ABC transporter ATP-binding protein Candidatus Bathyarchaeota archaeon BA1
Methanoliparia
Methanosarcinales (ANME-2)
Syntrophoarchaeales
Methanosarcinales
Methanosarcinales
Methanocellales
Methanoflorentaceae
Methanomicrobiales
Methanococcales
Methanopyrales
Methanobacteriales
Methanonatronarchaeia
Methanomassiliicoccales
Methanofastidiosales
RZN50466.1 GCA 004212155 methyl coenzyme M reductase system component A2 archaeon (Nuwarchaeales)
Methanophagales
Verstraetearchaeota
TDA40604.1 GCA 004347975 methyl coenzyme M reductase system component A2 Candidatus Korarchaeota archaeon
TDA30838.1 GCA 004347865 methyl coenzyme M reductase system component A2 Archaeoglobi archaeon
Nezhaarchaeota
GCA 003935005RSFX01000021.1 27248792655814 27 GCA 003935005 1 ID 14 27 partial 00 Archaeoglobi archaeon
Syntrophoarchaeales
GCA 003935005RSFX01000019.1 13128311 1 GCA 003935005 1 ID 11 1 partial 10 Archaeoglobi archaeon
OFV66399.1 GCA 001766825 ABC transporter ATP-binding protein Candidatus Syntrophoarchaeum butanivorans
RLG36918.1 GCA 003661185 methyl coenzyme M reductase system component A2 Methanosarcinales archaeon
Helarchaeota
KPV61798.1 GCA 001399795 putative ABC transporter ATP-binding protein Candidatus Bathyarchaeota archaeon BA2
TDA31473.1 GCA 004347925 methyl coenzyme M reductase system component A2 Hadesarchaea archaeon
Methanoliparia
Tree scale: 0.1

**Supplementary Figure 7.** ML phylogeny of methanogenesis marker m10. Black circles indicate strongly supported branches (ultrafast bootstrap >=95, aLRT SH-like >=80).

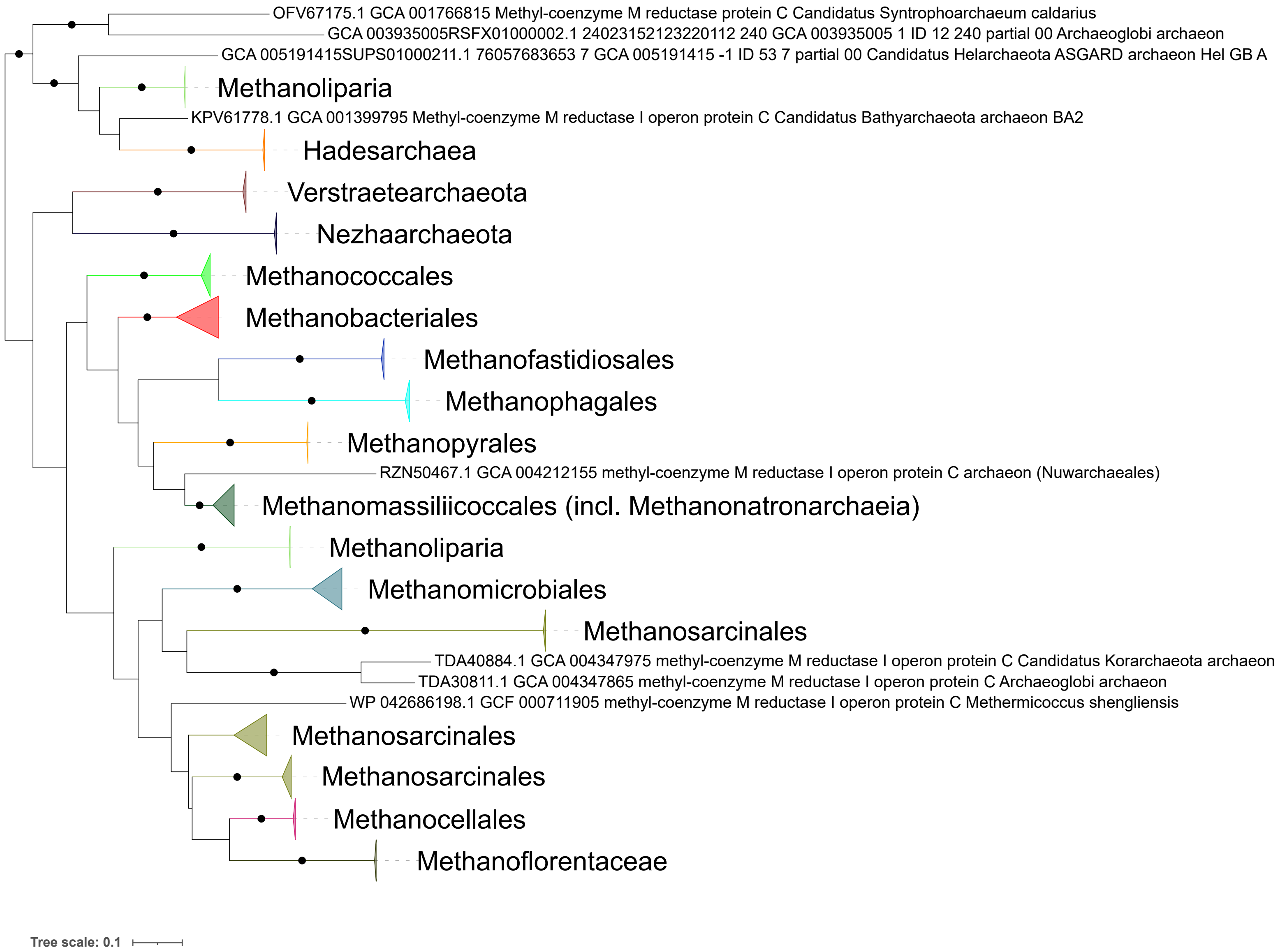
OFV67175.1 GCA 001766815 Methyl-coenzyme M reductase protein C Candidatus Syntrophoarchaeum caldarius
GCA 003935005RSFX01000002.1 24023152123220112 240 GCA 003935005 1 ID 12 240 partial 00 Archaeoglobi archaeon
GCA 005191415SUPS01000211.1 76057683653 7 GCA 005191415 -1 ID 53 7 partial 00 Candidatus Helarchaeota ASGARD archaeon Hel GB A
Methanoliparia
KPV61778.1 GCA 001399795 Methyl-coenzyme M reductase I operon protein C Candidatus Bathyarchaeota archaeon BA2
Hadesarchaea
Verstraetearchaeota
Nezhaarchaeota
Methanococcales
Methanobacteriales
Methanofastidiosales
Methanophagales
Methanopyrales
RZN50467.1 GCA 004212155 methyl-coenzyme M reductase I operon protein C archaeon (Nuwarchaeales)
Methanomassiliicoccales (incl. Methanonatronarchaeia)
Methanoliparia
Methanomicrobiales
Methanosarcinales
TDA40884.1 GCA 004347975 methyl-coenzyme M reductase I operon protein C Candidatus Korarchaeota archaeon
TDA30811.1 GCA 004347865 methyl-coenzyme M reductase I operon protein C Archaeoglobi archaeon
WP 042686198.1 GCF 000711905 methyl-coenzyme M reductase I operon protein C Methermicoccus shengliensis
Methanosarcinales
Methanosarcinales
Methanocellales
Methanoflorentaceae
Tree scale: 0.1

**Supplementary Figure 8.** ML phylogeny of methanogenesis marker m11. Black circles indicate strongly supported branches (ultrafast bootstrap >=95, aLRT SH-like >=80).

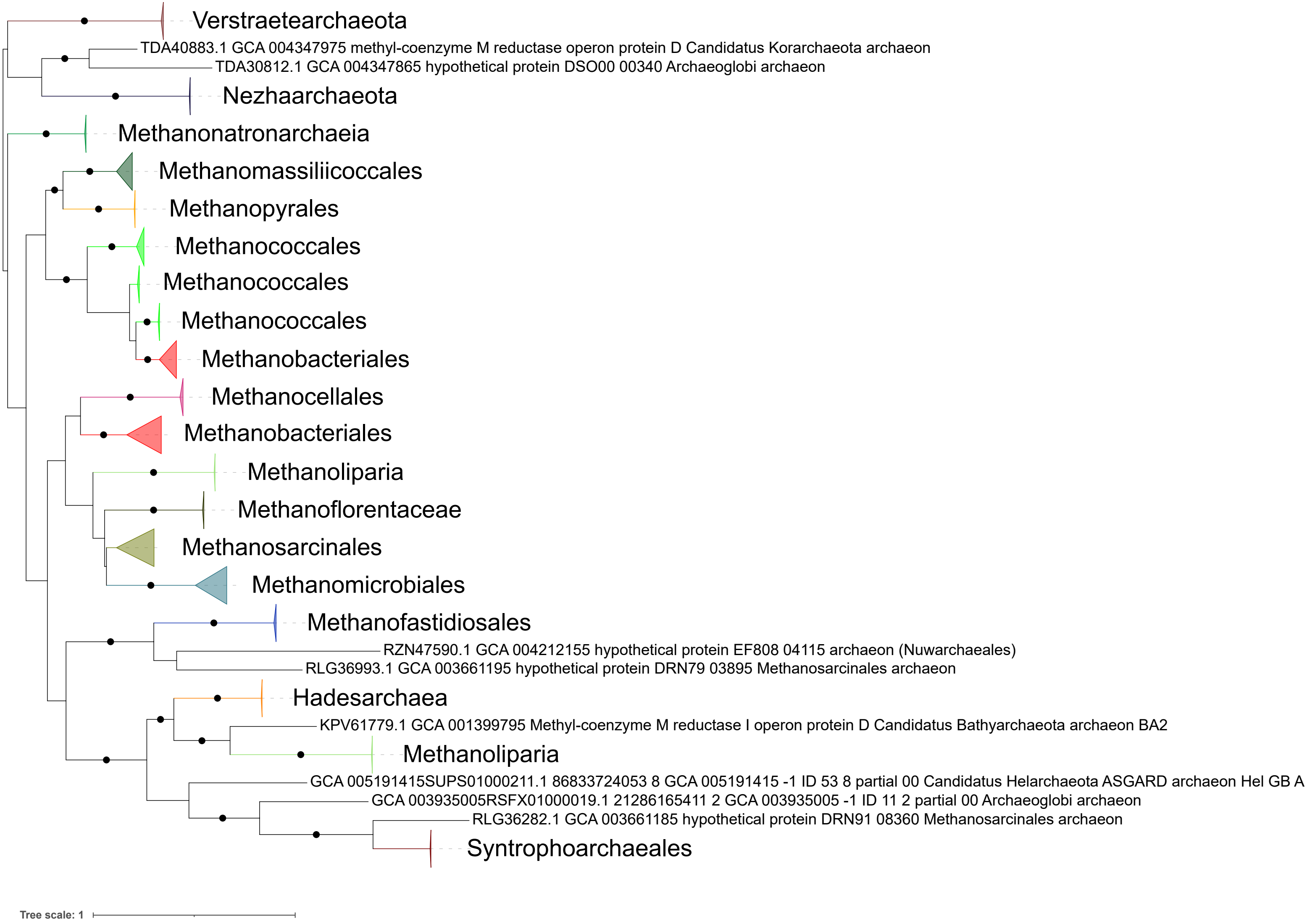
Verstraetearchaeota
TDA40883.1 GCA 004347975 methyl-coenzyme M reductase operon protein D Candidatus Korarchaeota archaeon
TDA30812.1 GCA 004347865 hypothetical protein DSO00 00340 Archaeoglobi archaeon
Nezhaarchaeota
Methanonatronarchaeia
Methanomassiliicoccales
Methanopyrales
Methanococcales
Methanococcales
Methanococcales
Methanobacteriales
Methanocellales
Methanobacteriales
Methanoliparia
Methanoflorentaceae
Methanosarcinales
Methanomicrobiales
Methanofastidiosales
RZN47590.1 GCA 004212155 hypothetical protein EF808 04115 archaeon (Nuwarchaeales)
RLG36993.1 GCA 003661195 hypothetical protein DRN79 03895 Methanosarcinales archaeon
Hadesarchaea
KPV61779.1 GCA 001399795 Methyl-coenzyme M reductase I operon protein D Candidatus Bathyarchaeota archaeon BA2
Methanoliparia
GCA 005191415SUPS01000211.1 86833724053 8 GCA 005191415 -1 ID 53 8 partial 00 Candidatus Helarchaeota ASGARD archaeon Hel GB A
GCA 003935005RSFX01000019.1 21286165411 2 GCA 003935005 -1 ID 11 2 partial 00 Archaeoglobi archaeon
RLG36282.1 GCA 003661185 hypothetical protein DRN91 08360 Methanosarcinales archaeon
Syntrophoarchaeales
Tree scale: 1

**Supplementary Figure 9.** ML phylogeny of methanogenesis marker m12. Black circles indicate strongly supported branches (ultrafast bootstrap >=95, aLRT SH-like >=80).

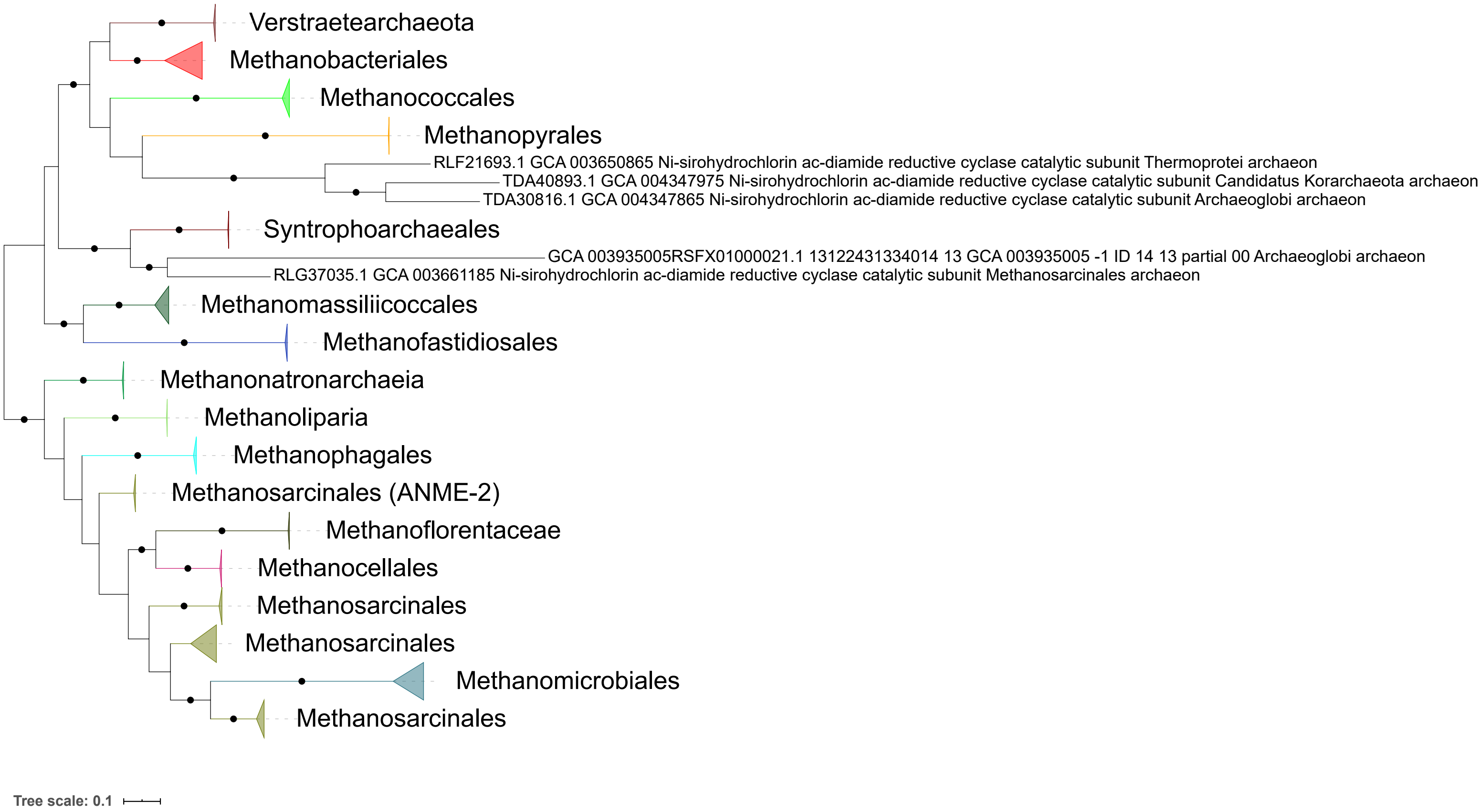
Verstraetearchaeota
Methanobacteriales
Methanococcales
Methanopyrales
RLF21693.1 GCA 003650865 Ni-sirohydrochlorin ac-diamide reductive cyclase catalytic subunit Thermoprotei archaeon
TDA40893.1 GCA 004347975 Ni-sirohydrochlorin ac-diamide reductive cyclase catalytic subunit Candidatus Korarchaeota archaeon
TDA30816.1 GCA 004347865 Ni-sirohydrochlorin ac-diamide reductive cyclase catalytic subunit Archaeoglobi archaeon
Syntrophoarchaeales
GCA 003935005RSFX01000021.1 13122431334014 13 GCA 003935005 -1 ID 14 13 partial 00 Archaeoglobi archaeon
RLG37035.1 GCA 003661185 Ni-sirohydrochlorin ac-diamide reductive cyclase catalytic subunit Methanosarcinales archaeon
Methanomassiliicoccales
Methanofastidiosales
Methanonatronarchaeia
Methanoliparia
Methanophagales
Methanosarcinales (ANME-2)
Methanoflorentaceae
Methanocellales
Methanosarcinales
Methanosarcinales
Methanomicrobiales
Methanosarcinales
Tree scale: 0.1

**Supplementary Figure 10.** ML phylogeny of methanogenesis marker m13. Black circles indicate strongly supported branches (ultrafast bootstrap >=95, aLRT SH-like >=80).

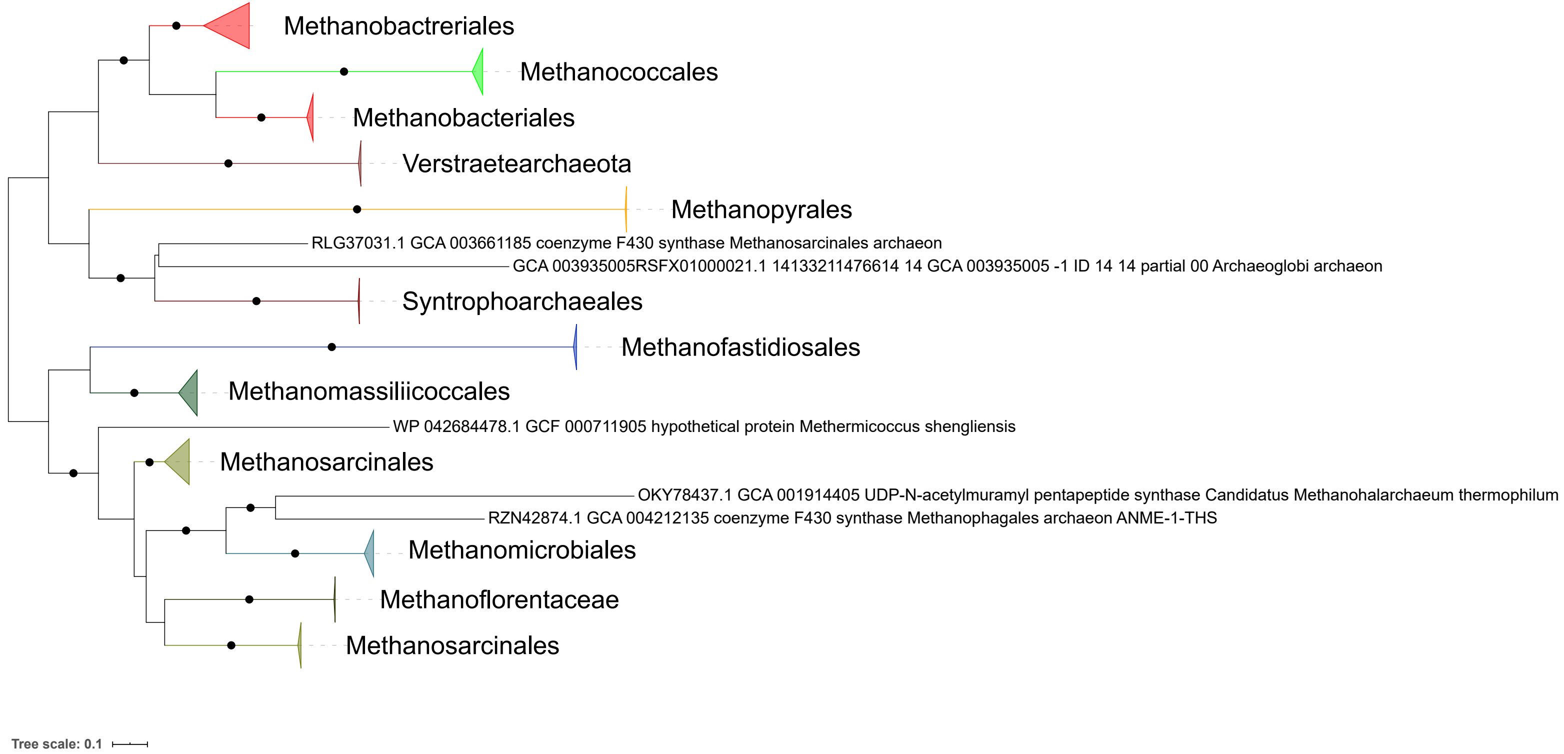
Methanobactreriales
Methanococcales
Methanobacteriales
Verstraetearchaeota
Methanopyrales
RLG37031.1 GCA 003661185 coenzyme F430 synthase Methanosarcinales archaeon
GCA 003935005RSFX01000021.1 14133211476614 14 GCA 003935005 -1 ID 14 14 partial 00 Archaeoglobi archaeon
Syntrophoarchaeales
Methanofastidiosales
Methanomassiliicoccales
WP 042684478.1 GCF 000711905 hypothetical protein Methermicoccus shengliensis
Methanosarcinales
OKY78437.1 GCA 001914405 UDP-N-acetylmuramyl pentapeptide synthase Candidatus Methanohalarchaeum thermophilum
RZN42874.1 GCA 004212135 coenzyme F430 synthase Methanophagales archaeon ANME-1-THS
Methanomicrobiales
Methanoflorentaceae
Methanosarcinales
Tree scale: 0.1

**Supplementary Figure 11.** ML phylogeny of methanogenesis marker m14. Black circles indicate strongly supported branches (ultrafast bootstrap >=95, aLRT SH-like >=80).

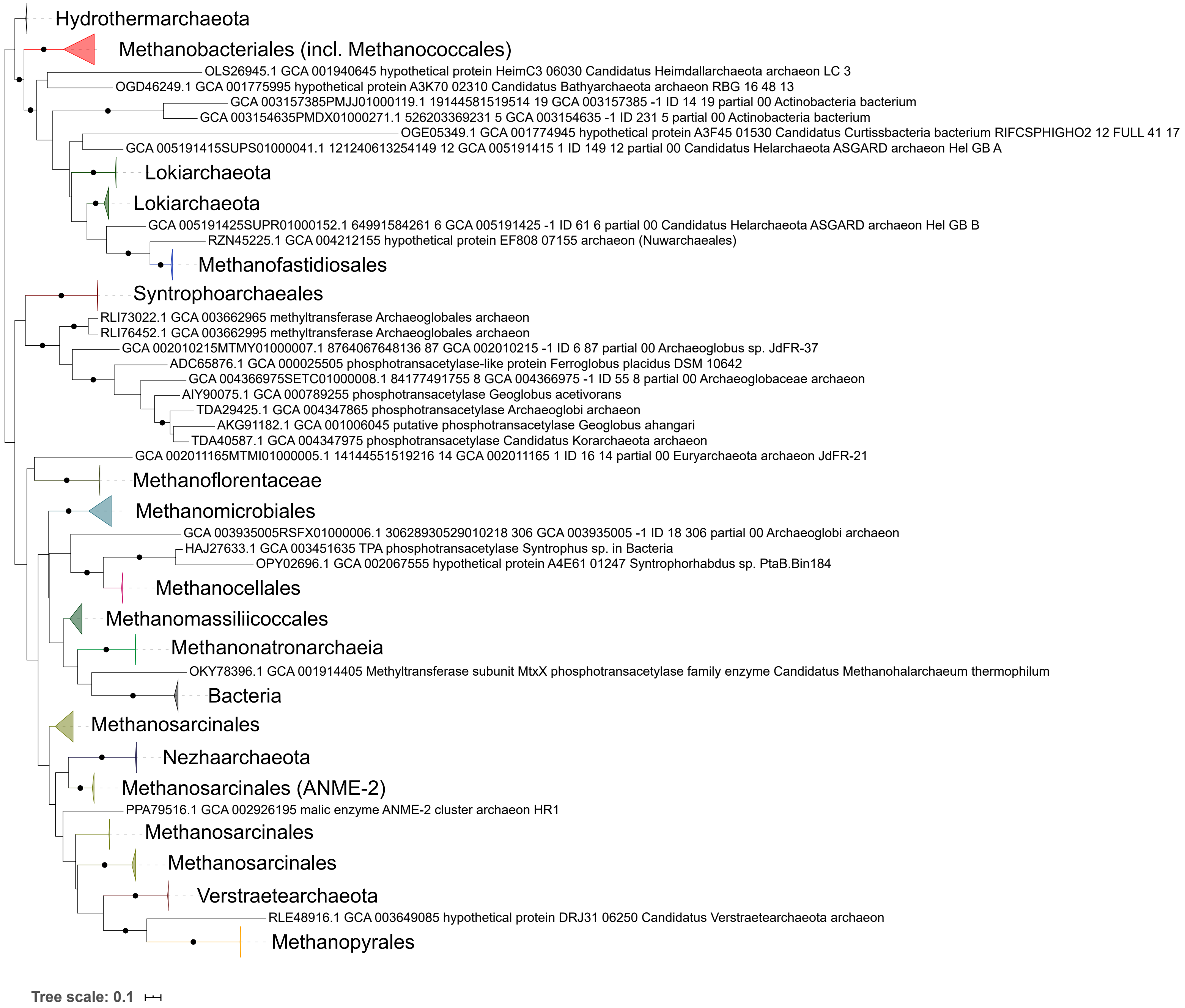

Hydrothermarchaeota
Methanobacteriales (incl. Methanococcales)
OLS26945.1 GCA 001940645 hypothetical protein HeimC3 06030 Candidatus Heimdallarchaeota archaeon LC 3
OGD46249.1 GCA 001775995 hypothetical protein A3K70 02310 Candidatus Bathyarchaeota archaeon RBG 16 48 13
GCA 003157385PMJJ01000119.1 19144581519514 19 GCA 003157385 -1 ID 14 19 partial 00 Actinobacteria bacterium
GCA 003154635PMDX01000271.1 526203369231 5 GCA 003154635 -1 ID 231 5 partial 00 Actinobacteria bacterium
OGE05349.1 GCA 001774945 hypothetical protein A3F45 01530 Candidatus Curtissbacteria bacterium RIFCSPHIGHO2 12 FULL 41 17
GCA 005191415SUPS01000041.1 121240613254149 12 GCA 005191415 1 ID 149 12 partial 00 Candidatus Helarchaeota ASGARD archaeon Hel GB A
Lokiarchaeota
Lokiarchaeota
GCA 005191425SUPR01000152.1 64991584261 6 GCA 005191425 -1 ID 61 6 partial 00 Candidatus Helarchaeota ASGARD archaeon Hel GB B
RZN45225.1 GCA 004212155 hypothetical protein EF808 07155 archaeon (Nuwarchaeales)
Methanofastidiosales
Syntrophoarchaeales
RLI73022.1 GCA 003662965 methyltransferase Archaeoglobales archaeon
RLI76452.1 GCA 003662995 methyltransferase Archaeoglobales archaeon
GCA 002010215MTMY01000007.1 8764067648136 87 GCA 002010215 -1 ID 6 87 partial 00 Archaeoglobus sp. JdFR-37
ADC65876.1 GCA 000025505 phosphotransacetylase-like protein Ferroglobus placidus DSM 10642
GCA 004366975SETC01000008.1 84177491755 8 GCA 004366975 -1 ID 55 8 partial 00 Archaeoglobaceae archaeon
AIY90075.1 GCA 000789255 phosphotransacetylase Geoglobus acetivorans
TDA29425.1 GCA 004347865 phosphotransacetylase Archaeoglobi archaeon
AKG91182.1 GCA 001006045 putative phosphotransacetylase Geoglobus ahangari
TDA40587.1 GCA 004347975 phosphotransacetylase Candidatus Korarchaeota archaeon
GCA 002011165MTMI01000005.1 14144551519216 14 GCA 002011165 1 ID 16 14 partial 00 Euryarchaeota archaeon JdFR-21
Methanoflorentaceae
Methanomicrobiales
GCA 003935005RSFX01000006.1 30628930529010218 306 GCA 003935005 -1 ID 18 306 partial 00 Archaeoglobi archaeon
HAJ27633.1 GCA 003451635 TPA phosphotransacetylase Syntrophus sp. in Bacteria
OPY02696.1 GCA 002067555 hypothetical protein A4E61 01247 Syntrophorhabdus sp. PtaB.Bin184
Methanocellales
Methanomassiliicoccales
Methanonatronarchaeia
OKY78396.1 GCA 001914405 Methyltransferase subunit MtxX phosphotransacetylase family enzyme Candidatus Methanohalarchaeum thermophilum
Bacteria
Methanosarcinales
Nezhaarchaeota
Methanosarcinales (ANME-2)
PPA79516.1 GCA 002926195 malic enzyme ANME-2 cluster archaeon HR1
Methanosarcinales
Methanosarcinales
Verstraetearchaeota
RLE48916.1 GCA 003649085 hypothetical protein DRJ31 06250 Candidatus Verstraetearchaeota archaeon
Methanopyrales
Tree scale: 0.1

**Supplementary Figure 12.** ML phylogeny of methanogenesis marker m15. Black circles indicate strongly supported branches (ultrafast bootstrap >=95, aLRT SH-like >=80).

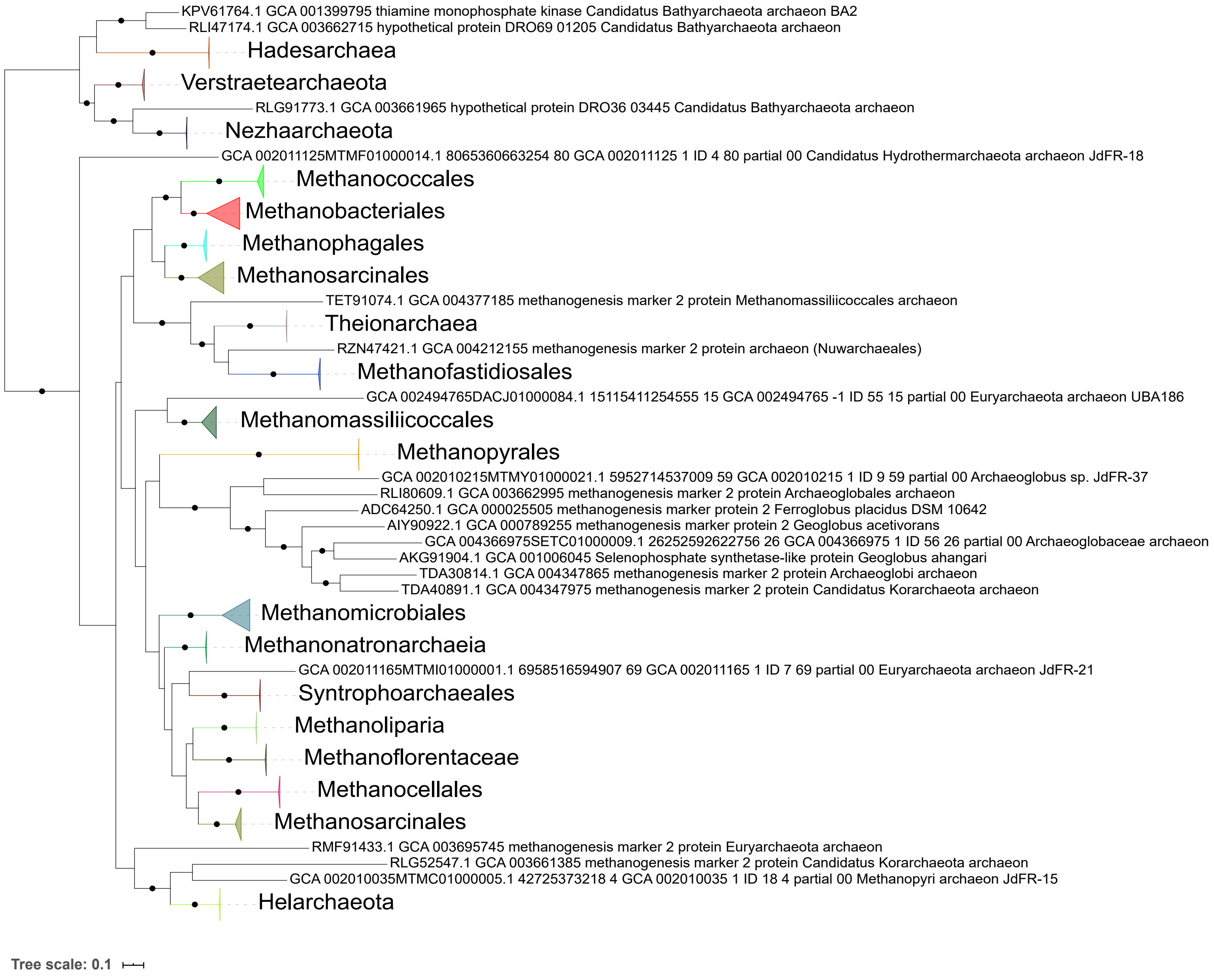

KPV61764.1 GCA 001399795 thiamine monophosphate kinase Candidatus Bathyarchaeota archaeon BA2
RLI47174.1 GCA 003662715 hypothetical protein DRO69 01205 Candidatus Bathyarchaeota archaeon
Hadesarchaea
Verstraetearchaeota
RLG91773.1 GCA 003661965 hypothetical protein DRO36 03445 Candidatus Bathyarchaeota archaeon
Nezhaarchaeota
GCA 002011125MTMF01000014.1 8065360663254 80 GCA 002011125 1 ID 4 80 partial 00 Candidatus Hydrothermarchaeota archaeon JdFR-18
Methanococcales
Methanobacteriales
Methanophagales
Methanosarcinales
TET91074.1 GCA 004377185 methanogenesis marker 2 protein Methanomassiliicoccales archaeon
Theionarchaea
RZN47421.1 GCA 004212155 methanogenesis marker 2 protein archaeon (Nuwarchaeales)
Methanofastidiosales
GCA 002494765DACJ01000084.1 15115411254555 15 GCA 002494765 -1 ID 55 15 partial 00 Euryarchaeota archaeon UBA186
Methanomassiliicoccales
Methanopyrales
GCA 002010215MTMY01000021.1 5952714537009 59 GCA 002010215 1 ID 9 59 partial 00 Archaeoglobus sp. JdFR-37
RLI80609.1 GCA 003662995 methanogenesis marker 2 protein Archaeoglobales archaeon
ADC64250.1 GCA 000025505 methanogenesis marker protein 2 Ferroglobus placidus DSM 10642
AIY90922.1 GCA 000789255 methanogenesis marker protein 2 Geoglobus acetivorans
GCA 004366975SETC01000009.1 26252592622756 26 GCA 004366975 1 ID 56 26 partial 00 Archaeoglobaceae archaeon
AKG91904.1 GCA 001006045 Selenophosphate synthetase-like protein Geoglobus ahangari
TDA30814.1 GCA 004347865 methanogenesis marker 2 protein Archaeoglobi archaeon
TDA40891.1 GCA 004347975 methanogenesis marker 2 protein Candidatus Korarchaeota archaeon
Methanomicrobiales
Methanonatronarchaeia
GCA 002011165MTMI01000001.1 6958516594907 69 GCA 002011165 1 ID 7 69 partial 00 Euryarchaeota archaeon JdFR-21
Syntrophoarchaeales
Methanoliparia
Methanoflorentaceae
Methanocellales
Methanosarcinales
RMF91433.1 GCA 003695745 methanogenesis marker 2 protein Euryarchaeota archaeon
RLG52547.1 GCA 003661385 methanogenesis marker 2 protein Candidatus Korarchaeota archaeon
GCA 002010035MTMC01000005.1 42725373218 4 GCA 002010035 1 ID 18 4 partial 00 Methanopyri archaeon JdFR-15
Helarchaeota
Tree scale: 0.1

**Supplementary Figure 13.** ML phylogeny of methanogenesis marker m16. Black circles indicate strongly supported branches (ultrafast bootstrap >=95, aLRT SH-like >=80).

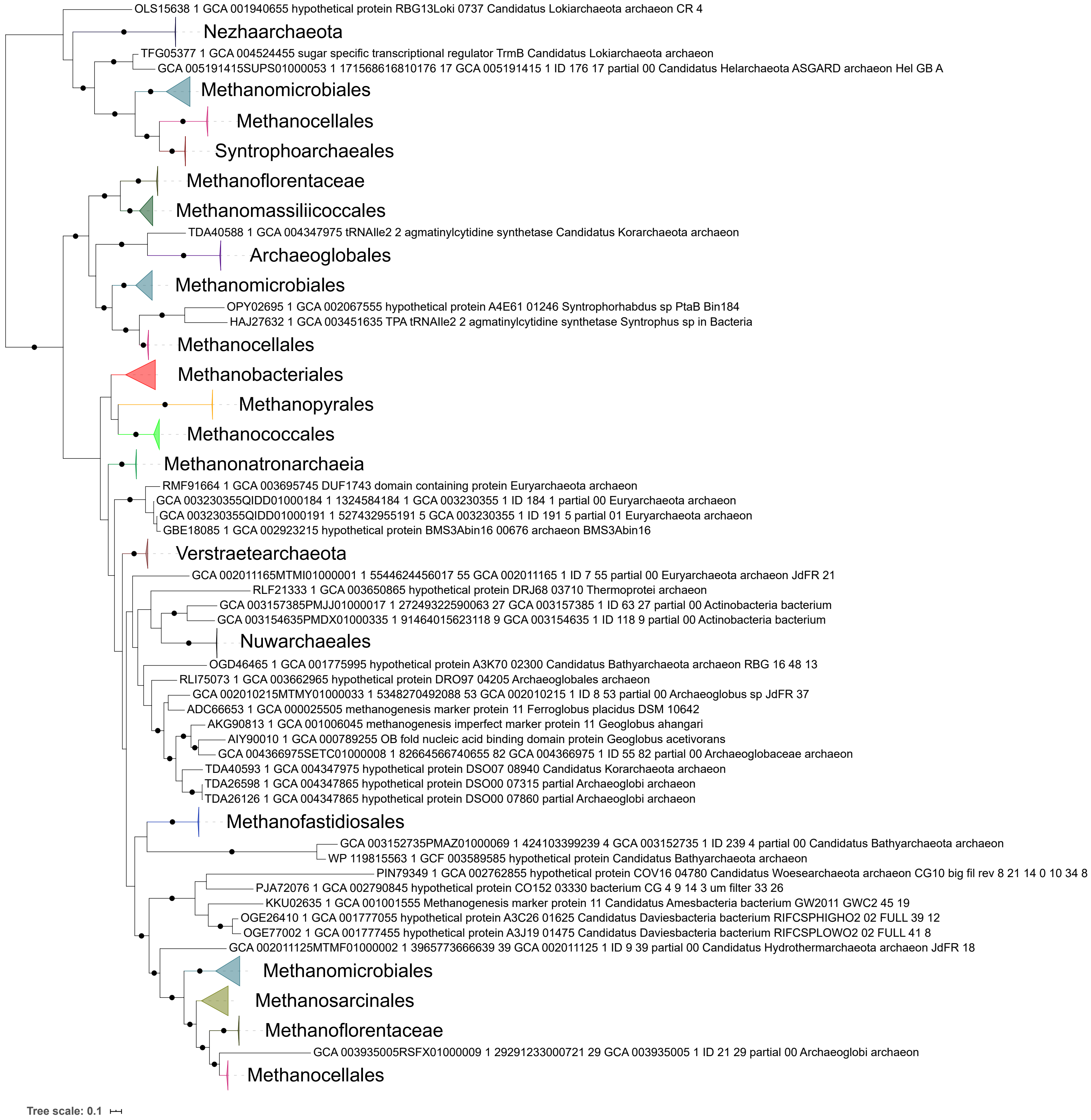
OLS15638 1 GCA 001940655 hypothetical protein RBG13Loki 0737 Candidatus Lokiarchaeota archaeon CR 4
Nezhaarchaeota
TFG05377 1 GCA 004524455 sugar specific transcriptional regulator TrmB Candidatus Lokiarchaeota archaeon
GCA 005191415SUPS01000053 1 171568616810176 17 GCA 005191415 1 ID 176 17 partial 00 Candidatus Helarchaeota ASGARD archaeon Hel GB A
Methanomicrobiales
Methanocellales
Syntrophoarchaeales
Methanoflorentaceae
Methanomassiliicoccales
TDA40588 1 GCA 004347975 tRNAIle2 2 agmatinylcytidine synthetase Candidatus Korarchaeota archaeon
Archaeoglobales
Methanomicrobiales
OPY02695 1 GCA 002067555 hypothetical protein A4E61 01246 Syntrophorhabdus sp PtaB Bin184
HAJ27632 1 GCA 003451635 TPA tRNAIle2 2 agmatinylcytidine synthetase Syntrophus sp in Bacteria
Methanocellales
Methanobacteriales
Methanopyrales
Methanococcales
Methanonatronarchaeia
RMF91664 1 GCA 003695745 DUF1743 domain containing protein Euryarchaeota archaeon
GCA 003230355QIDD01000184 1 1324584184 1 GCA 003230355 1 ID 184 1 partial 00 Euryarchaeota archaeon
GCA 003230355QIDD01000191 1 527432955191 5 GCA 003230355 1 ID 191 5 partial 01 Euryarchaeota archaeon
GBE18085 1 GCA 002923215 hypothetical protein BMS3Abin16 00676 archaeon BMS3Abin16
Verstraetearchaeota
GCA 002011165MTMI01000001 1 5544624456017 55 GCA 002011165 1 ID 7 55 partial 00 Euryarchaeota archaeon JdFR 21
RLF21333 1 GCA 003650865 hypothetical protein DRJ68 03710 Thermoprotei archaeon
GCA 003157385PMJJ01000017 1 27249322590063 27 GCA 003157385 1 ID 63 27 partial 00 Actinobacteria bacterium
GCA 003154635PMDX01000335 1 91464015623118 9 GCA 003154635 1 ID 118 9 partial 00 Actinobacteria bacterium
Nuwarchaeales
OGD46465 1 GCA 001775995 hypothetical protein A3K70 02300 Candidatus Bathyarchaeota archaeon RBG 16 48 13
RLI75073 1 GCA 003662965 hypothetical protein DRO97 04205 Archaeoglobales archaeon
GCA 002010215MTMY01000033 1 5348270492088 53 GCA 002010215 1 ID 8 53 partial 00 Archaeoglobus sp JdFR 37
ADC66653 1 GCA 000025505 methanogenesis marker protein 11 Ferroglobus placidus DSM 10642
AKG90813 1 GCA 001006045 methanogenesis imperfect marker protein 11 Geoglobus ahangari
AIY90010 1 GCA 000789255 OB fold nucleic acid binding domain protein Geoglobus acetivorans
GCA 004366975SETC01000008 1 82664566740655 82 GCA 004366975 1 ID 55 82 partial 00 Archaeoglobaceae archaeon
TDA40593 1 GCA 004347975 hypothetical protein DSO07 08940 Candidatus Korarchaeota archaeon
TDA26598 1 GCA 004347865 hypothetical protein DSO00 07315 partial Archaeoglobi archaeon
TDA26126 1 GCA 004347865 hypothetical protein DSO00 07860 partial Archaeoglobi archaeon
Methanofastidiosales
GCA 003152735PMAZ01000069 1 424103399239 4 GCA 003152735 1 ID 239 4 partial 00 Candidatus Bathyarchaeota archaeon
WP 119815563 1 GCF 003589585 hypothetical protein Candidatus Bathyarchaeota archaeon
PIN79349 1 GCA 002762855 hypothetical protein COV16 04780 Candidatus Woesearchaeota archaeon CG10 big fil rev 8 21 14 0 10 34 8
PJA72076 1 GCA 002790845 hypothetical protein CO152 03330 bacterium CG 4 9 14 3 um filter 33 26
KKU02635 1 GCA 001001555 Methanogenesis marker protein 11 Candidatus Amesbacteria bacterium GW2011 GWC2 45 19
OGE26410 1 GCA 001777055 hypothetical protein A3C26 01625 Candidatus Daviesbacteria bacterium RIFCSPHIGHO2 02 FULL 39 12
OGE77002 1 GCA 001777455 hypothetical protein A3J19 01475 Candidatus Daviesbacteria bacterium RIFCSPLOWO2 02 FULL 41 8
GCA 002011125MTMF01000002 1 3965773666639 39 GCA 002011125 1 ID 9 39 partial 00 Candidatus Hydrothermarchaeota archaeon JdFR 18
Methanomicrobiales
Methanosarcinales
Methanoflorentaceae
GCA 003935005RSFX01000009 1 29291233000721 29 GCA 003935005 1 ID 21 29 partial 00 Archaeoglobi archaeon
Methanocellales
Tree scale: 0.1

**Supplementary Figure 14.** ML phylogeny of methanogenesis marker m17. Black circles indicate strongly supported branches (ultrafast bootstrap >=95, aLRT SH-like >=80).

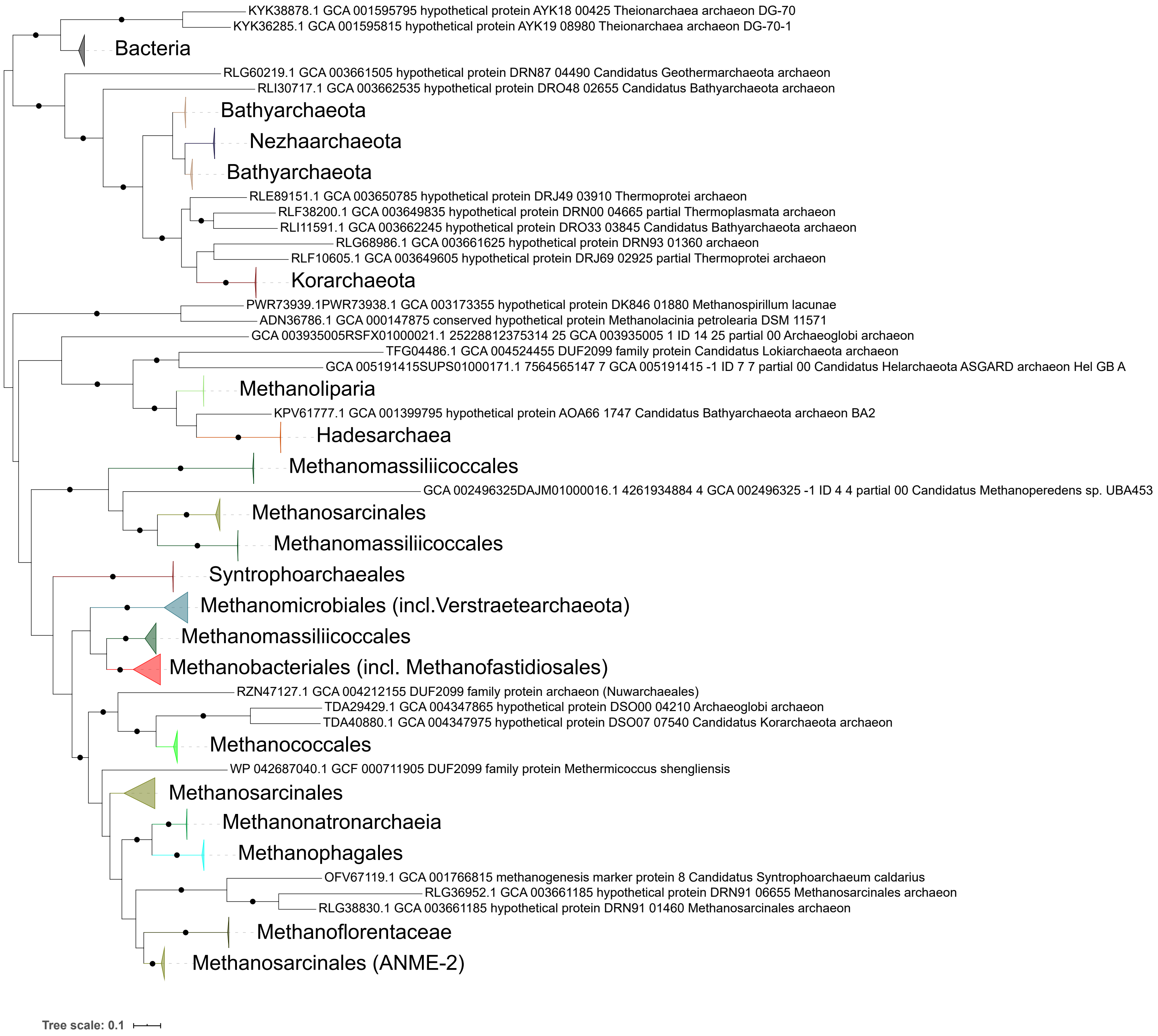

KYK38878.1 GCA 001595795 hypothetical protein AYK18 00425 Theionarchaea archaeon DG-70
KYK36285.1 GCA 001595815 hypothetical protein AYK19 08980 Theionarchaea archaeon DG-70-1
Bacteria
RLG60219.1 GCA 003661505 hypothetical protein DRN87 04490 Candidatus Geothermarchaeota archaeon
RLI30717.1 GCA 003662535 hypothetical protein DRO48 02655 Candidatus Bathyarchaeota archaeon
Bathyarchaeota
Nezhaarchaeota
Bathyarchaeota
RLE89151.1 GCA 003650785 hypothetical protein DRJ49 03910 Thermoprotei archaeon
RLF38200.1 GCA 003649835 hypothetical protein DRN00 04665 partial Thermoplasmata archaeon
RLI11591.1 GCA 003662245 hypothetical protein DRO33 03845 Candidatus Bathyarchaeota archaeon
RLG68986.1 GCA 003661625 hypothetical protein DRN93 01360 archaeon
RLF10605.1 GCA 003649605 hypothetical protein DRJ69 02925 partial Thermoprotei archaeon
Korarchaeota
PWR73939.1PWR73938.1 GCA 003173355 hypothetical protein DK846 01880 Methanospirillum lacunae
ADN36786.1 GCA 000147875 conserved hypothetical protein Methanolacinia petrolearia DSM 11571
GCA 003935005RSFX01000021.1 25228812375314 25 GCA 003935005 1 ID 14 25 partial 00 Archaeoglobi archaeon
TFG04486.1 GCA 004524455 DUF2099 family protein Candidatus Lokiarchaeota archaeon
GCA 005191415SUPS01000171.1 7564565147 7 GCA 005191415 -1 ID 7 7 partial 00 Candidatus Helarchaeota ASGARD archaeon Hel GB A
Methanoliparia
KPV61777.1 GCA 001399795 hypothetical protein AOA66 1747 Candidatus Bathyarchaeota archaeon BA2
Hadesarchaea
Methanomassiliicoccales
GCA 002496325DAJM01000016.1 4261934884 4 GCA 002496325 -1 ID 4 4 partial 00 Candidatus Methanoperedens sp. UBA453
Methanosarcinales
Methanomassiliicoccales
Syntrophoarchaeales
Methanomicrobiales (incl.Verstraetearchaeota)
Methanomassiliicoccales
Methanobacteriales (incl. Methanofastidiosales)
RZN47127.1 GCA 004212155 DUF2099 family protein archaeon (Nuwarchaeales)
TDA29429.1 GCA 004347865 hypothetical protein DSO00 04210 Archaeoglobi archaeon
TDA40880.1 GCA 004347975 hypothetical protein DSO07 07540 Candidatus Korarchaeota archaeon
Methanococcales
WP 042687040.1 GCF 000711905 DUF2099 family protein Methermicoccus shengliensis
Methanosarcinales
Methanonatronarchaeia
Methanophagales
OFV67119.1 GCA 001766815 methanogenesis marker protein 8 Candidatus Syntrophoarchaeum caldarius
RLG36952.1 GCA 003661185 hypothetical protein DRN91 06655 Methanosarcinales archaeon
RLG38830.1 GCA 003661185 hypothetical protein DRN91 01460 Methanosarcinales archaeon
Methanoflorentaceae
Methanosarcinales (ANME-2)
Tree scale: 0.1

**Supplementary Figure 15.** ML phylogeny of methanogenesis marker m18. Black circles indicate strongly supported branches (ultrafast bootstrap >=95, aLRT SH-like >=80).

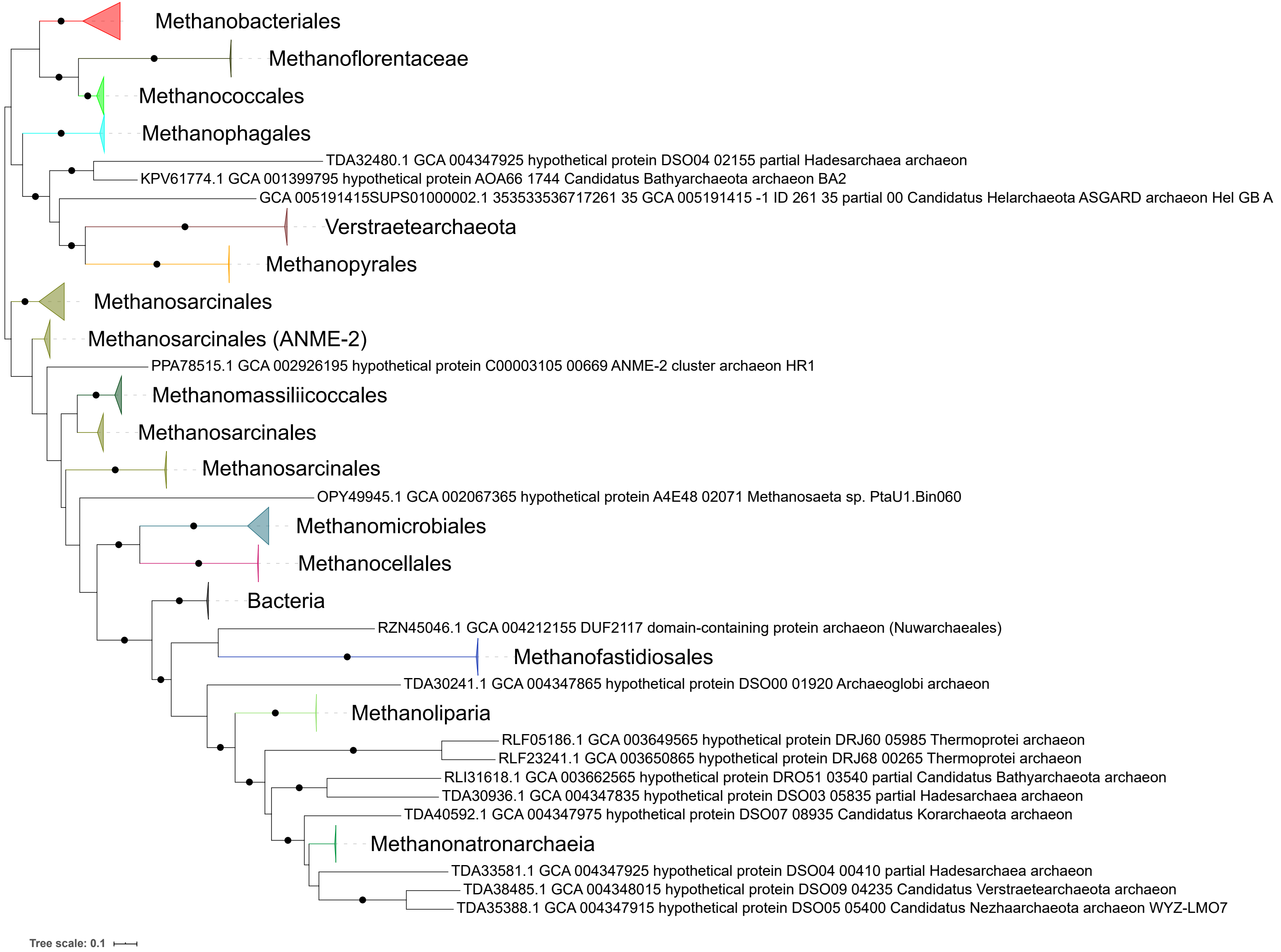
Methanobacteriales
Methanoflorentaceae
Methanococcales
Methanophagales
TDA32480.1 GCA 004347925 hypothetical protein DSO04 02155 partial Hadesarchaea archaeon
KPV61774.1 GCA 001399795 hypothetical protein AOA66 1744 Candidatus Bathyarchaeota archaeon BA2
GCA 005191415SUPS01000002.1 353533536717261 35 GCA 005191415 -1 ID 261 35 partial 00 Candidatus Helarchaeota ASGARD archaeon Hel GB A
Verstraetearchaeota
Methanopyrales
Methanosarcinales
Methanosarcinales (ANME-2)
PPA78515.1 GCA 002926195 hypothetical protein C00003105 00669 ANME-2 cluster archaeon HR1
Methanomassiliicoccales
Methanosarcinales
Methanosarcinales
OPY49945.1 GCA 002067365 hypothetical protein A4E48 02071 Methanosaeta sp. PtaU1.Bin060
Methanomicrobiales
Methanocellales
Bacteria
RZN45046.1 GCA 004212155 DUF2117 domain-containing protein archaeon (Nuwarchaeales)
Methanofastidiosales
TDA30241.1 GCA 004347865 hypothetical protein DSO00 01920 Archaeoglobi archaeon
Methanoliparia
RLF05186.1 GCA 003649565 hypothetical protein DRJ60 05985 Thermoprotei archaeon
RLF23241.1 GCA 003650865 hypothetical protein DRJ68 00265 Thermoprotei archaeon
RLI31618.1 GCA 003662565 hypothetical protein DRO51 03540 partial Candidatus Bathyarchaeota archaeon
TDA30936.1 GCA 004347835 hypothetical protein DSO03 05835 partial Hadesarchaea archaeon
TDA40592.1 GCA 004347975 hypothetical protein DSO07 08935 Candidatus Korarchaeota archaeon
Methanonatronarchaeia
TDA33581.1 GCA 004347925 hypothetical protein DSO04 00410 partial Hadesarchaea archaeon
TDA38485.1 GCA 004348015 hypothetical protein DSO09 04235 Candidatus Verstraetearchaeota archaeon
TDA35388.1 GCA 004347915 hypothetical protein DSO05 05400 Candidatus Nezhaarchaeota archaeon WYZ-LMO7
Tree scale: 0.1

**Supplementary Figure 16.** ML phylogeny of methanogenesis marker m19. Black circles indicate strongly supported branches (ultrafast bootstrap >=95, aLRT SH-like >=80).

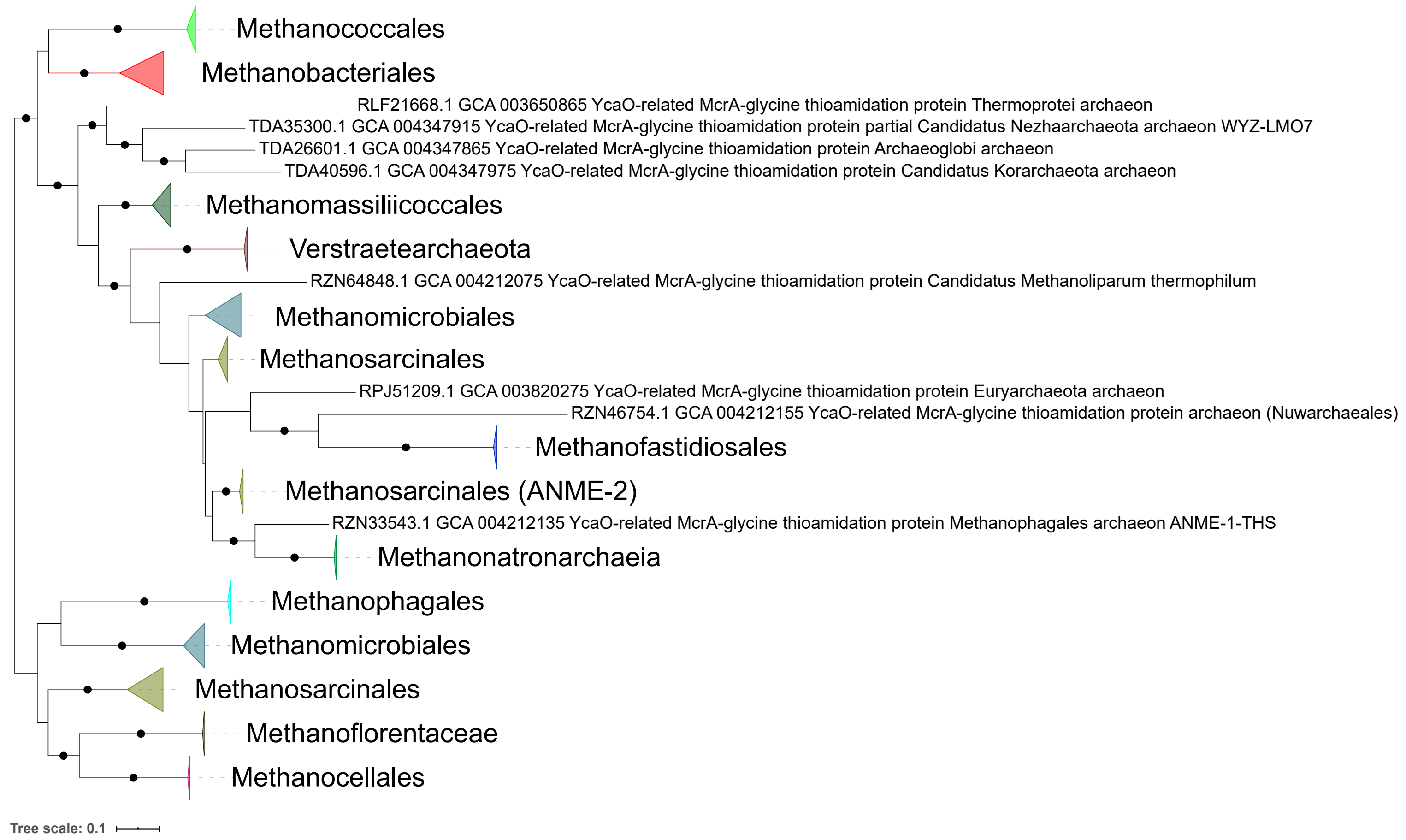
Methanococcales
Methanobacteriales
RLF21668.1 GCA 003650865 YcaO-related McrA-glycine thioamidation protein Thermoprotei archaeon
TDA35300.1 GCA 004347915 YcaO-related McrA-glycine thioamidation protein partial Candidatus Nezhaarchaeota archaeon WYZ-LMO7
TDA26601.1 GCA 004347865 YcaO-related McrA-glycine thioamidation protein Archaeoglobi archaeon
TDA40596.1 GCA 004347975 YcaO-related McrA-glycine thioamidation protein Candidatus Korarchaeota archaeon
Methanomassiliicoccales
Verstraetearchaeota
RZN64848.1 GCA 004212075 YcaO-related McrA-glycine thioamidation protein Candidatus Methanoliparum thermophilum
Methanomicrobiales
Methanosarcinales
RPJ51209.1 GCA 003820275 YcaO-related McrA-glycine thioamidation protein Euryarchaeota archaeon
RZN46754.1 GCA 004212155 YcaO-related McrA-glycine thioamidation protein archaeon (Nuwarchaeales)
Methanofastidiosales
Methanosarcinales (ANME-2)
RZN33543.1 GCA 004212135 YcaO-related McrA-glycine thioamidation protein Methanophagales archaeon ANME-1-THS
Methanonatronarchaeia
Methanophagales
Methanomicrobiales
Methanosarcinales
Methanoflorentaceae
Methanocellales
Tree scale: 0.1

**Supplementary Figure 17.** ML phylogeny of methanogenesis marker m20. Black circles indicate strongly supported branches (ultrafast bootstrap >=95, aLRT SH-like >=80).

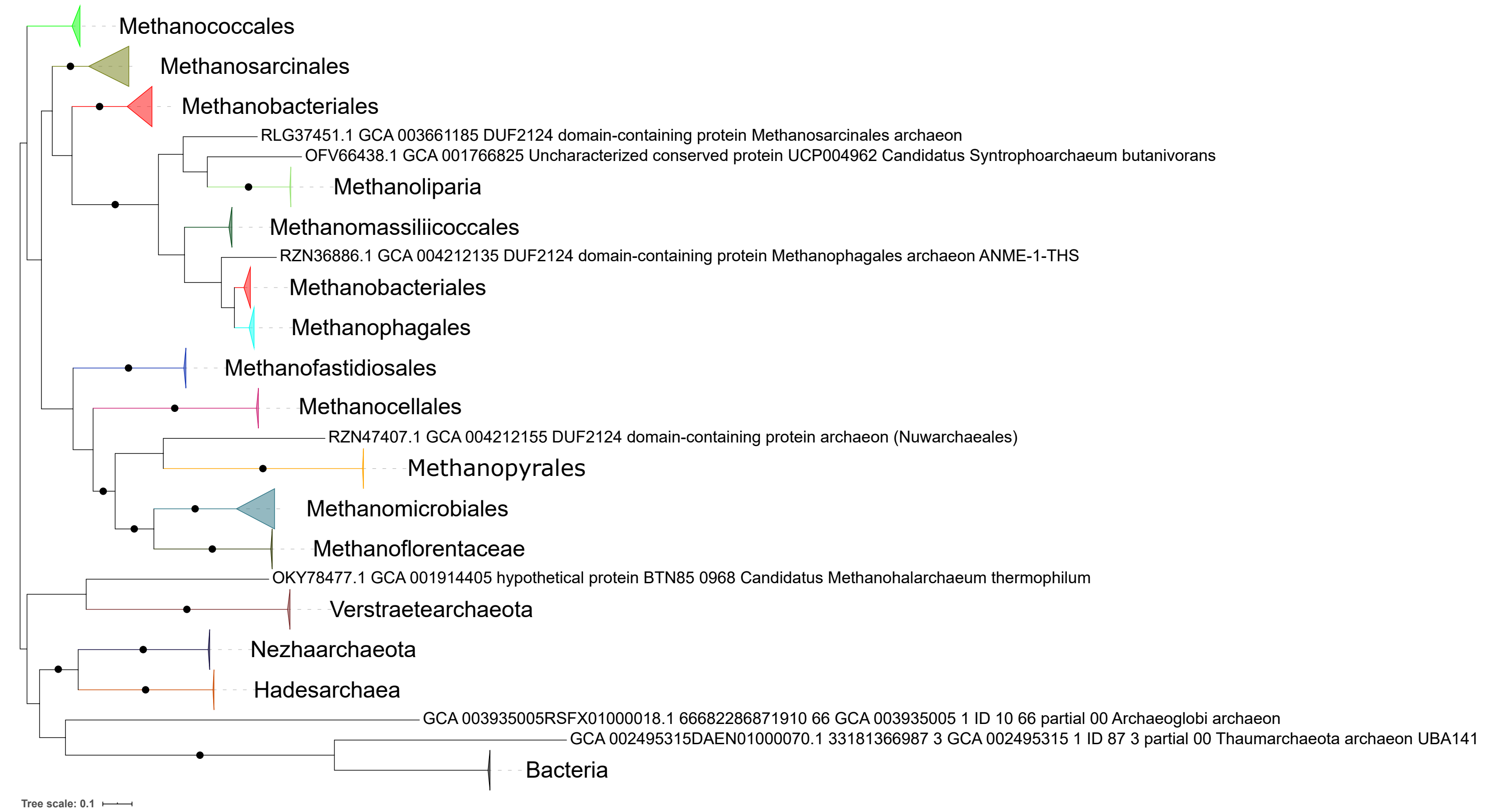
Methanococcales
Methanosarcinales
Methanobacteriales
RLG37451.1 GCA 003661185 DUF2124 domain-containing protein Methanosarcinales archaeon
OFV66438.1 GCA 001766825 Uncharacterized conserved protein UCP004962 Candidatus Syntrophoarchaeum butanivorans
Methanoliparia
Methanomassiliicoccales
RZN36886.1 GCA 004212135 DUF2124 domain-containing protein Methanophagales archaeon ANME-1-THS
Methanobacteriales
Methanophagales
Methanofastidiosales
Methanocellales
RZN47407.1 GCA 004212155 DUF2124 domain-containing protein archaeon (Nuwarchaeales)
Methanopyrales
Methanomicrobiales
Methanoflorentaceae
OKY78477.1 GCA 001914405 hypothetical protein BTN85 0968 Candidatus Methanohalarchaeum thermophilum
Verstraetearchaeota
Nezhaarchaeota
Hadesarchaea
GCA 003935005RSFX01000018.1 66682286871910 66 GCA 003935005 1 ID 10 66 partial 00 Archaeoglobi archaeon
GCA 002495315DAEN01000070.1 33181366987 3 GCA 002495315 1 ID 87 3 partial 00 Thaumarchaeota archaeon UBA141
Bacteria
Tree scale: 0.1

**Supplementary Figure 18.** ML phylogeny of methanogenesis marker m21. Black circles indicate strongly supported branches (ultrafast bootstrap >=95, aLRT SH-like >=80).

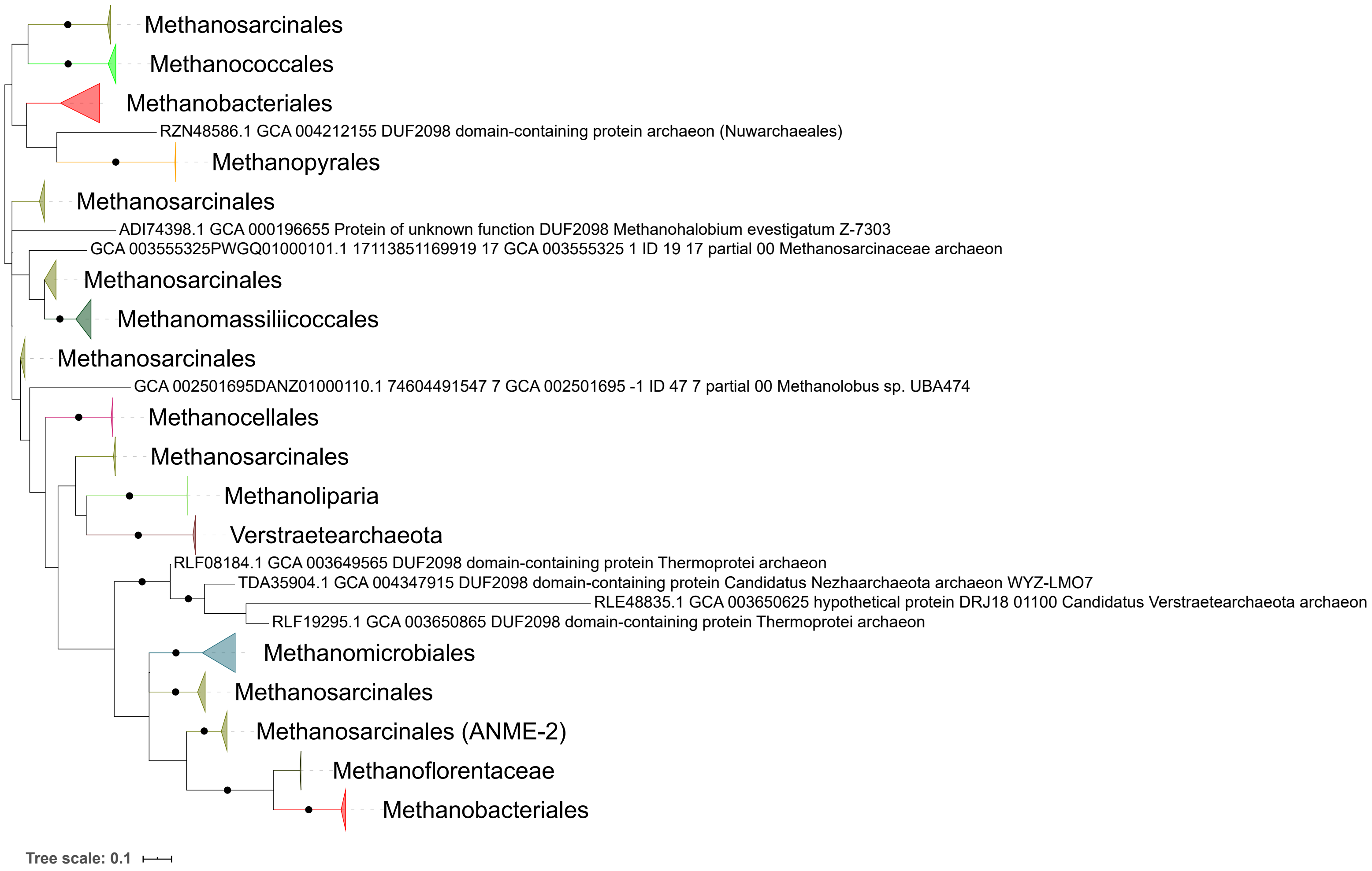
Methanosarcinales
Methanococcales
Methanobacteriales
RZN48586.1 GCA 004212155 DUF2098 domain-containing protein archaeon (Nuwarchaeales)
Methanopyrales
Methanosarcinales
ADI74398.1 GCA 000196655 Protein of unknown function DUF2098 Methanohalobium evestigatum Z-7303
GCA 003555325PWGQ01000101.1 17113851169919 17 GCA 003555325 1 ID 19 17 partial 00 Methanosarcinaceae archaeon
Methanosarcinales
Methanomassiliicoccales
Methanosarcinales
GCA 002501695DANZ01000110.1 74604491547 7 GCA 002501695 -1 ID 47 7 partial 00 Methanolobus sp. UBA474
Methanocellales
Methanosarcinales
Methanoliparia
Verstraetearchaeota
RLF08184.1 GCA 003649565 DUF2098 domain-containing protein Thermoprotei archaeon
TDA35904.1 GCA 004347915 DUF2098 domain-containing protein Candidatus Nezhaarchaeota archaeon WYZ-LMO7
RLE48835.1 GCA 003650625 hypothetical protein DRJ18 01100 Candidatus Verstraetearchaeota archaeon
RLF19295.1 GCA 003650865 DUF2098 domain-containing protein Thermoprotei archaeon
Methanomicrobiales
Methanosarcinales
Methanosarcinales (ANME-2)
Methanoflorentaceae
Methanobacteriales
Tree scale: 0.1

**Supplementary Figure 19.** ML phylogeny of methanogenesis marker m22. Black circles indicate strongly supported branches (ultrafast bootstrap >=95, aLRT SH-like >=80).

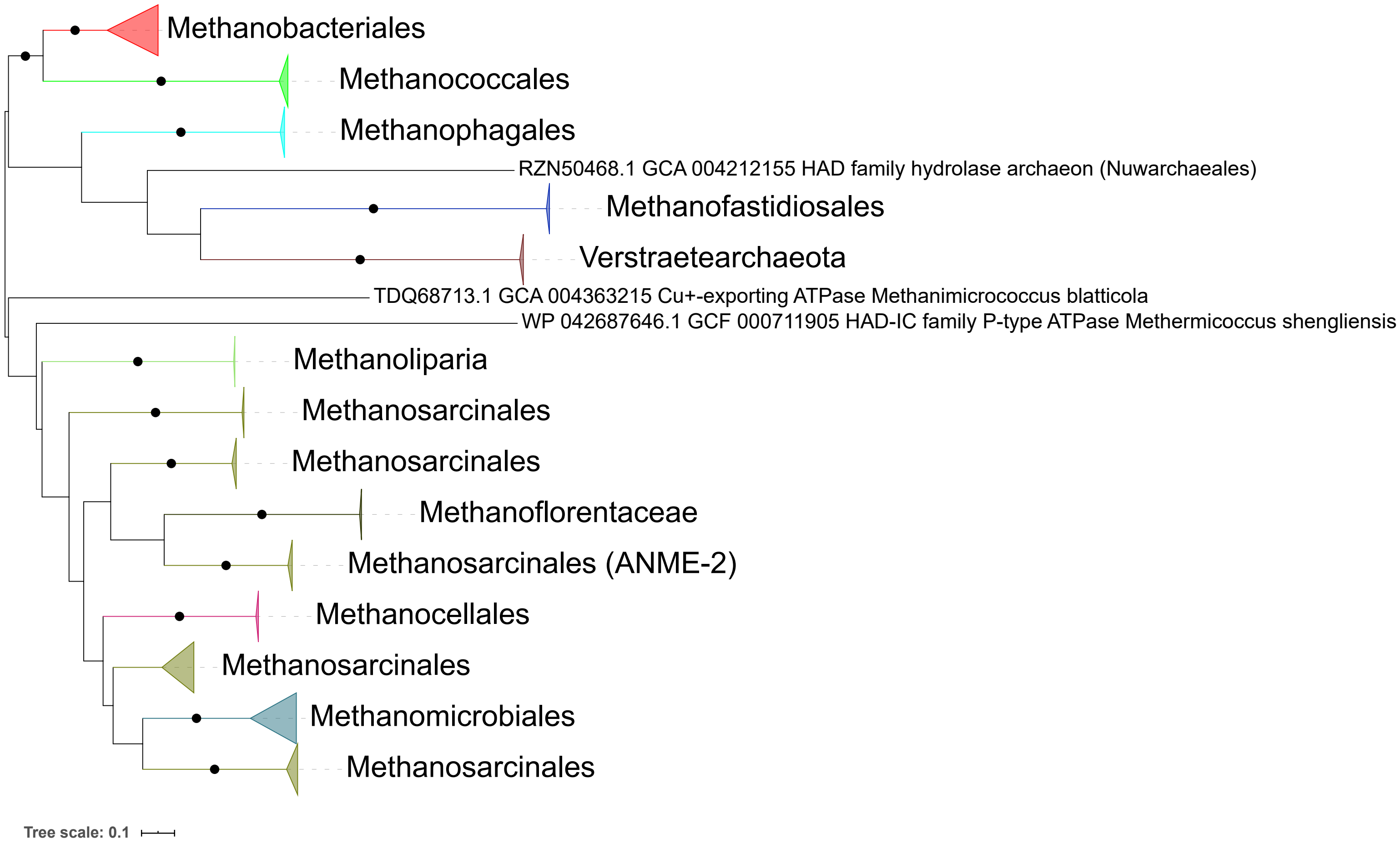
Methanobacteriales
Methanococcales
Methanophagales
RZN50468.1 GCA 004212155 HAD family hydrolase archaeon (Nuwarchaeales)
Methanofastidiosales
Verstraetearchaeota
TDQ68713.1 GCA 004363215 Cu+-exporting ATPase Methanimicrococcus blatticola
WP 042687646.1 GCF 000711905 HAD-IC family P-type ATPase Methermicoccus shengliensis
Methanoliparia
Methanosarcinales
Methanosarcinales
Methanoflorentaceae
Methanosarcinales (ANME-2)
Methanocellales
Methanosarcinales
Methanomicrobiales
Methanosarcinales
Tree scale: 0.1

**Supplementary Figure 20.** ML phylogeny of methanogenesis marker m23. Black circles indicate strongly supported branches (ultrafast bootstrap >=95, aLRT SH-like >=80).

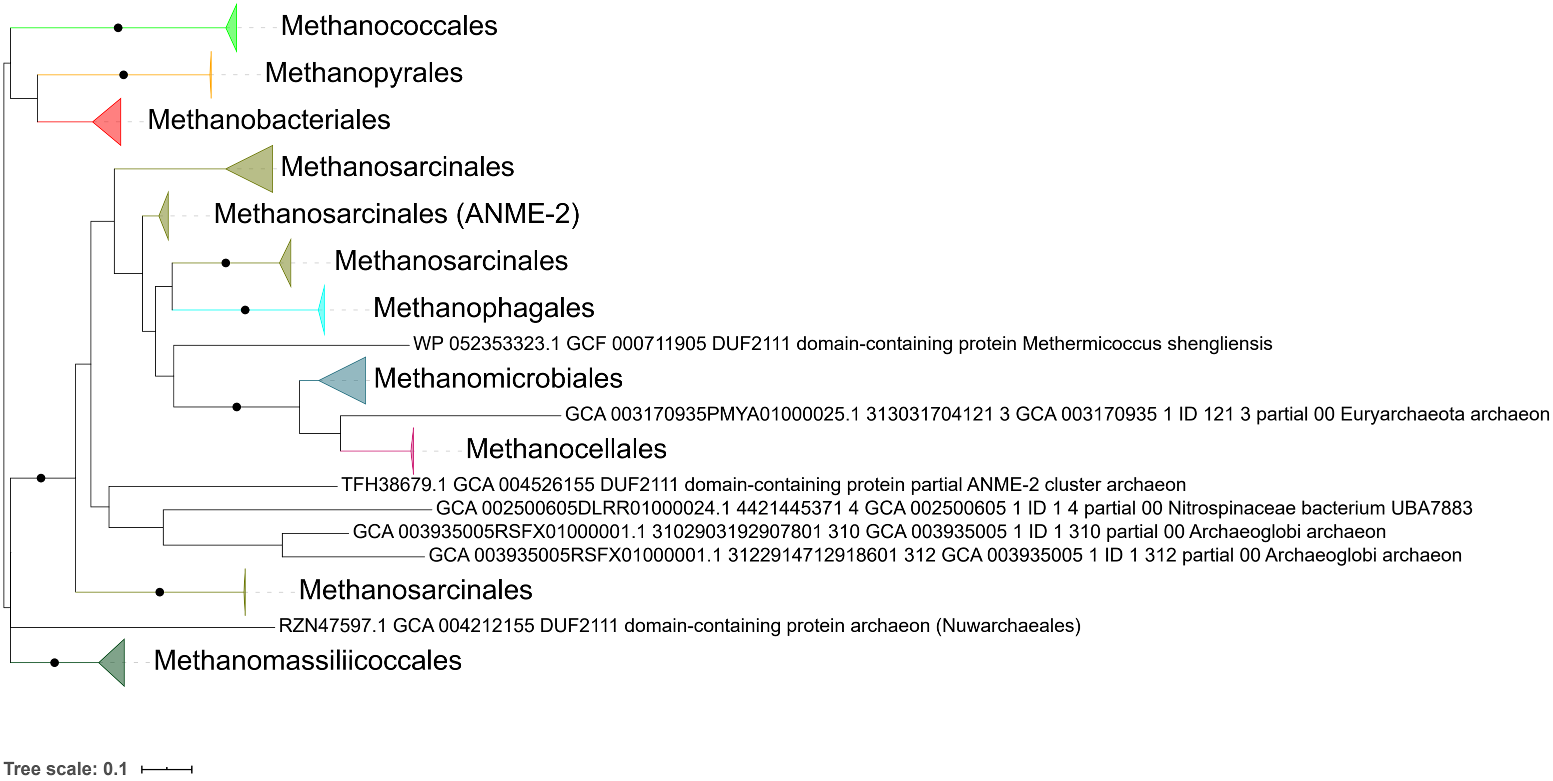

Methanococcales
Methanopyrales
Methanobacteriales
Methanosarcinales
Methanosarcinales (ANME-2)
Methanosarcinales
Methanophagales
WP 052353323.1 GCF 000711905 DUF2111 domain-containing protein Methermicoccus shengliensis
Methanomicrobiales
GCA 003170935PMYA01000025.1 313031704121 3 GCA 003170935 1 ID 121 3 partial 00 Euryarchaeota archaeon
Methanocellales
TFH38679.1 GCA 004526155 DUF2111 domain-containing protein partial ANME-2 cluster archaeon
GCA 002500605DLRR01000024.1 4421445371 4 GCA 002500605 1 ID 1 4 partial 00 Nitrospinaceae bacterium UBA7883
GCA 003935005RSFX01000001.1 3102903192907801 310 GCA 003935005 1 ID 1 310 partial 00 Archaeoglobi archaeon
GCA 003935005RSFX01000001.1 3122914712918601 312 GCA 003935005 1 ID 1 312 partial 00 Archaeoglobi archaeon
Methanosarcinales
RZN47597.1 GCA 004212155 DUF2111 domain-containing protein archaeon (Nuwarchaeales)
Methanomassiliicoccales
Tree scale: 0.1

**Supplementary Figure 21.** ML phylogeny of methanogenesis marker m24. Black circles indicate strongly supported branches (ultrafast bootstrap >=95, aLRT SH-like >=80).

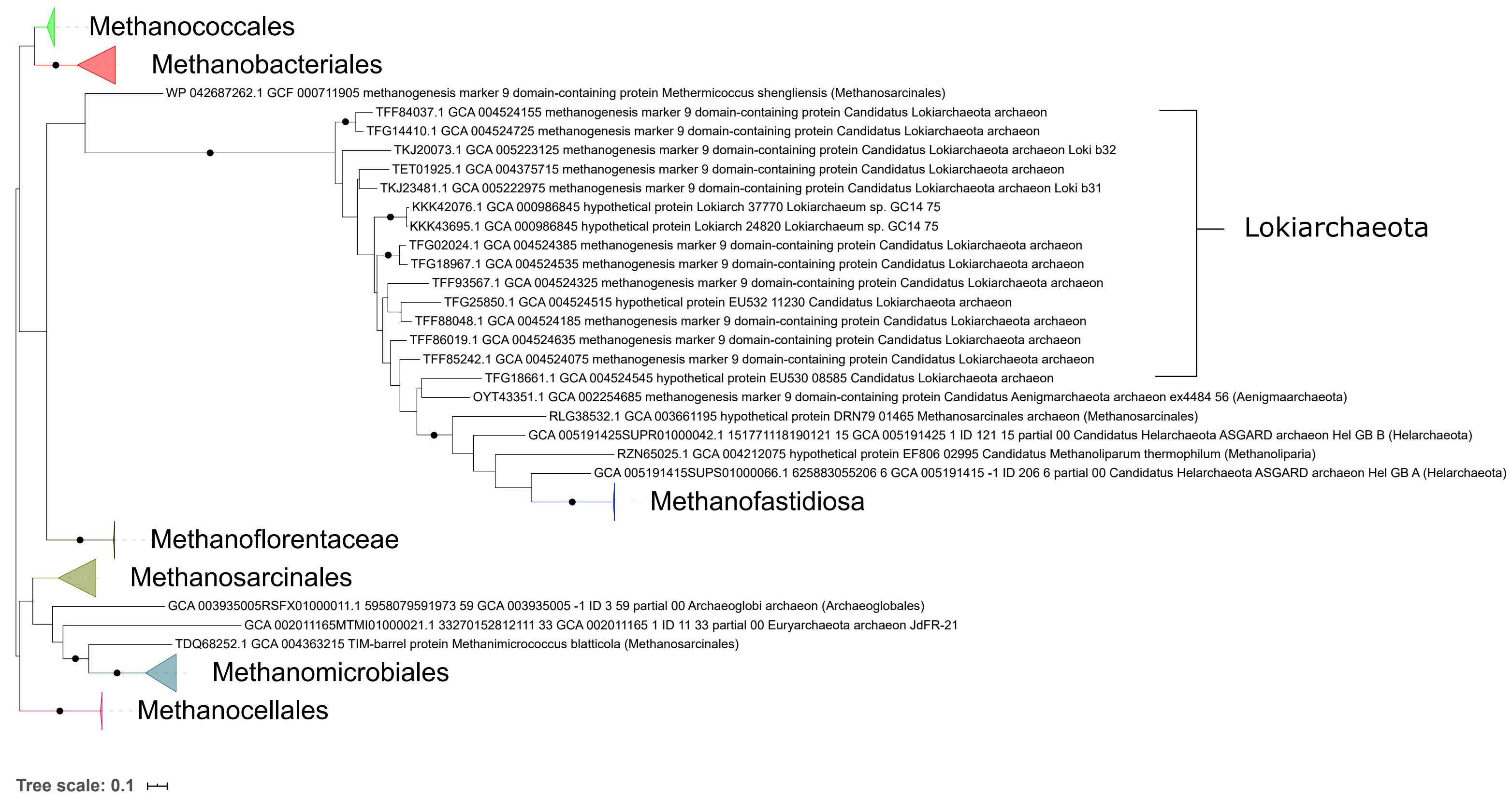
Methanococcales
Methanobacteriales
WP 042687262.1 GCF 000711905 methanogenesis marker 9 domain-containing protein Methermicoccus shengliensis (Methanosarcinales)
TFF84037.1 GCA 004524155 methanogenesis marker 9 domain-containing protein Candidatus Lokiarchaeota archaeon
TFG14410.1 GCA 004524725 methanogenesis marker 9 domain-containing protein Candidatus Lokiarchaeota archaeon
TKJ20073.1 GCA 005223125 methanogenesis marker 9 domain-containing protein Candidatus Lokiarchaeota archaeon Loki b32
TET01925.1 GCA 004375715 methanogenesis marker 9 domain-containing protein Candidatus Lokiarchaeota archaeon
TKJ23481.1 GCA 005222975 methanogenesis marker 9 domain-containing protein Candidatus Lokiarchaeota archaeon Loki b31
KKK42076.1 GCA 000986845 hypothetical protein Lokiarch 37770 Lokiarchaeum sp. GC14 75
KKK43695.1 GCA 000986845 hypothetical protein Lokiarch 24820 Lokiarchaeum sp. GC14 75
TFG02024.1 GCA 004524385 methanogenesis marker 9 domain-containing protein Candidatus Lokiarchaeota archaeon
TFG18967.1 GCA 004524535 methanogenesis marker 9 domain-containing protein Candidatus Lokiarchaeota archaeon
TFF93567.1 GCA 004524325 methanogenesis marker 9 domain-containing protein Candidatus Lokiarchaeota archaeon
TFG25850.1 GCA 004524515 hypothetical protein EU532 11230 Candidatus Lokiarchaeota archaeon
TFF88048.1 GCA 004524185 methanogenesis marker 9 domain-containing protein Candidatus Lokiarchaeota archaeon
TFF86019.1 GCA 004524635 methanogenesis marker 9 domain-containing protein Candidatus Lokiarchaeota archaeon
TFF85242.1 GCA 004524075 methanogenesis marker 9 domain-containing protein Candidatus Lokiarchaeota archaeon
TFG18661.1 GCA 004524545 hypothetical protein EU530 08585 Candidatus Lokiarchaeota archaeon
Lokiarchaeota
OYT43351.1 GCA 002254685 methanogenesis marker 9 domain-containing protein Candidatus Aenigmarchaeota archaeon ex4484 56 (Aenigmaarchaeota)
RLG38532.1 GCA 003661195 hypothetical protein DRN79 01465 Methanosarcinales archaeon (Methanosarcinales)
GCA 005191425SUPR01000042.1 151771118190121 15 GCA 005191425 1 ID 121 15 partial 00 Candidatus Helarchaeota ASGARD archaeon Hel GB B (Helarchaeota)
RZN65025.1 GCA 004212075 hypothetical protein EF806 02995 Candidatus Methanoliparum thermophilum (Methanoliparia)
GCA 005191415SUPS01000066.1 625883055206 6 GCA 005191415 -1 ID 206 6 partial 00 Candidatus Helarchaeota ASGARD archaeon Hel GB A (Helarchaeota)
Methanofastidiosa
Methanoflorentaceae
Methanosarcinales
GCA 003935005RSFX01000011.1 5958079591973 59 GCA 003935005 -1 ID 3 59 partial 00 Archaeoglobi archaeon (Archaeoglobales)
GCA 002011165MTMI01000021.1 33270152812111 33 GCA 002011165 1 ID 11 33 partial 00 Euryarchaeota archaeon JdFR-21
TDQ68252.1 GCA 004363215 TIM-barrel protein Methanimicrococcus blatticola (Methanosarcinales)
Methanomicrobiales
Methanocellales
Tree scale: 0.1

**Supplementary Figure 22.** ML phylogeny of methanogenesis marker m25. Black circles indicate strongly supported branches (ultrafast bootstrap >=95, aLRT SH-like >=80).

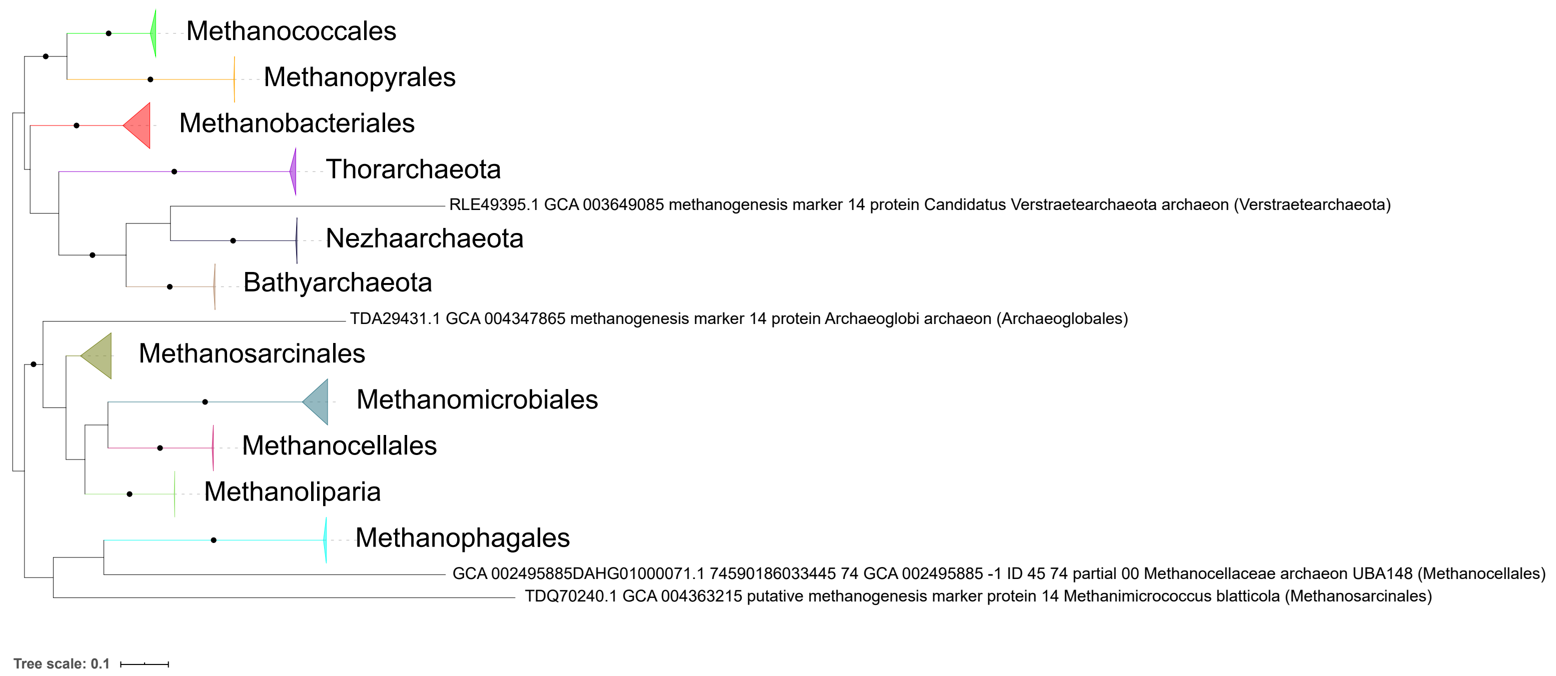
Methanococcales
Methanopyrales
Methanobacteriales
Thorarchaeota
RLE49395.1 GCA 003649085 methanogenesis marker 14 protein Candidatus Verstraetearchaeota archaeon (Verstraetearchaeota)
Nezhaarchaeota
Bathyarchaeota
TDA29431.1 GCA 004347865 methanogenesis marker 14 protein Archaeoglobi archaeon (Archaeoglobales)
Methanosarcinales
Methanomicrobiales
Methanocellales
Methanoliparia
Methanophagales
GCA 002495885DAHG01000071.1 74590186033445 74 GCA 002495885 -1 ID 45 74 partial 00 Methanocellaceae archaeon UBA148 (Methanocellales)
TDQ70240.1 GCA 004363215 putative methanogenesis marker protein 14 Methanimicrococcus blatticola (Methanosarcinales)
Tree scale: 0.1

**Supplementary Figure 23.** ML phylogeny of methanogenesis marker m26. Black circles indicate strongly supported branches (ultrafast bootstrap >=95, aLRT SH-like >=80).

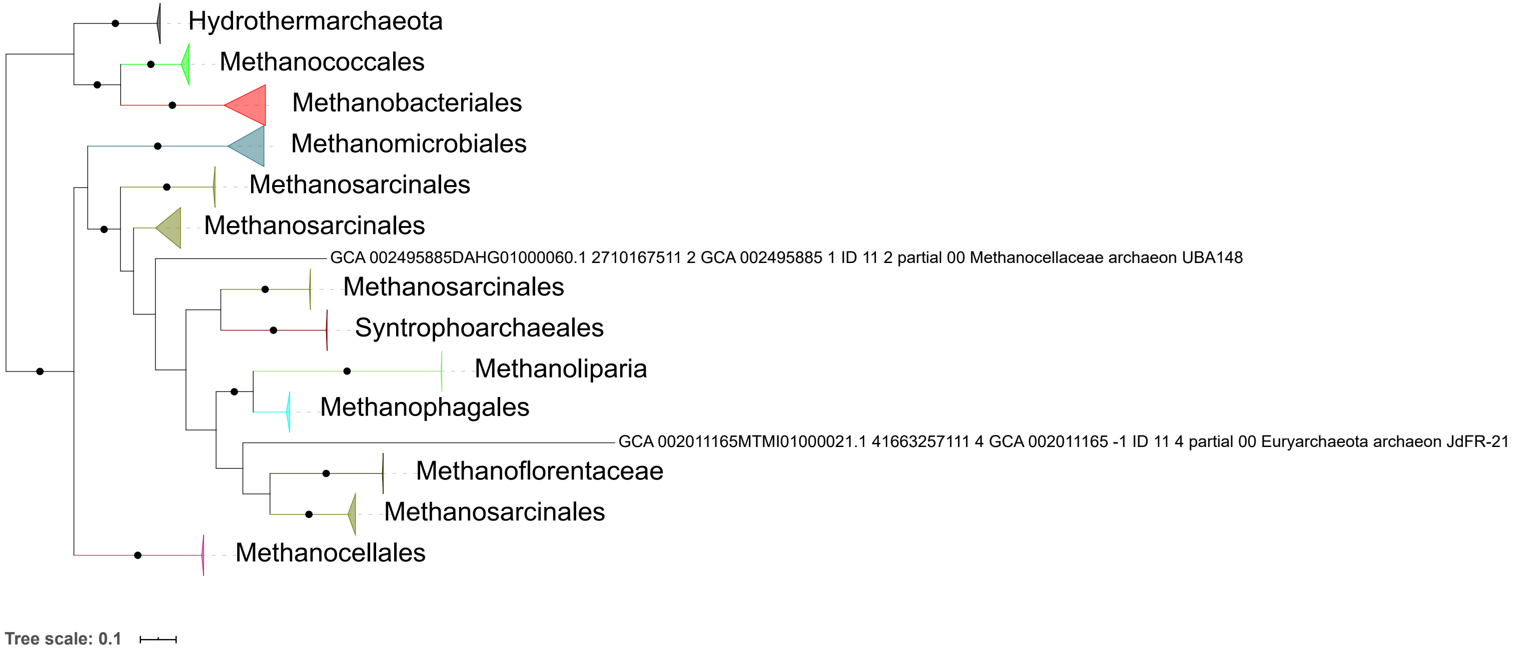
Hydrothermarchaeota
Methanococcales
Methanobacteriales
Methanomicrobiales
Methanosarcinales
Methanosarcinales
GCA 002495885DAHG01000060.1 2710167511 2 GCA 002495885 1 ID 11 2 partial 00 Methanocellaceae archaeon UBA148
Methanosarcinales
Syntrophoarchaeales
Methanoliparia
Methanophagales
GCA 002011165MTMI01000021.1 41663257111 4 GCA 002011165 -1 ID 11 4 partial 00 Euryarchaeota archaeon JdFR-21
Methanoflorentaceae
Methanosarcinales
Methanocellales
Tree scale: 0.1

**Supplementary Figure 24.** ML phylogeny of methanogenesis marker m32. Black circles indicate strongly supported branches (ultrafast bootstrap >=95, aLRT SH-like >=80).

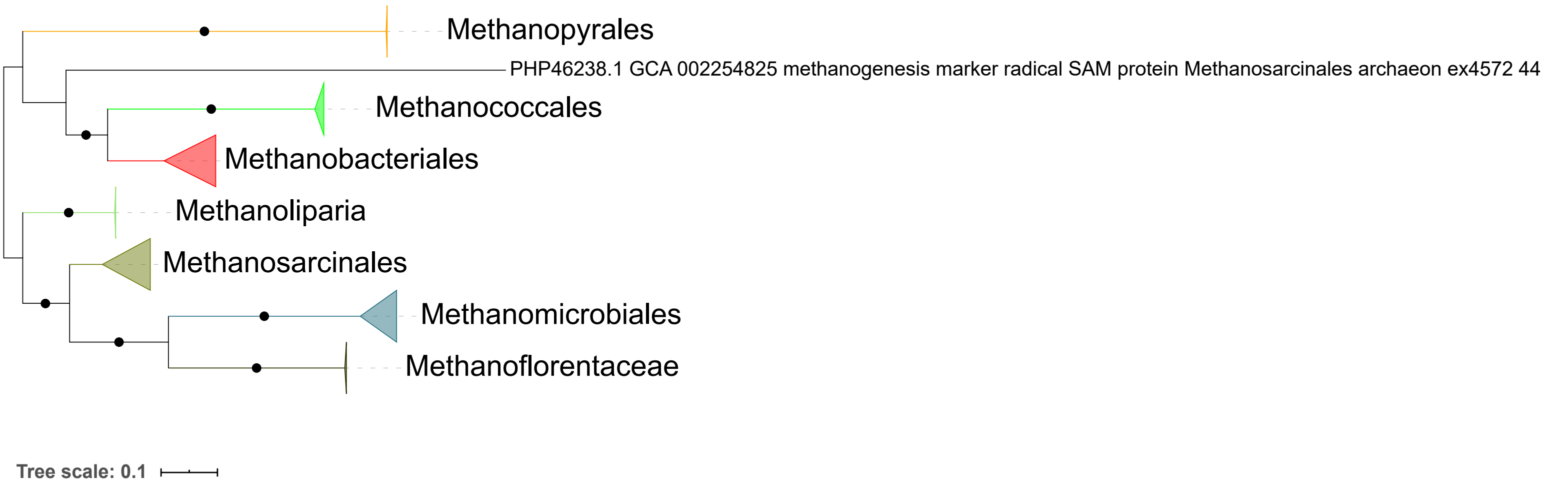
Methanopyrales
PHP46238.1 GCA 002254825 methanogenesis marker radical SAM protein Methanosarcinales archaeon ex4572 44
Methanococcales
Methanobacteriales
Methanoliparia
Methanosarcinales
Methanomicrobiales
Methanoflorentaceae
Tree scale: 0.1

**Supplementary Figure 25.** ML phylogeny of methanogenesis marker m33. Black circles indicate strongly supported branches (ultrafast bootstrap >=95, aLRT SH-like >=80).

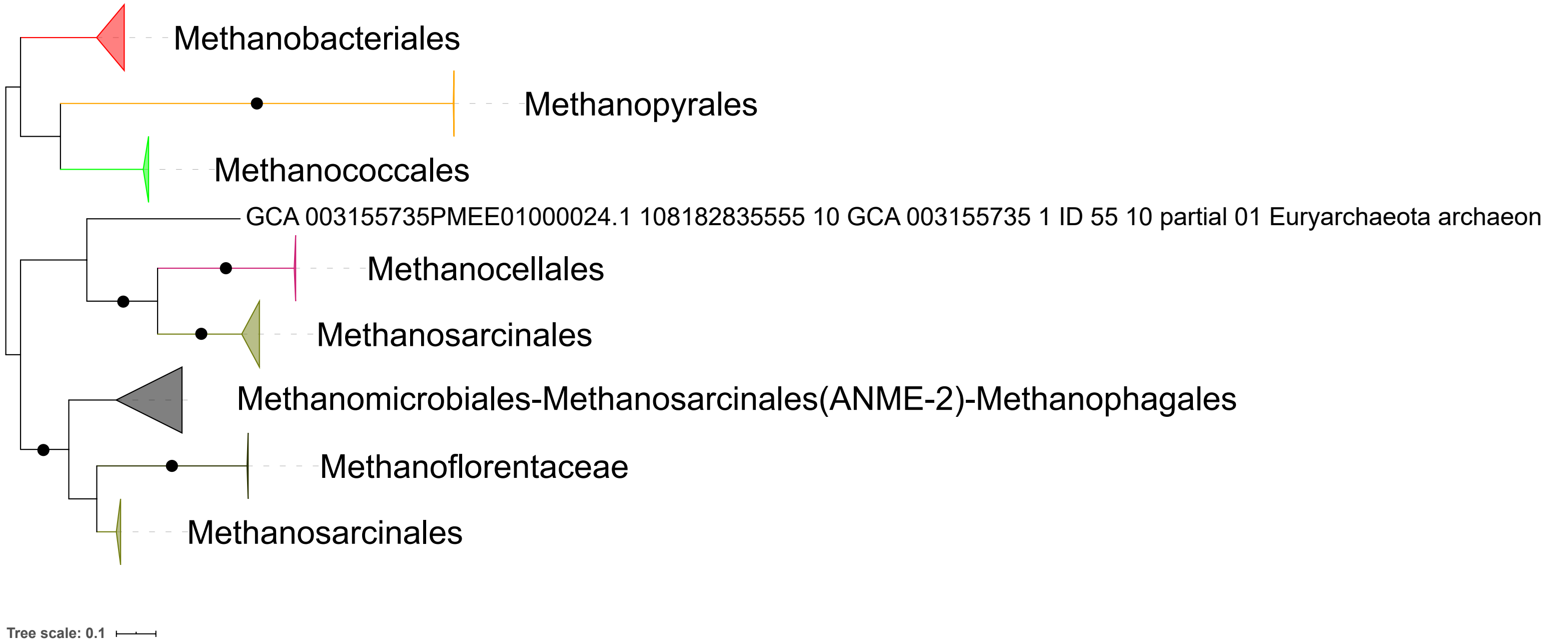
Methanobacteriales
Methanopyrales
Methanococcales
GCA 003155735PMEE01000024.1 108182835555 10 GCA 003155735 1 ID 55 10 partial 01 Euryarchaeota archaeon
Methanocellales
Methanosarcinales
Methanomicrobiales-Methanosarcinales(ANME-2)-Methanophagales
Methanoflorentaceae
Methanosarcinales
Tree scale: 0.1

**Supplementary Figure 26.** ML phylogeny of methanogenesis marker m34. Black circles indicate strongly supported branches (ultrafast bootstrap >=95, aLRT SH-like >=80).

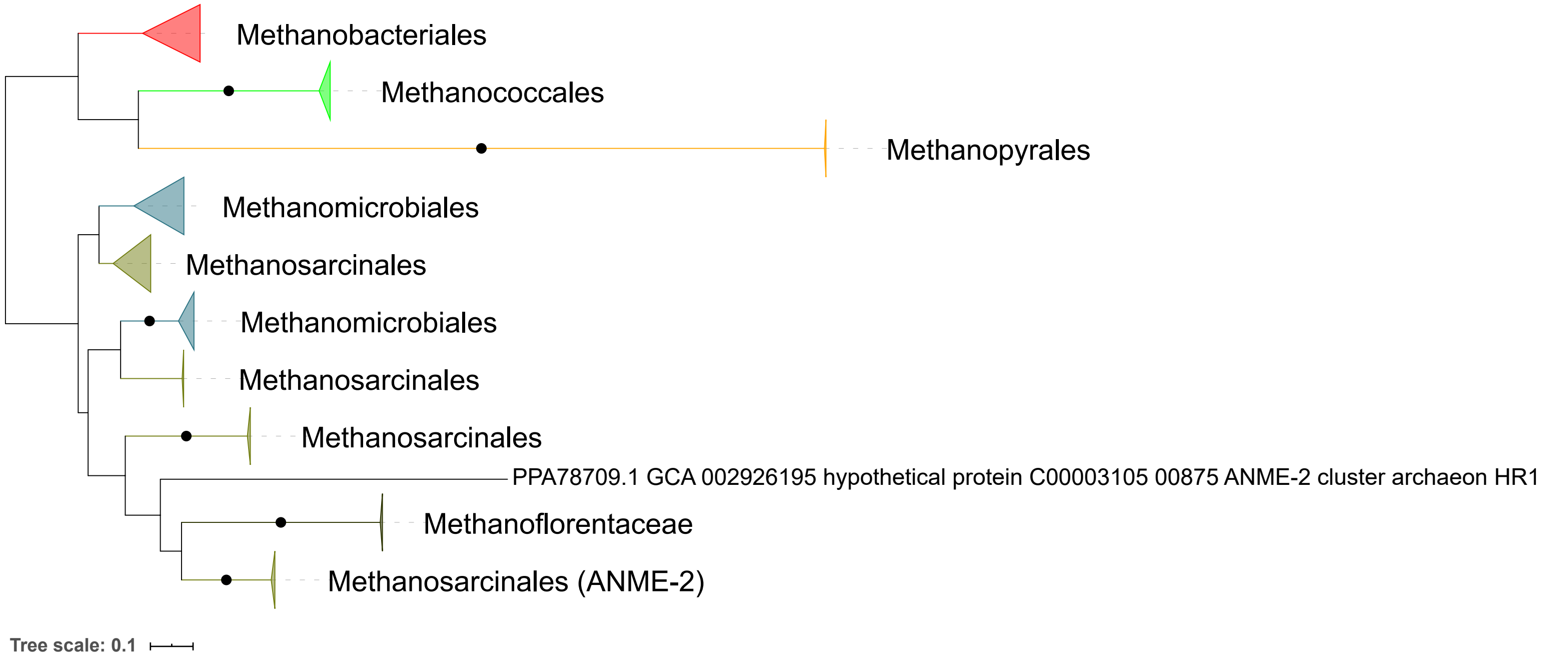
Methanobacteriales
Methanococcales
Methanopyrales
Methanomicrobiales
Methanosarcinales
Methanomicrobiales
Methanosarcinales
Methanosarcinales
PPA78709.1 GCA 002926195 hypothetical protein C00003105 00875 ANME-2 cluster archaeon HR1
Methanoflorentaceae
Methanosarcinales (ANME-2)
Tree scale: 0.1

**Supplementary Figure 27.** ML phylogeny of methanogenesis marker m35. Black circles indicate strongly supported branches (ultrafast bootstrap >=95, aLRT SH-like >=80).

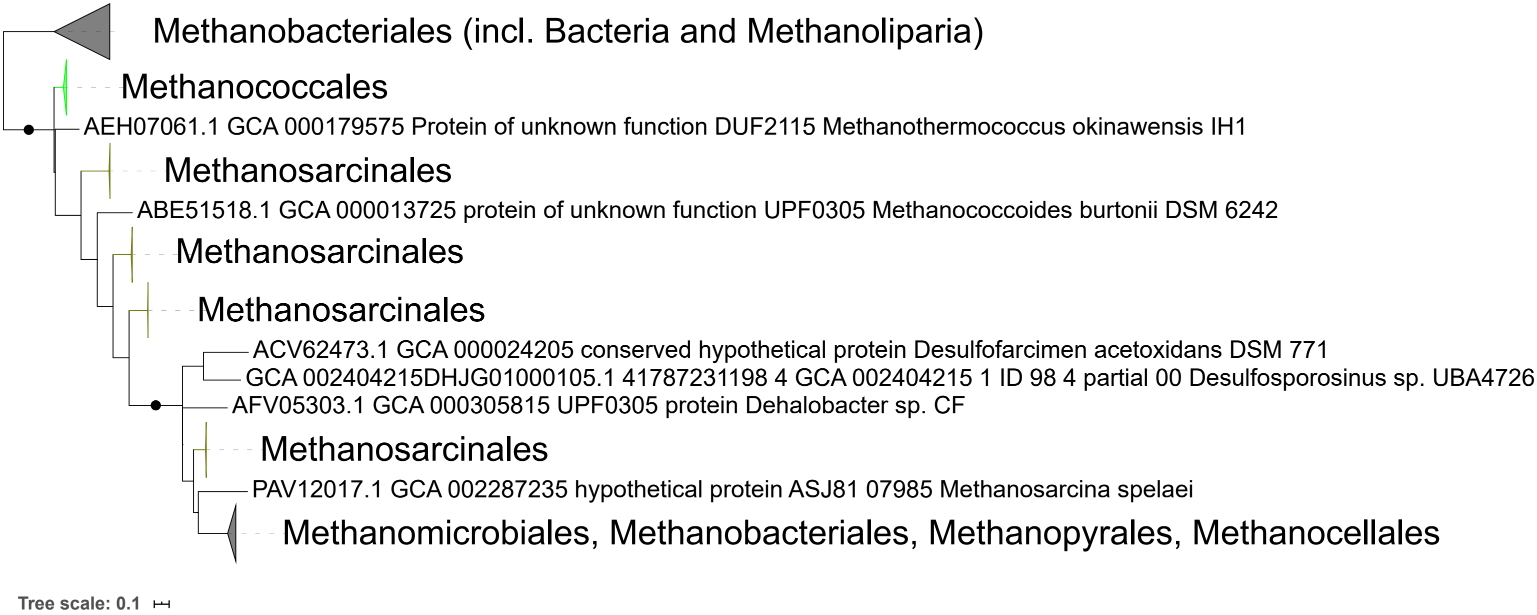
Methanobacteriales (incl. Bacteria and Methanoliparia)
Methanococcales
AEH07061.1 GCA 000179575 Protein of unknown function DUF2115 Methanothermococcus okinawensis IH1
Methanosarcinales
ABE51518.1 GCA 000013725 protein of unknown function UPF0305 Methanococcoides burtonii DSM 6242
Methanosarcinales
Methanosarcinales
ACV62473.1 GCA 000024205 conserved hypothetical protein Desulfofarcimen acetoxidans DSM 771
GCA 002404215DHJG01000105.1 41787231198 4 GCA 002404215 1 ID 98 4 partial 00 Desulfosporosinus sp. UBA4726
AFV05303.1 GCA 000305815 UPF0305 protein Dehalobacter sp. CF
Methanosarcinales
PAV12017.1 GCA 002287235 hypothetical protein ASJ81 07985 Methanosarcina spelaei
Methanomicrobiales, Methanobacteriales, Methanopyrales, Methanocellales
Tree scale: 0.1

**Supplementary Figure 28.** ML phylogeny of methanogenesis marker m36. Black circles indicate strongly supported branches (ultrafast bootstrap >=95, aLRT SH-like >=80).

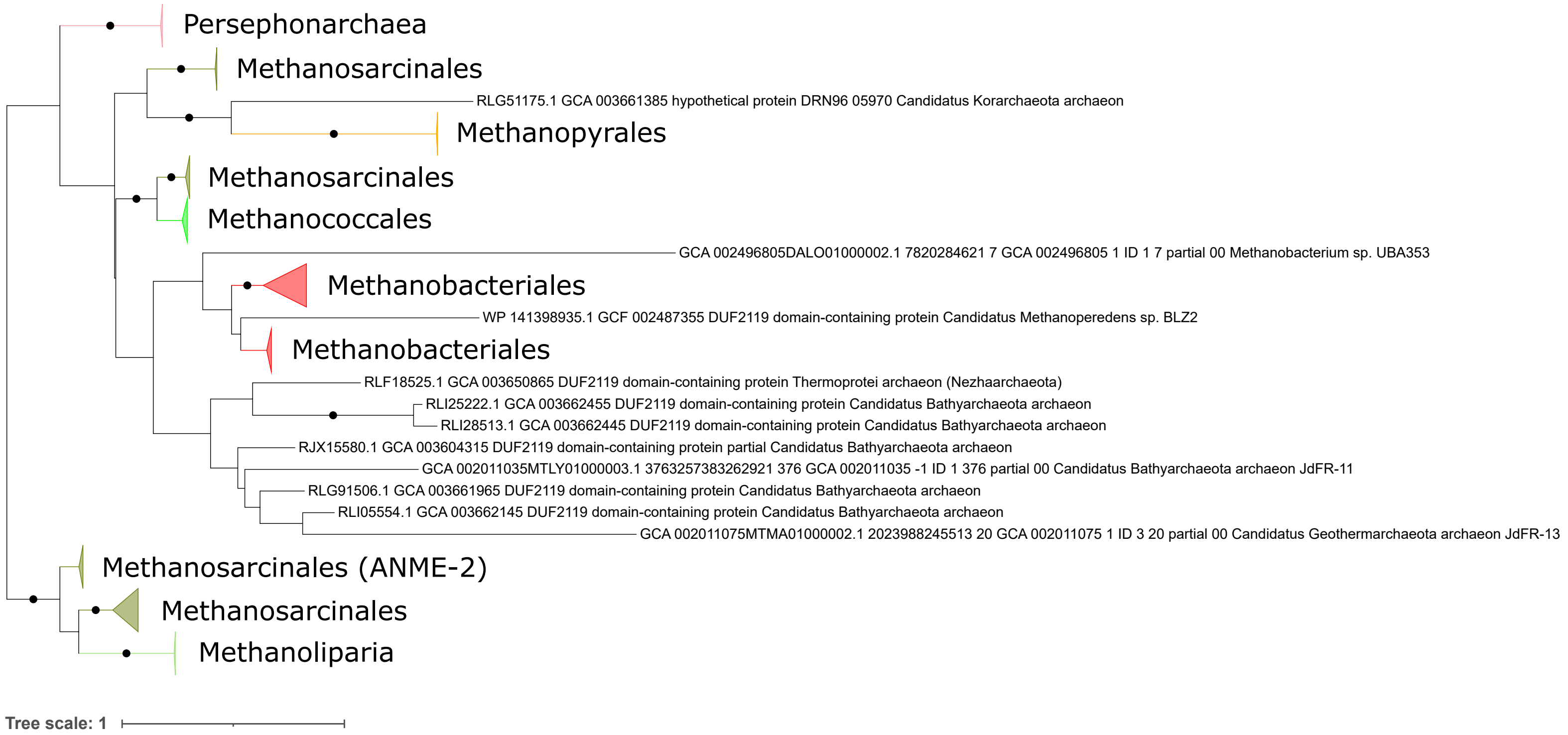
Persephonarchaea
Methanosarcinales
RLG51175.1 GCA 003661385 hypothetical protein DRN96 05970 Candidatus Korarchaeota archaeon
Methanopyrales
Methanosarcinales
Methanococcales
GCA 002496805DALO01000002.1 7820284621 7 GCA 002496805 1 ID 1 7 partial 00 Methanobacterium sp. UBA353
Methanobacteriales
WP 141398935.1 GCF 002487355 DUF2119 domain-containing protein Candidatus Methanoperedens sp. BLZ2
Methanobacteriales
RLF18525.1 GCA 003650865 DUF2119 domain-containing protein Thermoprotei archaeon (Nezhaarchaeota)
RLI25222.1 GCA 003662455 DUF2119 domain-containing protein Candidatus Bathyarchaeota archaeon
RLI28513.1 GCA 003662445 DUF2119 domain-containing protein Candidatus Bathyarchaeota archaeon
RJX15580.1 GCA 003604315 DUF2119 domain-containing protein partial Candidatus Bathyarchaeota archaeon
GCA 002011035MTLY01000003.1 3763257383262921 376 GCA 002011035 -1 ID 1 376 partial 00 Candidatus Bathyarchaeota archaeon JdFR-11
RLG91506.1 GCA 003661965 DUF2119 domain-containing protein Candidatus Bathyarchaeota archaeon
RLI05554.1 GCA 003662145 DUF2119 domain-containing protein Candidatus Bathyarchaeota archaeon
GCA 002011075MTMA01000002.1 2023988245513 20 GCA 002011075 1 ID 3 20 partial 00 Candidatus Geothermarchaeota archaeon JdFR-13
Methanosarcinales (ANME-2)
Methanosarcinales
Methanoliparia
Tree scale: 1

**Supplementary Figure 29.** ML phylogeny of methanogenesis marker m37. Black circles indicate strongly supported branches (ultrafast bootstrap >=95, aLRT SH-like >=80).

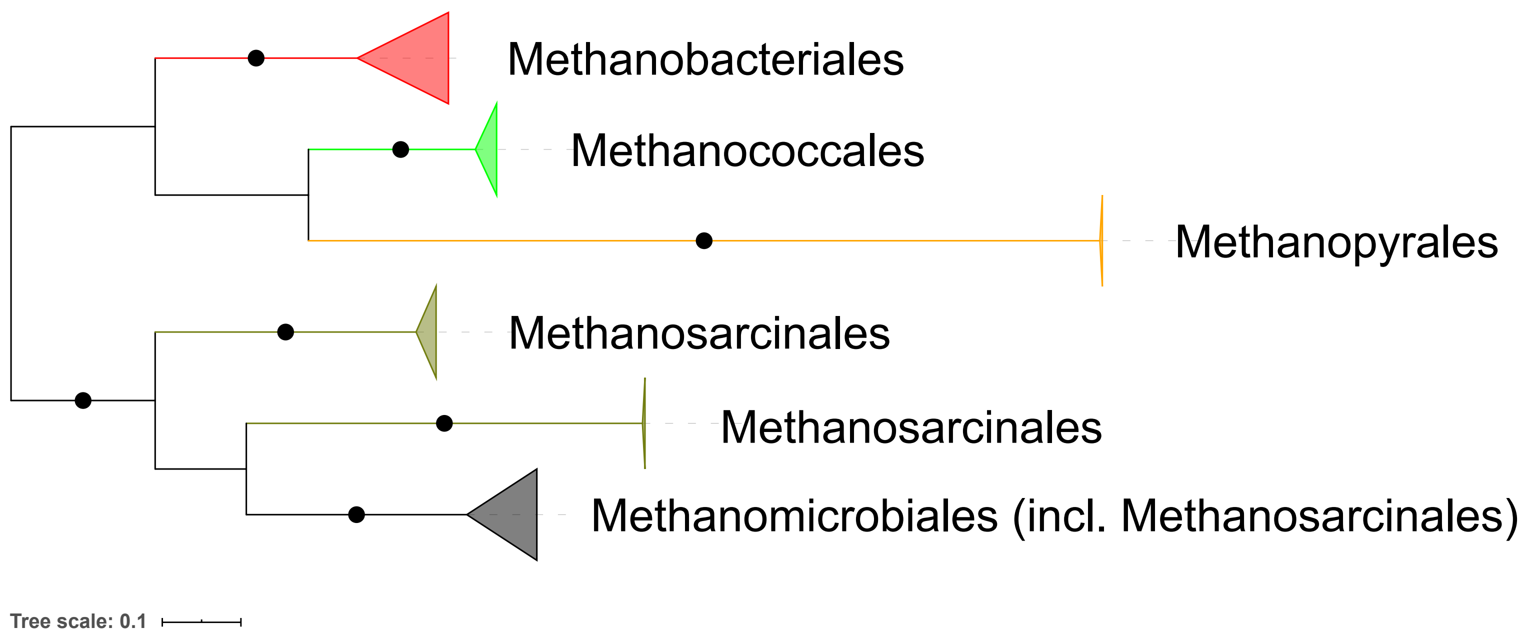
Methanobacteriales
Methanococcales
Methanopyrales
Methanosarcinales
Methanosarcinales
Methanomicrobiales (incl. Methanosarcinales)
Tree scale: 0.1

**Supplementary Figure 30.** ML phylogeny of methanogenesis marker m3. Black circles indicate strongly supported branches (ultrafast bootstrap >=95, aLRT SH-like >=80).

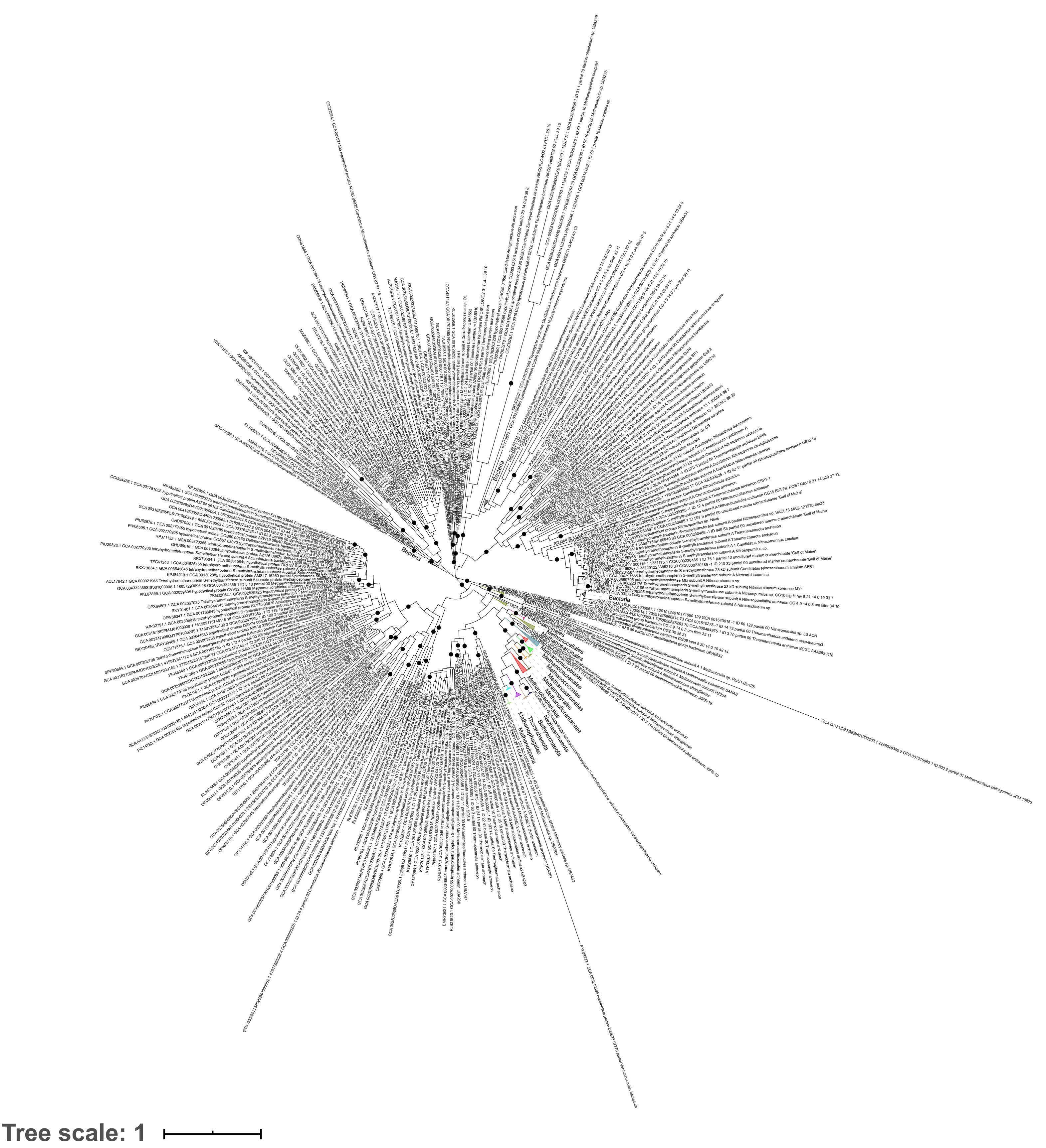
Bacteria
Bacteria
Bacteria
Methanosarcinales
Methanocellales
Methanomicrobiales
Methanobacteriales
Methanococcales
Methanopyrales
Bathyarchaeota
Tree scale: 1

**Supplementary Figure 31.** ML phylogeny of MtrA. Black circles indicate strongly supported branches (ultrafast bootstrap >=95, aLRT SH-like >=80).

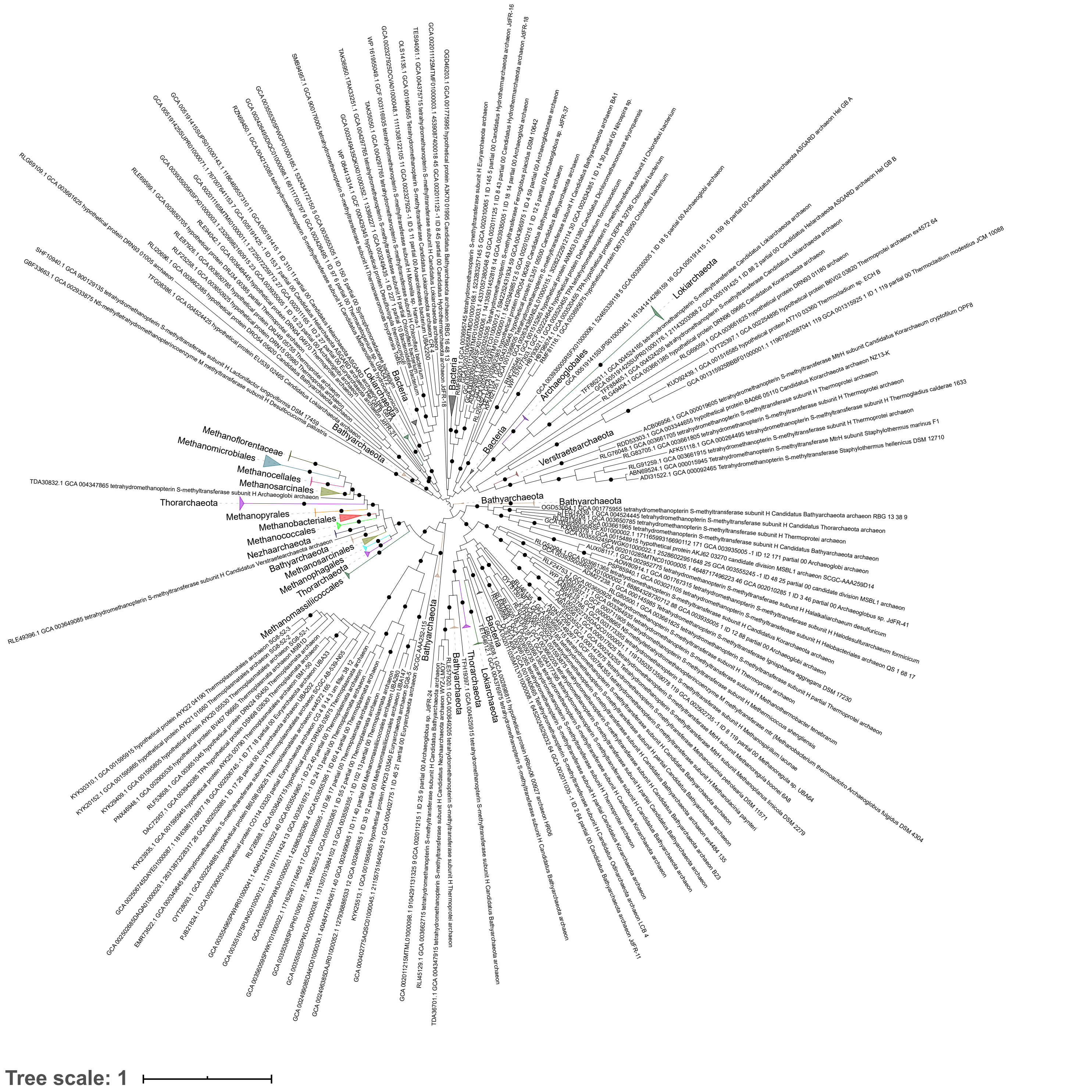

Lokiarchaeota
Achaeoglobales
Bacteria
Verstraetearchaeota
Bathyarchaeota
Thorarchaeota
Methanoflorentaceae
Methanomicrobiales
Methanocellales
Methanosarcinales
Methanopyrales
Methanobacteriales
Methanococcales
Nezhaarchaeota
Methanophagales
Methanomassiliicoccales
Tree scale: 1

**Supplementary Figure 32.** ML phylogeny of MtrH. Black circles indicate strongly supported branches (ultrafast bootstrap >=95, aLRT SH-like >=80).

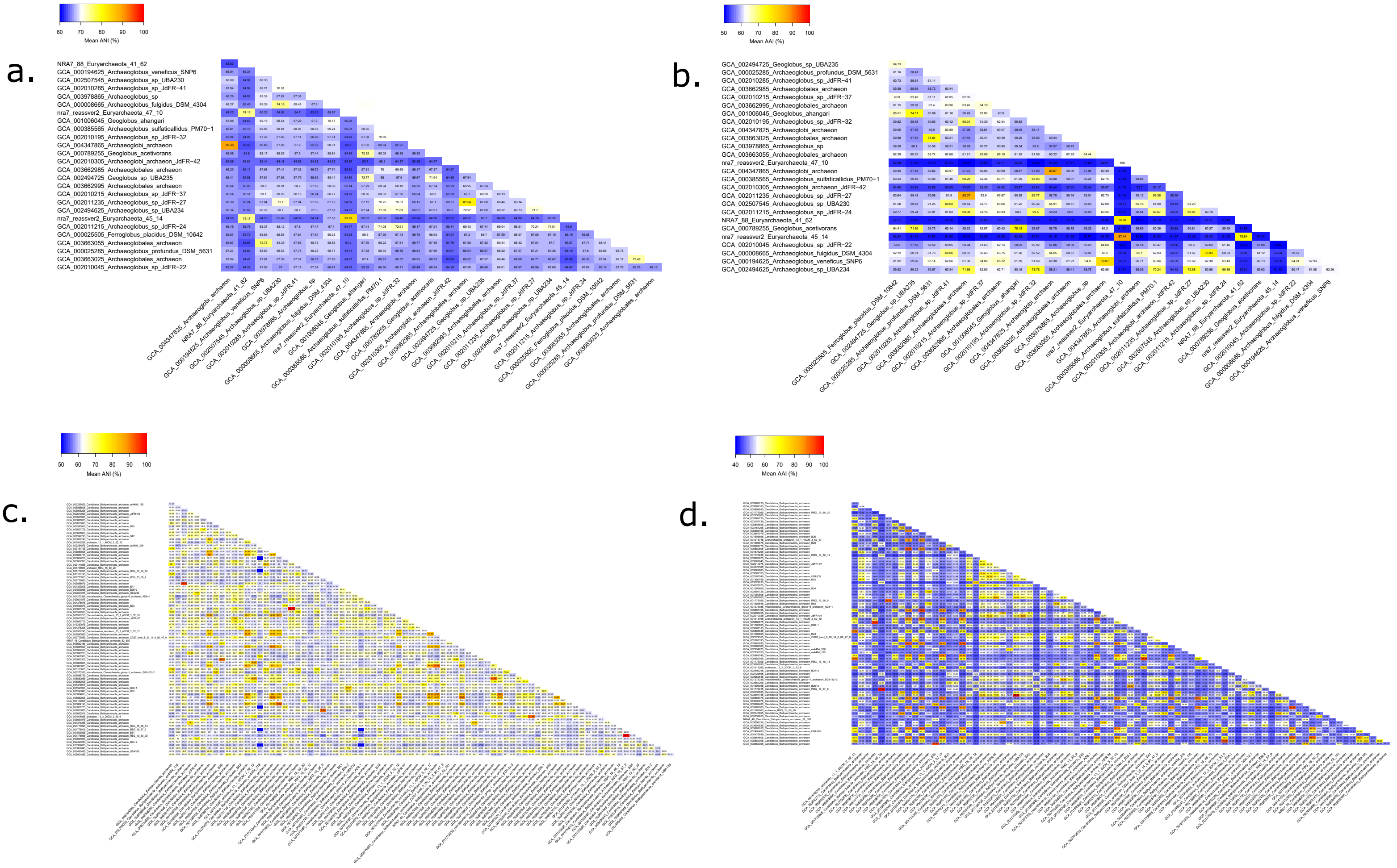

a.
60 70 80 90 100
Mean ANI (%)
NRA7_88_Euryarchaeota_41_62
GCA_000194625_Archaeoglobus_veneficus_SNP6
GCA_002507545_Archaeoglobus_sp_UBA230
GCA_002010285_Archaeoglobus_sp_JdFR−41
GCA_003978865_Archaeoglobus_sp
GCA_000008665_Archaeoglobus_fulgidus_DSM_4304
nra7_reassver2_Euryarchaeota_47_10
GCA_001006045_Geoglobus_ahangari
GCA_000385565_Archaeoglobus_sulfaticallidus_PM70−1
GCA_002010195_Archaeoglobus_sp_JdFR−32
GCA_004347865_Archaeoglobi_archaeon
GCA_000789255_Geoglobus_acetivorans
GCA_002010305_Archaeoglobi_archaeon_JdFR−42
GCA_003662985_Archaeoglobales_archaeon
GCA_002494725_Geoglobus_sp_UBA235
GCA_003662995_Archaeoglobales_archaeon
GCA_002010215_Archaeoglobus_sp_JdFR−37
GCA_002011235_Archaeoglobus_sp_JdFR−27
GCA_002494625_Archaeoglobus_sp_UBA234
nra7_reassver2_Euryarchaeota_45_14
GCA_002011215_Archaeoglobus_sp_JdFR−24
GCA_000025505_Ferroglobus_placidus_DSM_10642
GCA_003663055_Archaeoglobales_archaeon
GCA_000025285_Archaeoglobus_profundus_DSM_5631
GCA_003663025_Archaeoglobales_archaeon
GCA_002010045_Archaeoglobus_sp_JdFR−22
b.
50 60 70 80 90 100
Mean AAI (%)
GCA_002494725_Geoglobus_sp_UBA235
GCA_000025285_Archaeoglobus_profundus_DSM_5631
GCA_002010285_Archaeoglobus_sp_JdFR−41
GCA_003662985_Archaeoglobales_archaeon
GCA_002010215_Archaeoglobus_sp_JdFR−37
GCA_003662995_Archaeoglobales_archaeon
GCA_001006045_Geoglobus_ahangari
GCA_002010195_Archaeoglobus_sp_JdFR−32
GCA_004347825_Archaeoglobi_archaeon
GCA_003663025_Archaeoglobales_archaeon
GCA_003978865_Archaeoglobus_sp
GCA_003663055_Archaeoglobales_archaeon
nra7_reassver2_Euryarchaeota_47_10
GCA_004347865_Archaeoglobi_archaeon
GCA_000385565_Archaeoglobus_sulfaticallidus_PM70−1
GCA_002010305_Archaeoglobi_archaeon_JdFR−42
GCA_002011235_Archaeoglobus_sp_JdFR−27
GCA_002507545_Archaeoglobus_sp_UBA230
GCA_002011215_Archaeoglobus_sp_JdFR−24
NRA7_88_Euryarchaeota_41_62
GCA_000789255_Geoglobus_acetivorans
nra7_reassver2_Euryarchaeota_45_14
GCA_002010045_Archaeoglobus_sp_JdFR−22
GCA_000008665_Archaeoglobus_fulgidus_DSM_4304
GCA_000194625_Archaeoglobus_veneficus_SNP6
GCA_002494625_Archaeoglobus_sp_UBA234
c.
50 60 70 80 90 100
Mean ANI (%)
d.
40 50 60 70 80 90 100
Mean AAI (%)

**Supplementary Figure 33.** Heatmap graphs of (a) mean pairwise ANI (%), (b) mean pairwise AAI (%) of Archaeoglobi GTDB representative genomes with the Mnemosynellales from this study (*Ca.* M. biddleae substituting JdFR-21), (c) mean pairwise ANI (%), and (d) mean pairwise AAI (%) of Bathyarchaeia GTDB representative genomes with *Ca.* H. orcuttiae substituting JdFR-11.

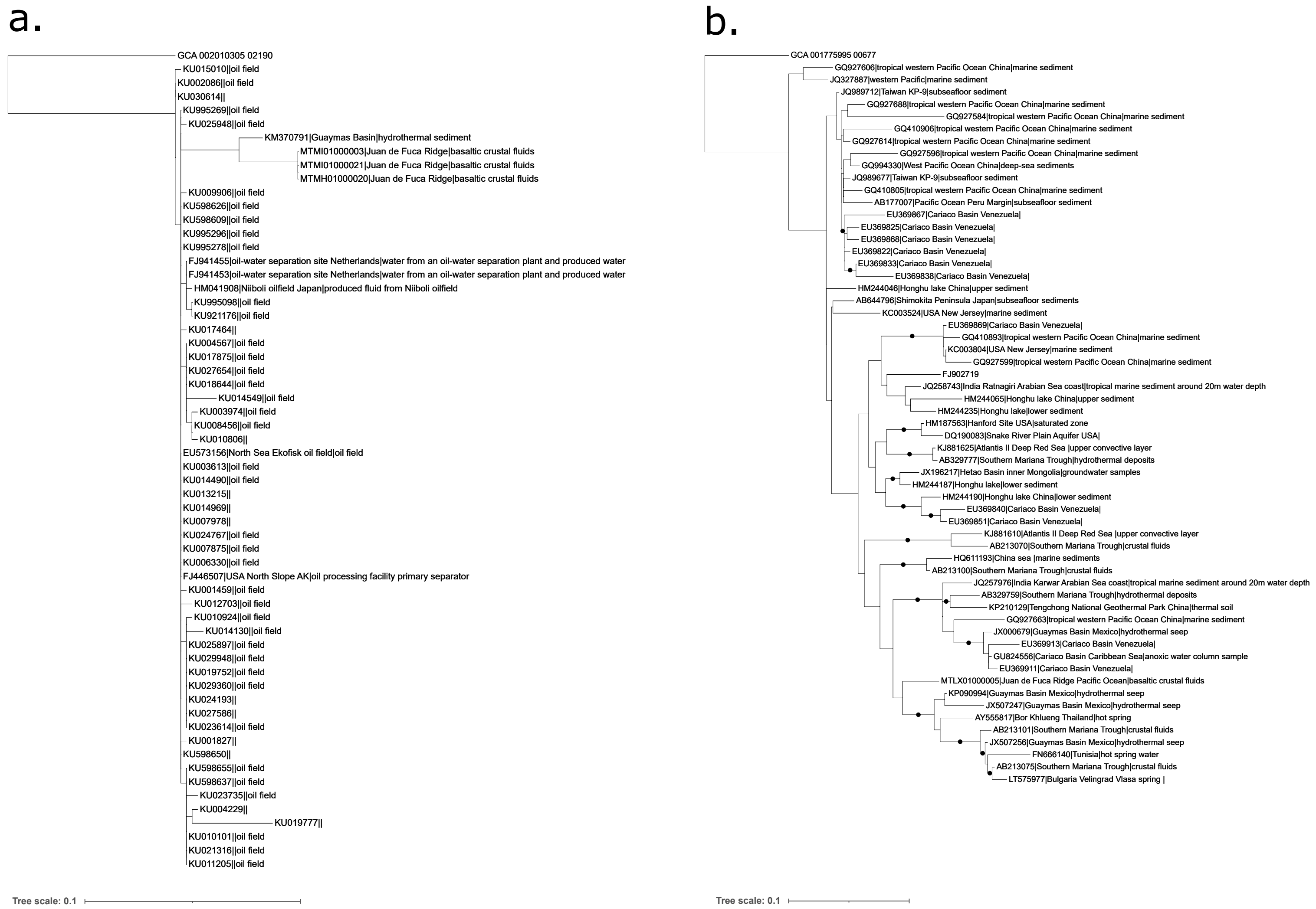
a.
GCA 002010305 02190
KU015010||oil field
KU002086||oil field
KU030614||
KU995269||oil field
KU025948||oil field
KM370791|Guaymas Basin|hydrothermal sediment
MTMI01000003|Juan de Fuca Ridge|basaltic crustal fluids
MTMI01000021|Juan de Fuca Ridge|basaltic crustal fluids
MTMH01000020|Juan de Fuca Ridge|basaltic crustal fluids
KU009906||oil field
KU598626||oil field
KU598609||oil field
KU995296||oil field
KU995278||oil field
FJ941455|oil-water separation site Netherlands|water from an oil-water separation plant and produced water
FJ941453|oil-water separation site Netherlands|water from an oil-water separation plant and produced water
HM041908|Niiboli oilfield Japan|produced fluid from Niiboli oilfield
KU995098||oil field
KU921176||oil field
KU017464||
KU004567||oil field
KU017875||oil field
KU027654||oil field
KU018644||oil field
KU014549||oil field
KU003974||oil field
KU008456||oil field
KU010806||
EU573156|North Sea Ekofisk oil field|oil field
KU003613||oil field
KU014490||oil field
KU013215||
KU014969||
KU007978||
KU024767||oil field
KU007875||oil field
KU006330||oil field
FJ446507|USA North Slope AK|oil processing facility primary separator
KU001459||oil field
KU012703||oil field
KU010924||oil field
KU014130||oil field
KU025897||oil field
KU029948||oil field
KU019752||oil field
KU029360||oil field
KU024193||
KU027586||
KU023614||oil field
KU001827||
KU598650||
KU598655||oil field
KU598637||oil field
KU023735||oil field
KU004229||
KU019777||
KU010101||oil field
KU021316||oil field
KU011205||oil field
Tree scale: 0.1
b.
GCA 001775995 00677
GQ927606|tropical western Pacific Ocean China|marine sediment
JQ327887|western Pacific|marine sediment
JQ989712|Taiwan KP-9|subseafloor sediment
GQ927688|tropical western Pacific Ocean China|marine sediment
GQ927584|tropical western Pacific Ocean China|marine sediment
GQ410906|tropical western Pacific Ocean China|marine sediment
GQ927614|tropical western Pacific Ocean China|marine sediment
GQ927596|tropical western Pacific Ocean China|marine sediment
GQ994330|West Pacific Ocean China|deep-sea sediments
JQ989677|Taiwan KP-9|subseafloor sediment
GQ410805|tropical western Pacific Ocean China|marine sediment
AB177007|Pacific Ocean Peru Margin|subseafloor sediment
EU369867|Cariaco Basin Venezuela|
EU369825|Cariaco Basin Venezuela|
EU369868|Cariaco Basin Venezuela|
EU369822|Cariaco Basin Venezuela|
EU369833|Cariaco Basin Venezuela|
EU369838|Cariaco Basin Venezuela|
HM244046|Honghu lake China|upper sediment
AB644796|Shimokita Peninsula Japan|subseafloor sediments
KC003524|USA New Jersey|marine sediment
EU369869|Cariaco Basin Venezuela|
GQ410893|tropical western Pacific Ocean China|marine sediment
KC003804|USA New Jersey|marine sediment
GQ927599|tropical western Pacific Ocean China|marine sediment
FJ902719
JQ258743|India Ratnagiri Arabian Sea coast|tropical marine sediment around 20m water depth
HM244065|Honghu lake China|upper sediment
HM244235|Honghu lake|lower sediment
HM187563|Hanford Site USA|saturated zone
DQ190083|Snake River Plain Aquifer USA|
KJ881625|Atlantis II Deep Red Sea |upper convective layer
AB329777|Southern Mariana Trough|hydrothermal deposits
JX196217|Hetao Basin inner Mongolia|groundwater samples
HM244187|Honghu lake|lower sediment
HM244190|Honghu lake China|lower sediment
EU369840|Cariaco Basin Venezuela|
EU369851|Cariaco Basin Venezuela|
KJ881610|Atlantis II Deep Red Sea |upper convective layer
AB213070|Southern Mariana Trough|crustal fluids
HQ611193|China sea |marine sediments
AB213100|Southern Mariana Trough|crustal fluids
JQ257976|India Karwar Arabian Sea coast|tropical marine sediment around 20m water depth
AB329759|Southern Mariana Trough|hydrothermal deposits
KP210129|Tengchong National Geothermal Park China|thermal soil
GQ927663|tropical western Pacific Ocean China|marine sediment
JX000679|Guaymas Basin Mexico|hydrothermal seep
EU369913|Cariaco Basin Venezuela|
GU824556|Cariaco Basin Caribbean Sea|anoxic water column sample
EU369911|Cariaco Basin Venezuela|
MTLX01000005|Juan de Fuca Ridge Pacific Ocean|basaltic crustal fluids
KP090994|Guaymas Basin Mexico|hydrothermal seep
JX507247|Guaymas Basin Mexico|hydrothermal seep
AY555817|Bor Khlueng Thailand|hot spring
AB213101|Southern Mariana Trough|crustal fluids
JX507256|Guaymas Basin Mexico|hydrothermal seep
FN666140|Tunisia|hot spring water
AB213075|Southern Mariana Trough|crustal fluids
LT575977|Bulgaria Velingrad Vlasa spring |
Tree scale: 0.1

**Supplementary Figure 34.** ML phylogenies of the 16S genes in (a) Mnemosynellales and (b) Hecatellales. Each leaf contains the sequence accession, pipe separated with its sampling locality, and environment, depending on their availability in each sequence's metadata in SILVA. Black circles indicate strongly supported branches (ultrafast bootstrap >=95, aLRT SH-like >=80).

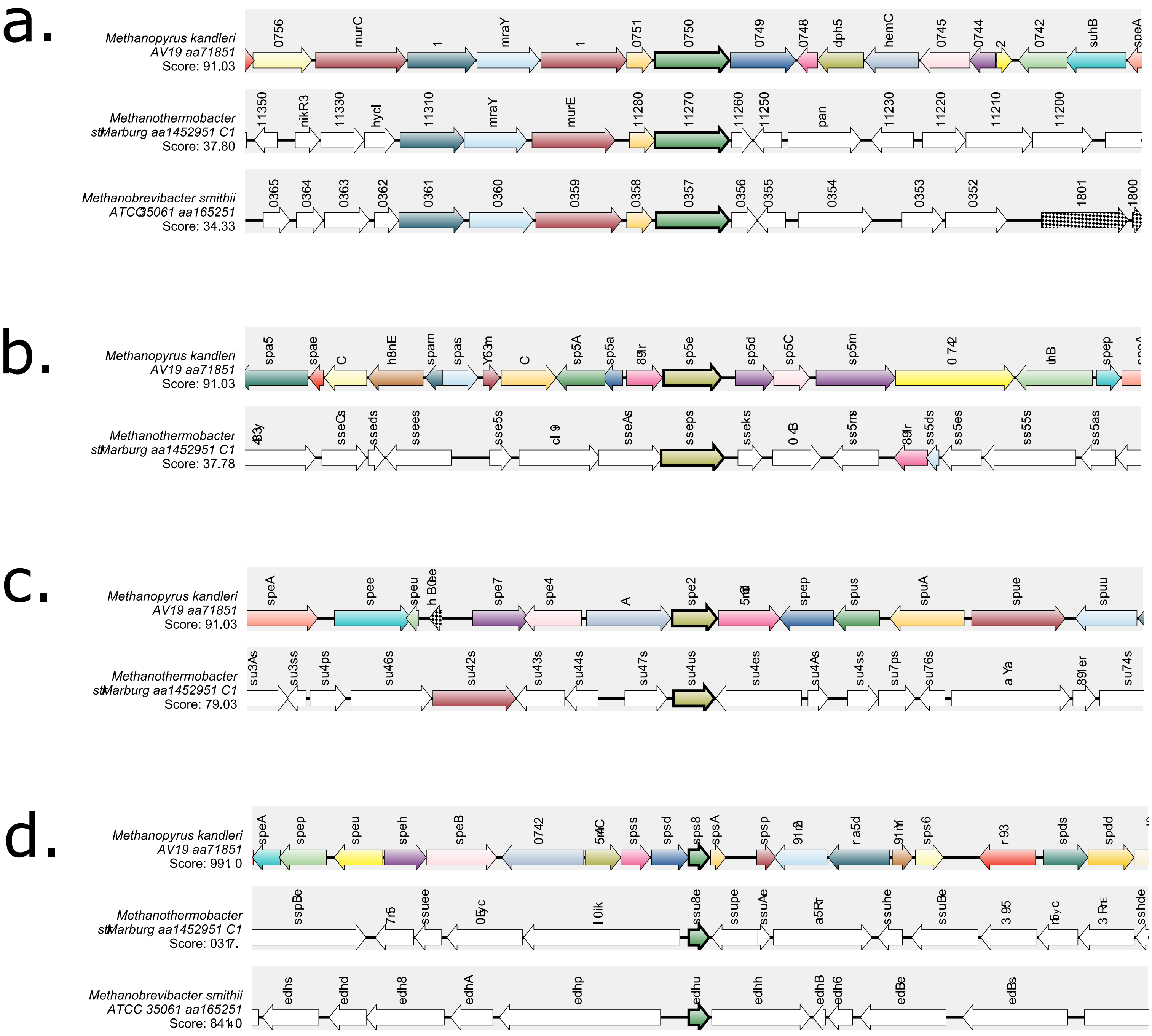

Methanopyrus kandleri
AV19 aa71851
Score: 91.03
0756
murC
mraY
0751
0750
0749
0748
dph5
hemC
0745
0744
0742
suhB
speA
Methanothermobacter
stMarburg aa1452951 C1
Score: 37.80
11350
nikR3
11330
hycI
11310
mraY
murE
11280
11270
11260
11250
pan
11230
11220
11210
11200
Methanobrevibacter smithii
ATCC 35061 aa165251
Score: 34.33
0365
0364
0363
0362
0361
0360
0359
0358
0357
0356
0355
0354
0353
0352
1801
1800
Methanopyrus kandleri
AV19 aa71851
Score: 91.03
Methanothermobacter
stMarburg aa1452951 C1
Score: 37.78
Methanopyrus kandleri
AV19 aa71851
Score: 91.03
Methanothermobacter
stMarburg aa1452951 C1
Score: 79.03
Methanopyrus kandleri
AV19 aa71851
Methanothermobacter
stMarburg aa1452951 C1
Methanobrevibacter smithii
ATCC 35061 aa165251

**Supplementary Figure 35.** Graphical representation of synteny for genes (a) MK0750, (b) MK0854, (c) MK0927, and (d) MK1513 in the genomes of *Methanopyrus kandleri*, *Methanothermobacter marburgensis str. Marburg*, and *Methanobrevibacter smithii*.

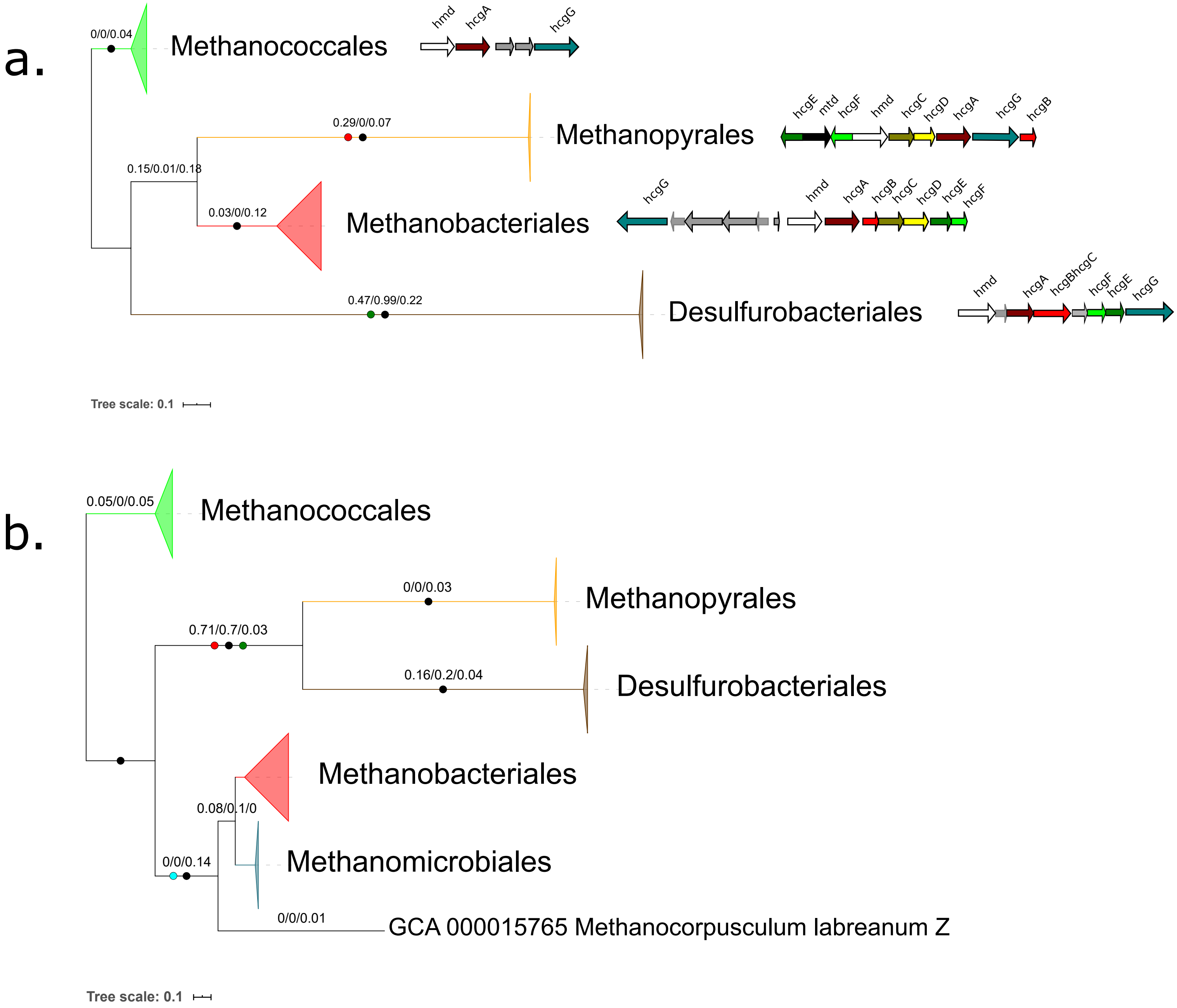

a.
0/0/0.04
Methanococcales
hmd
hcgA
hcgG
0.29/0/0.07
Methanopyrales
hcgE
mtd
hcgF
hmd
hcgC
hcgD
hcgA
hcgG
hcgB
0.15/0.01/0.18
0.03/0/0.12
Methanobacteriales
hcgG
hmd
hcgA
hcgB
hcgC
hcgD
hcgE
hcgF
0.47/0.99/0.22
Desulfurobacteriales
hmd
hcgA
hcgBhcgC
hcgF
hcgE
hcgG
Tree scale: 0.1
b.
0.05/0/0.05
Methanococcales
0/0/0.03
Methanopyrales
0.71/0.7/0.03
0.16/0.2/0.04
Desulfurobacteriales
Methanobacteriales
0.08/0.1/0
Methanomicrobiales
0/0/0.14
0/0/0.01
GCA 000015765 Methanocorpusculum labreanum Z
Tree scale: 0.1

**Supplementary Figure 36.** ML phylogenies of (a) HcgAEFG (938 aa positions) along with the genomic organization of the clusters in a representative genome for each major clade, (b) HcgBC (366 aa positions). Black circles indicate strongly supported branches (ultrafast bootstrap >=95, aLRT SH-like >=80), red circles correspond to the MAD root, green to MinVar, light blue to NONREV. Branch values correspond to rootstrap supports for MAD, MinVar, and NONREV respectively. For HcgAEFG, the NONREV root is within a collapsed clade. Genomes used to compare clusters were the same as Figure 3, with additionally Desulfurobacterium thermolithotrophum DSM 11699 (GCA_000191045; Desulfurobacteriales).

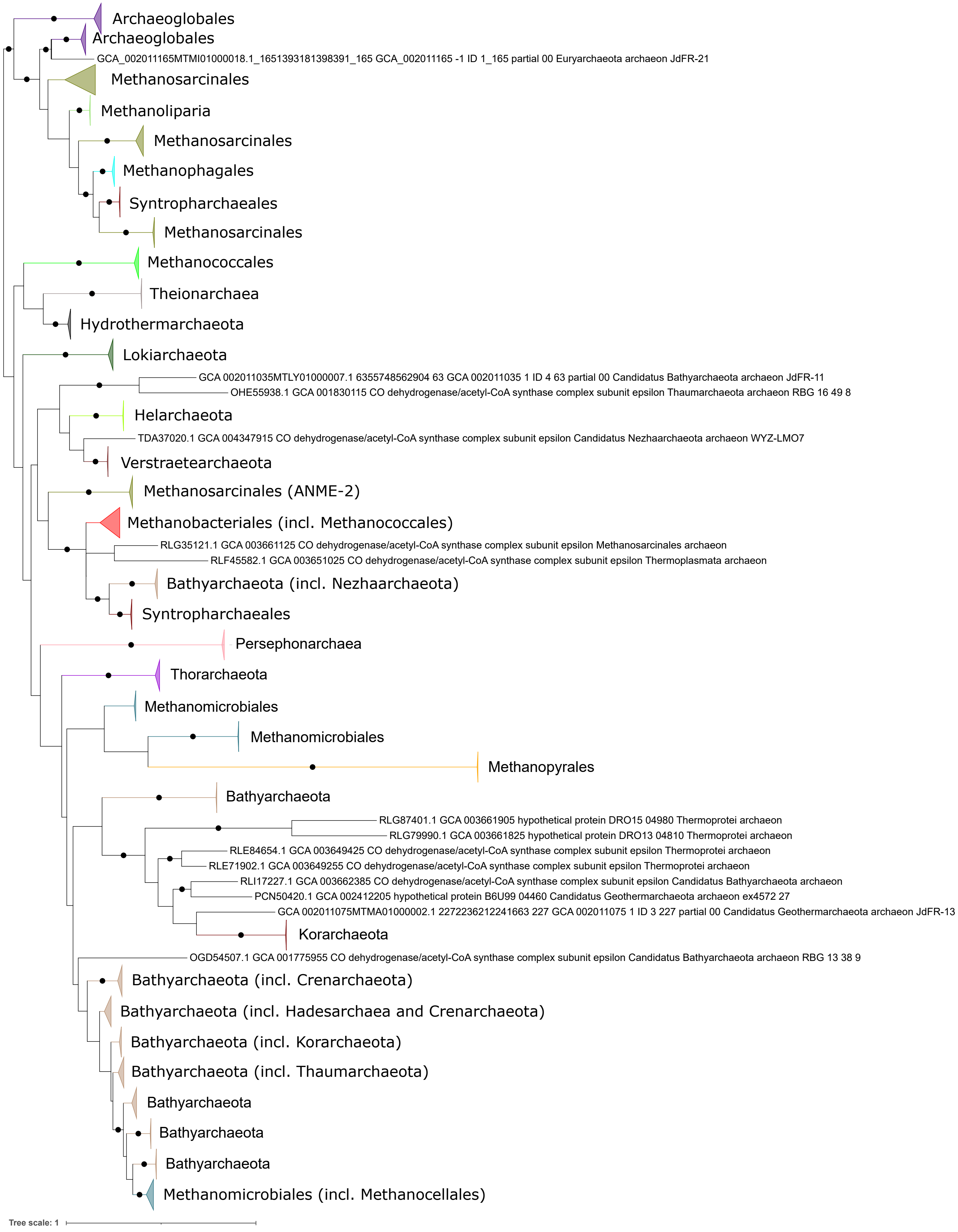

Archaeoglobales
Archaeoglobales
GCA_002011165MTMI01000018.1_1651393181398391_165 GCA_002011165 -1 ID 1_165 partial 00 Euryarchaeota archaeon JdFR-21
Methanosarcinales
Methanoliparia
Methanosarcinales
Methanophagales
Syntropharchaeales
Methanosarcinales
Methanococcales
Theionarchaea
Hydrothermarchaeota
Lokiarchaeota
GCA 002011035MTLY01000007.1 6355748562904 63 GCA 002011035 1 ID 4 63 partial 00 Candidatus Bathyarchaeota archaeon JdFR-11
OHE55938.1 GCA 001830115 CO dehydrogenase/acetyl-CoA synthase complex subunit epsilon Thaumarchaeota archaeon RBG 16 49 8
Helarchaeota
TDA37020.1 GCA 004347915 CO dehydrogenase/acetyl-CoA synthase complex subunit epsilon Candidatus Nezhaarchaeota archaeon WYZ-LMO7
Verstraetearchaeota
Methanosarcinales (ANME-2)
Methanobacteriales (incl. Methanococcales)
RLG35121.1 GCA 003661125 CO dehydrogenase/acetyl-CoA synthase complex subunit epsilon Methanosarcinales archaeon
RLF45582.1 GCA 003651025 CO dehydrogenase/acetyl-CoA synthase complex subunit epsilon Thermoplasmata archaeon
Bathyarchaeota (incl. Nezhaarchaeota)
Syntropharchaeales
Persephonarchaea
Thorarchaeota
Methanomicrobiales
Methanomicrobiales
Methanopyrales
Bathyarchaeota
RLG87401.1 GCA 003661905 hypothetical protein DRO15 04980 Thermoprotei archaeon
RLG79990.1 GCA 003661825 hypothetical protein DRO13 04810 Thermoprotei archaeon
RLE84654.1 GCA 003649425 CO dehydrogenase/acetyl-CoA synthase complex subunit epsilon Thermoprotei archaeon
RLE71902.1 GCA 003649255 CO dehydrogenase/acetyl-CoA synthase complex subunit epsilon Thermoprotei archaeon
RLI17227.1 GCA 003662385 CO dehydrogenase/acetyl-CoA synthase complex subunit epsilon Candidatus Bathyarchaeota archaeon
PCN50420.1 GCA 002412205 hypothetical protein B6U99 04460 Candidatus Geothermarchaeota archaeon ex4572 27
GCA 002011075MTMA01000002.1 2272236212241663 227 GCA 002011075 1 ID 3 227 partial 00 Candidatus Geothermarchaeota archaeon JdFR-13
Korarchaeota
OGD54507.1 GCA 001775955 CO dehydrogenase/acetyl-CoA synthase complex subunit epsilon Candidatus Bathyarchaeota archaeon RBG 13 38 9
Bathyarchaeota (incl. Crenarchaeota)
Bathyarchaeota (incl. Hadesarchaea and Crenarchaeota)
Bathyarchaeota (incl. Korarchaeota)
Bathyarchaeota (incl. Thaumarchaeota)
Bathyarchaeota
Bathyarchaeota
Bathyarchaeota
Methanomicrobiales (incl. Methanocellales)
Tree scale: 1

**Supplementary Figure 37.** ML phylogeny of CdhB. Black circles indicate strongly supported branches (ultrafast bootstrap >=95, aLRT SH-like >=80).

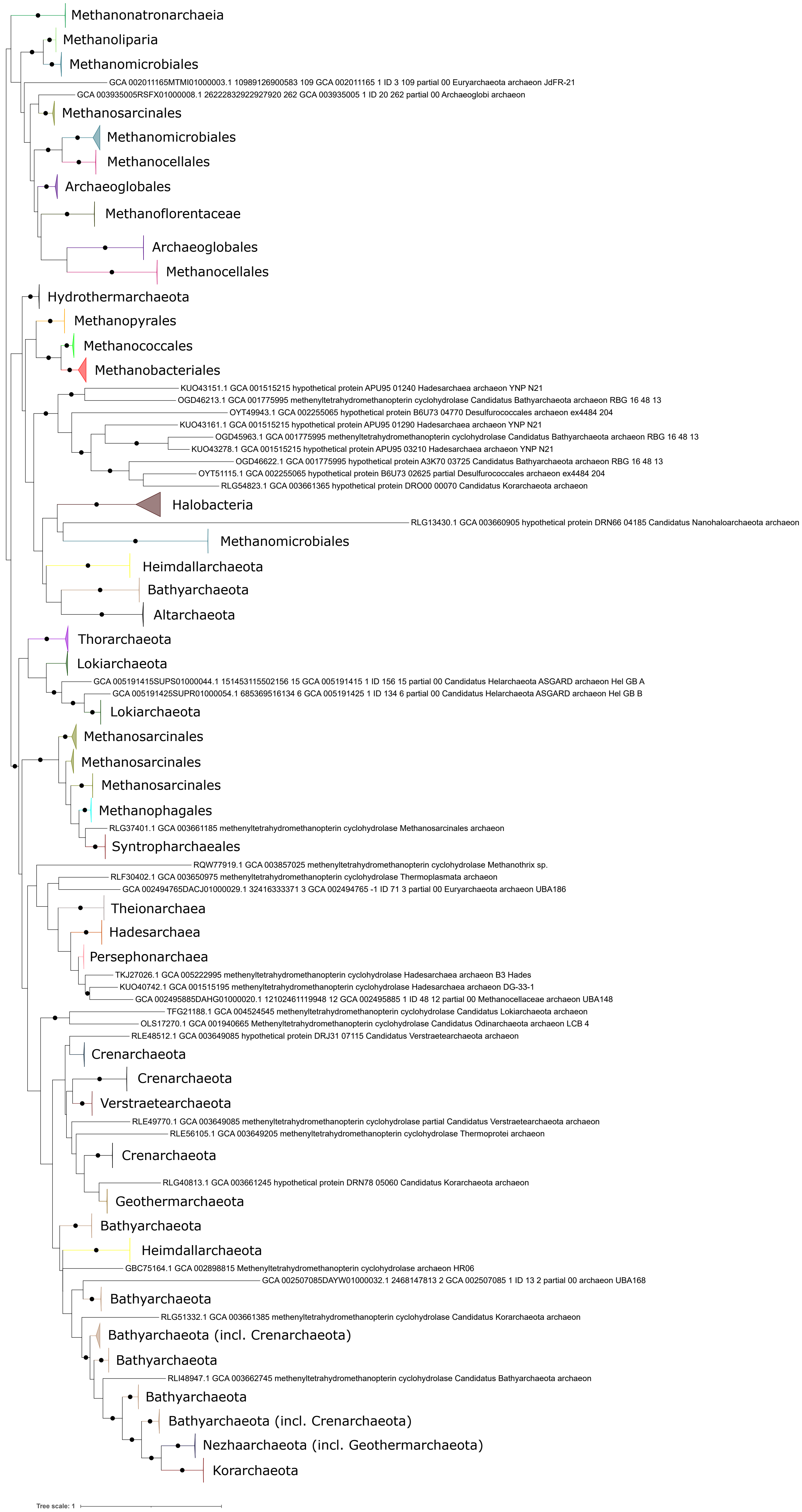
Methanonatronarchaeia
Methanoliparia
Methanomicrobiales
GCA 002011165MTMI01000003.1 10989126900583 109 GCA 002011165 1 ID 3 109 partial 00 Euryarchaeota archaeon JdFR-21
GCA 003935005RSFX01000008.1 26222832922927920 262 GCA 003935005 1 ID 20 262 partial 00 Archaeoglobi archaeon
Methanosarcinales
Methanomicrobiales
Methanocellales
Archaeoglobales
Methanoflorentaceae
Archaeoglobales
Methanocellales
Hydrothermarchaeota
Methanopyrales
Methanococcales
Methanobacteriales
KUO43151.1 GCA 001515215 hypothetical protein APU95 01240 Hadesarchaea archaeon YNP N21
OGD46213.1 GCA 001775995 methenyltetrahydromethanopterin cyclohydrolase Candidatus Bathyarchaeota archaeon RBG 16 48 13
OYT49943.1 GCA 002255065 hypothetical protein B6U73 04770 Desulfurococcales archaeon ex4484 204
KUO43161.1 GCA 001515215 hypothetical protein APU95 01290 Hadesarchaea archaeon YNP N21
OGD45963.1 GCA 001775995 methenyltetrahydromethanopterin cyclohydrolase Candidatus Bathyarchaeota archaeon RBG 16 48 13
KUO43278.1 GCA 001515215 hypothetical protein APU95 03210 Hadesarchaea archaeon YNP N21
OGD46622.1 GCA 001775995 hypothetical protein A3K70 03725 Candidatus Bathyarchaeota archaeon RBG 16 48 13
OYT51115.1 GCA 002255065 hypothetical protein B6U73 02625 partial Desulfurococcales archaeon ex4484 204
RLG54823.1 GCA 003661365 hypothetical protein DRO00 00070 Candidatus Korarchaeota archaeon
Halobacteria
RLG13430.1 GCA 003660905 hypothetical protein DRN66 04185 Candidatus Nanohaloarchaeota archaeon
Methanomicrobiales
Heimdallarchaeota
Bathyarchaeota
Altarchaeota
Thorarchaeota
Lokiarchaeota
GCA 005191415SUPS01000044.1 151453115502156 15 GCA 005191415 1 ID 156 15 partial 00 Candidatus Helarchaeota ASGARD archaeon Hel GB A
GCA 005191425SUPR01000054.1 685369516134 6 GCA 005191425 1 ID 134 6 partial 00 Candidatus Helarchaeota ASGARD archaeon Hel GB B
Lokiarchaeota
Methanosarcinales
Methanosarcinales
Methanosarcinales
Methanophagales
RLG37401.1 GCA 003661185 methenyltetrahydromethanopterin cyclohydrolase Methanosarcinales archaeon
Syntropharchaeales
RQW77919.1 GCA 003857025 methenyltetrahydromethanopterin cyclohydrolase Methanothrix sp.
RLF30402.1 GCA 003650975 methenyltetrahydromethanopterin cyclohydrolase Thermoplasmata archaeon
GCA 002494765DACJ01000029.1 32416333371 3 GCA 002494765 -1 ID 71 3 partial 00 Euryarchaeota archaeon UBA186
Theionarchaea
Hadesarchaea
Persephonarchaea
TKJ27026.1 GCA 005222995 methenyltetrahydromethanopterin cyclohydrolase Hadesarchaea archaeon B3 Hades
KUO40742.1 GCA 001515195 methenyltetrahydromethanopterin cyclohydrolase Hadesarchaea archaeon DG-33-1
GCA 002495885DAHG01000020.1 12102461119948 12 GCA 002495885 1 ID 48 12 partial 00 Methanocellaceae archaeon UBA148
TFG21188.1 GCA 004524545 methenyltetrahydromethanopterin cyclohydrolase Candidatus Lokiarchaeota archaeon
OLS17270.1 GCA 001940665 Methenyltetrahydromethanopterin cyclohydrolase Candidatus Odinarchaeota archaeon LCB 4
RLE48512.1 GCA 003649085 hypothetical protein DRJ31 07115 Candidatus Verstraetearchaeota archaeon
Crenarchaeota
Crenarchaeota
Verstraetearchaeota
RLE49770.1 GCA 003649085 methenyltetrahydromethanopterin cyclohydrolase partial Candidatus Verstraetearchaeota archaeon
RLE56105.1 GCA 003649205 methenyltetrahydromethanopterin cyclohydrolase Thermoprotei archaeon
Crenarchaeota
RLG40813.1 GCA 003661245 hypothetical protein DRN78 05060 Candidatus Korarchaeota archaeon
Geothermarchaeota
Bathyarchaeota
Heimdallarchaeota
GBC75164.1 GCA 002898815 Methenyltetrahydromethanopterin cyclohydrolase archaeon HR06
GCA 002507085DAYW01000032.1 2468147813 2 GCA 002507085 1 ID 13 2 partial 00 archaeon UBA168
Bathyarchaeota
RLG51332.1 GCA 003661385 methenyltetrahydromethanopterin cyclohydrolase Candidatus Korarchaeota archaeon
Bathyarchaeota (incl. Crenarchaeota)
Bathyarchaeota
RLI48947.1 GCA 003662745 methenyltetrahydromethanopterin cyclohydrolase Candidatus Bathyarchaeota archaeon
Bathyarchaeota
Bathyarchaeota (incl. Crenarchaeota)
Nezhaarchaeota (incl. Geothermarchaeota)
Korarchaeota
Tree scale: 1

**Supplementary Figure 38.** ML phylogeny of Mch. Black circles indicate strongly supported branches (ultrafast bootstrap >=95, aLRT SH-like >=80).

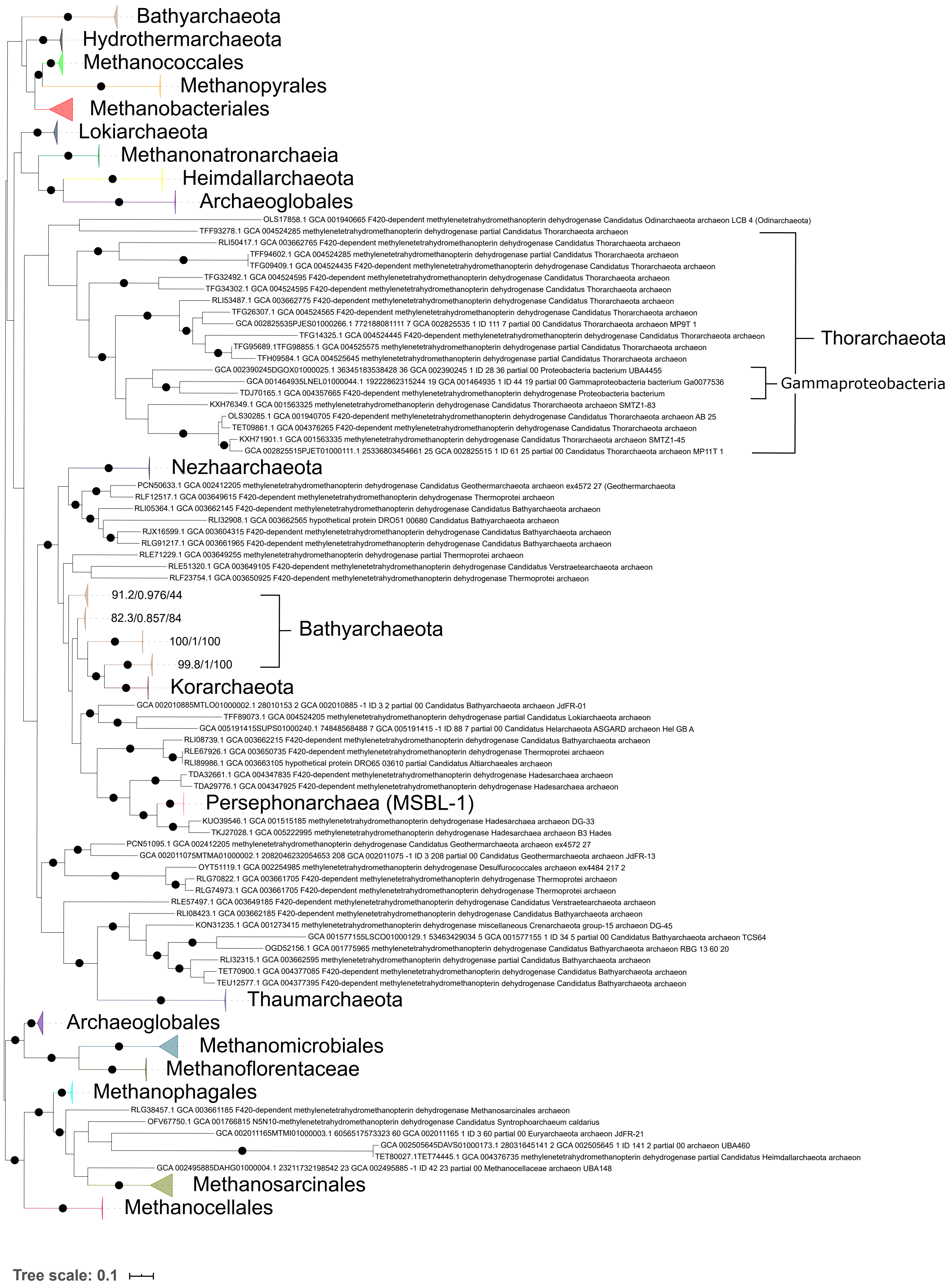
Bathyarchaeota
Hydrothermarchaeota
Methanococcales
Methanopyrales
Methanobacteriales
Lokiarchaeota
Methanonatronarchaeia
Heimdallarchaeota
Archaeoglobales
OLS17858.1 GCA 001940665 F420-dependent methylenetetrahydromethanopterin dehydrogenase Candidatus Odinarchaeota archaeon LCB 4 (Odinarchaeota)
TFF93278.1 GCA 004524285 methylenetetrahydromethanopterin dehydrogenase partial Candidatus Thorarchaeota archaeon
RLI50417.1 GCA 003662765 F420-dependent methylenetetrahydromethanopterin dehydrogenase Candidatus Thorarchaeota archaeon
TFF94602.1 GCA 004524285 methylenetetrahydromethanopterin dehydrogenase partial Candidatus Thorarchaeota archaeon
TFG09409.1 GCA 004524435 F420-dependent methylenetetrahydromethanopterin dehydrogenase Candidatus Thorarchaeota archaeon
TFG32492.1 GCA 004524595 F420-dependent methylenetetrahydromethanopterin dehydrogenase Candidatus Thorarchaeota archaeon
TFG34302.1 GCA 004524595 F420-dependent methylenetetrahydromethanopterin dehydrogenase Candidatus Thorarchaeota archaeon
RLI53487.1 GCA 003662775 F420-dependent methylenetetrahydromethanopterin dehydrogenase Candidatus Thorarchaeota archaeon
TFG26307.1 GCA 004524565 F420-dependent methylenetetrahydromethanopterin dehydrogenase Candidatus Thorarchaeota archaeon
GCA 002825535PJES01000266.1 772188081111 7 GCA 002825535 1 ID 111 7 partial 00 Candidatus Thorarchaeota archaeon MP9T 1
TFG14325.1 GCA 004524445 F420-dependent methylenetetrahydromethanopterin dehydrogenase Candidatus Thorarchaeota archaeon
TFG95689.1TFG98855.1 GCA 004525575 methylenetetrahydromethanopterin dehydrogenase partial Candidatus Thorarchaeota archaeon
TFH09584.1 GCA 004525645 methylenetetrahydromethanopterin dehydrogenase partial Candidatus Thorarchaeota archaeon
GCA 002390245DGOX01000025.1 36345183538428 36 GCA 002390245 1 ID 28 36 partial 00 Proteobacteria bacterium UBA4455
GCA 001464935LNEL01000044.1 19222862315244 19 GCA 001464935 1 ID 44 19 partial 00 Gammaproteobacteria bacterium Ga0077536
TDJ70165.1 GCA 004357665 F420-dependent methylenetetrahydromethanopterin dehydrogenase Proteobacteria bacterium
KXH76349.1 GCA 001563325 methylenetetrahydromethanopterin dehydrogenase Candidatus Thorarchaeota archaeon SMTZ1-83
OLS30285.1 GCA 001940705 F420-dependent methylenetetrahydromethanopterin dehydrogenase Candidatus Thorarchaeota archaeon AB 25
TET09861.1 GCA 004376265 F420-dependent methylenetetrahydromethanopterin dehydrogenase Candidatus Thorarchaeota archaeon
KXH71901.1 GCA 001563335 methylenetetrahydromethanopterin dehydrogenase Candidatus Thorarchaeota archaeon SMTZ1-45
GCA 002825515PJET01000111.1 25336803454661 25 GCA 002825515 1 ID 61 25 partial 00 Candidatus Thorarchaeota archaeon MP11T 1
Thorarchaeota
Gammaproteobacteria
Nezhaarchaeota
PCN50633.1 GCA 002412205 methylenetetrahydromethanopterin dehydrogenase Candidatus Geothermarchaeota archaeon ex4572 27 (Geothermarchaeota
RLF12517.1 GCA 003649615 F420-dependent methylenetetrahydromethanopterin dehydrogenase Thermoprotei archaeon
RLI05364.1 GCA 003662145 F420-dependent methylenetetrahydromethanopterin dehydrogenase Candidatus Bathyarchaeota archaeon
RLI32908.1 GCA 003662565 hypothetical protein DRO51 00680 Candidatus Bathyarchaeota archaeon
RJX16599.1 GCA 003604315 F420-dependent methylenetetrahydromethanopterin dehydrogenase Candidatus Bathyarchaeota archaeon
RLG91217.1 GCA 003661965 F420-dependent methylenetetrahydromethanopterin dehydrogenase Candidatus Bathyarchaeota archaeon
RLE71229.1 GCA 003649255 methylenetetrahydromethanopterin dehydrogenase partial Thermoprotei archaeon
RLE51320.1 GCA 003649105 F420-dependent methylenetetrahydromethanopterin dehydrogenase Candidatus Verstraetearchaeota archaeon
RLF23754.1 GCA 003650925 F420-dependent methylenetetrahydromethanopterin dehydrogenase Thermoprotei archaeon
91.2/0.976/44
82.3/0.857/84
100/1/100
99.8/1/100
Bathyarchaeota
Korarchaeota
GCA 002010885MTLO01000002.1 28010153 2 GCA 002010885 -1 ID 3 2 partial 00 Candidatus Bathyarchaeota archaeon JdFR-01
TFF89073.1 GCA 004524205 methylenetetrahydromethanopterin dehydrogenase partial Candidatus Lokiarchaeota archaeon
GCA 005191415SUPS01000240.1 74848568488 7 GCA 005191415 -1 ID 88 7 partial 00 Candidatus Helarchaeota ASGARD archaeon Hel GB A
RLI08739.1 GCA 003662215 F420-dependent methylenetetrahydromethanopterin dehydrogenase Candidatus Bathyarchaeota archaeon
RLE67926.1 GCA 003650735 F420-dependent methylenetetrahydromethanopterin dehydrogenase Thermoprotei archaeon
RLI89986.1 GCA 003663105 hypothetical protein DRO65 03610 partial Candidatus Altiarchaeales archaeon
TDA32661.1 GCA 004347835 F420-dependent methylenetetrahydromethanopterin dehydrogenase Hadesarchaea archaeon
TDA29776.1 GCA 004347925 F420-dependent methylenetetrahydromethanopterin dehydrogenase Hadesarchaea archaeon
Persephonarchaea (MSBL-1)
KUO39546.1 GCA 001515185 methylenetetrahydromethanopterin dehydrogenase Hadesarchaea archaeon DG-33
TKJ27028.1 GCA 005222995 methylenetetrahydromethanopterin dehydrogenase Hadesarchaea archaeon B3 Hades
PCN51095.1 GCA 002412205 methylenetetrahydromethanopterin dehydrogenase Candidatus Geothermarchaeota archaeon ex4572 27
GCA 002011075MTMA01000002.1 2082046232054653 208 GCA 002011075 -1 ID 3 208 partial 00 Candidatus Geothermarchaeota archaeon JdFR-13
OYT51119.1 GCA 002254985 methylenetetrahydromethanopterin dehydrogenase Desulfurococcales archaeon ex4484 217 2
RLG70822.1 GCA 003661705 F420-dependent methylenetetrahydromethanopterin dehydrogenase Thermoprotei archaeon
RLG74973.1 GCA 003661705 F420-dependent methylenetetrahydromethanopterin dehydrogenase Thermoprotei archaeon
RLE57497.1 GCA 003649185 F420-dependent methylenetetrahydromethanopterin dehydrogenase Candidatus Verstraetearchaeota archaeon
RLI08423.1 GCA 003662185 F420-dependent methylenetetrahydromethanopterin dehydrogenase Candidatus Bathyarchaeota archaeon
KON31235.1 GCA 001273415 methylenetetrahydromethanopterin dehydrogenase miscellaneous Crenarchaeota group-15 archaeon DG-45
GCA 001577155LSCO01000129.1 53463429034 5 GCA 001577155 1 ID 34 5 partial 00 Candidatus Bathyarchaeota archaeon TCS64
OGD52156.1 GCA 001775965 methylenetetrahydromethanopterin dehydrogenase Candidatus Bathyarchaeota archaeon RBG 13 60 20
RLI32315.1 GCA 003662595 methylenetetrahydromethanopterin dehydrogenase partial Candidatus Bathyarchaeota archaeon
TET70900.1 GCA 004377085 F420-dependent methylenetetrahydromethanopterin dehydrogenase Candidatus Bathyarchaeota archaeon
TEU12577.1 GCA 004377395 F420-dependent methylenetetrahydromethanopterin dehydrogenase Candidatus Bathyarchaeota archaeon
Thaumarchaeota
Archaeoglobales
Methanomicrobiales
Methanoflorentaceae
Methanophagales
RLG38457.1 GCA 003661185 F420-dependent methylenetetrahydromethanopterin dehydrogenase Methanosarcinales archaeon
OFV67750.1 GCA 001766815 N5N10-methylenetetrahydromethanopterin dehydrogenase Candidatus Syntrophoarchaeum caldarius
GCA 002011165MTMI01000003.1 6056517573323 60 GCA 002011165 1 ID 3 60 partial 00 Euryarchaeota archaeon JdFR-21
GCA 002505645DAVS01000173.1 28031645141 2 GCA 002505645 1 ID 141 2 partial 00 archaeon UBA460
TET80027.1TET74445.1 GCA 004376735 methylenetetrahydromethanopterin dehydrogenase partial Candidatus Heimdallarchaeota archaeon
GCA 002495885DAHG01000004.1 23211732198542 23 GCA 002495885 -1 ID 42 23 partial 00 Methanocellaceae archaeon UBA148
Methanosarcinales
Methanocellales
Tree scale: 0.1

**Supplementary Figure 39.** ML phylogeny of Mtd. Black circles indicate strongly supported branches (ultrafast bootstrap >=95, aLRT SH-like >=80).

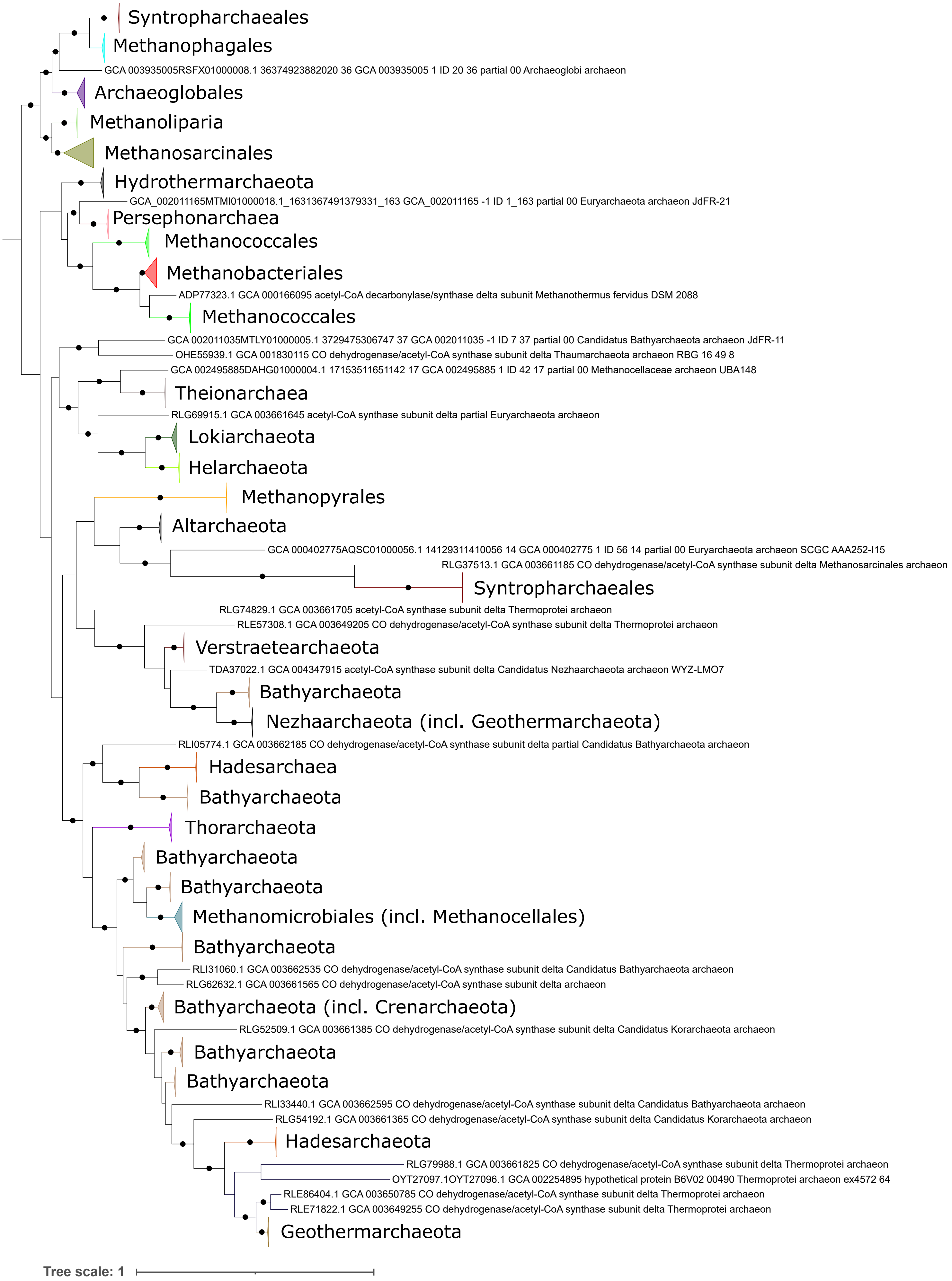
Syntropharchaeales
Methanophagales
GCA 003935005RSFX01000008.1 36374923882020 36 GCA 003935005 1 ID 20 36 partial 00 Archaeoglobi archaeon
Archaeoglobales
Methanoliparia
Methanosarcinales
Hydrothermarchaeota
GCA_002011165MTMI01000018.1_1631367491379331_163 GCA_002011165 -1 ID 1_163 partial 00 Euryarchaeota archaeon JdFR-21
Persephonarchaea
Methanococcales
Methanobacteriales
ADP77323.1 GCA 000166095 acetyl-CoA decarbonylase/synthase delta subunit Methanothermus fervidus DSM 2088
Methanococcales
GCA 002011035MTLY01000005.1 3729475306747 37 GCA 002011035 -1 ID 7 37 partial 00 Candidatus Bathyarchaeota archaeon JdFR-11
OHE55939.1 GCA 001830115 CO dehydrogenase/acetyl-CoA synthase subunit delta Thaumarchaeota archaeon RBG 16 49 8
GCA 002495885DAHG01000004.1 17153511651142 17 GCA 002495885 1 ID 42 17 partial 00 Methanocellaceae archaeon UBA148
Theionarchaea
RLG69915.1 GCA 003661645 acetyl-CoA synthase subunit delta partial Euryarchaeota archaeon
Lokiarchaeota
Helarchaeota
Methanopyrales
Altarchaeota
GCA 000402775AQSC01000056.1 14129311410056 14 GCA 000402775 1 ID 56 14 partial 00 Euryarchaeota archaeon SCGC AAA252-I15
RLG37513.1 GCA 003661185 CO dehydrogenase/acetyl-CoA synthase subunit delta Methanosarcinales archaeon
Syntropharchaeales
RLG74829.1 GCA 003661705 acetyl-CoA synthase subunit delta Thermoprotei archaeon
RLE57308.1 GCA 003649205 CO dehydrogenase/acetyl-CoA synthase subunit delta Thermoprotei archaeon
Verstraetearchaeota
TDA37022.1 GCA 004347915 acetyl-CoA synthase subunit delta Candidatus Nezhaarchaeota archaeon WYZ-LMO7
Bathyarchaeota
Nezhaarchaeota (incl. Geothermarchaeota)
RLI05774.1 GCA 003662185 CO dehydrogenase/acetyl-CoA synthase subunit delta partial Candidatus Bathyarchaeota archaeon
Hadesarchaea
Bathyarchaeota
Thorarchaeota
Bathyarchaeota
Bathyarchaeota
Methanomicrobiales (incl. Methanocellales)
Bathyarchaeota
RLI31060.1 GCA 003662535 CO dehydrogenase/acetyl-CoA synthase subunit delta Candidatus Bathyarchaeota archaeon
RLG62632.1 GCA 003661565 CO dehydrogenase/acetyl-CoA synthase subunit delta archaeon
Bathyarchaeota (incl. Crenarchaeota)
RLG52509.1 GCA 003661385 CO dehydrogenase/acetyl-CoA synthase subunit delta Candidatus Korarchaeota archaeon
Bathyarchaeota
Bathyarchaeota
RLI33440.1 GCA 003662595 CO dehydrogenase/acetyl-CoA synthase subunit delta Candidatus Bathyarchaeota archaeon
RLG54192.1 GCA 003661365 CO dehydrogenase/acetyl-CoA synthase subunit delta Candidatus Korarchaeota archaeon
Hadesarchaeota
RLG79988.1 GCA 003661825 CO dehydrogenase/acetyl-CoA synthase subunit delta Thermoprotei archaeon
OYT27097.1OYT27096.1 GCA 002254895 hypothetical protein B6V02 00490 Thermoprotei archaeon ex4572 64
RLE86404.1 GCA 003650785 CO dehydrogenase/acetyl-CoA synthase subunit delta Thermoprotei archaeon
RLE71822.1 GCA 003649255 CO dehydrogenase/acetyl-CoA synthase subunit delta Thermoprotei archaeon
Geothermarchaeota
Tree scale: 1

**Supplementary Figure 40.** ML phylogeny of CdhD. Black circles indicate strongly supported branches (ultrafast bootstrap >=95, aLRT SH-like >=80).

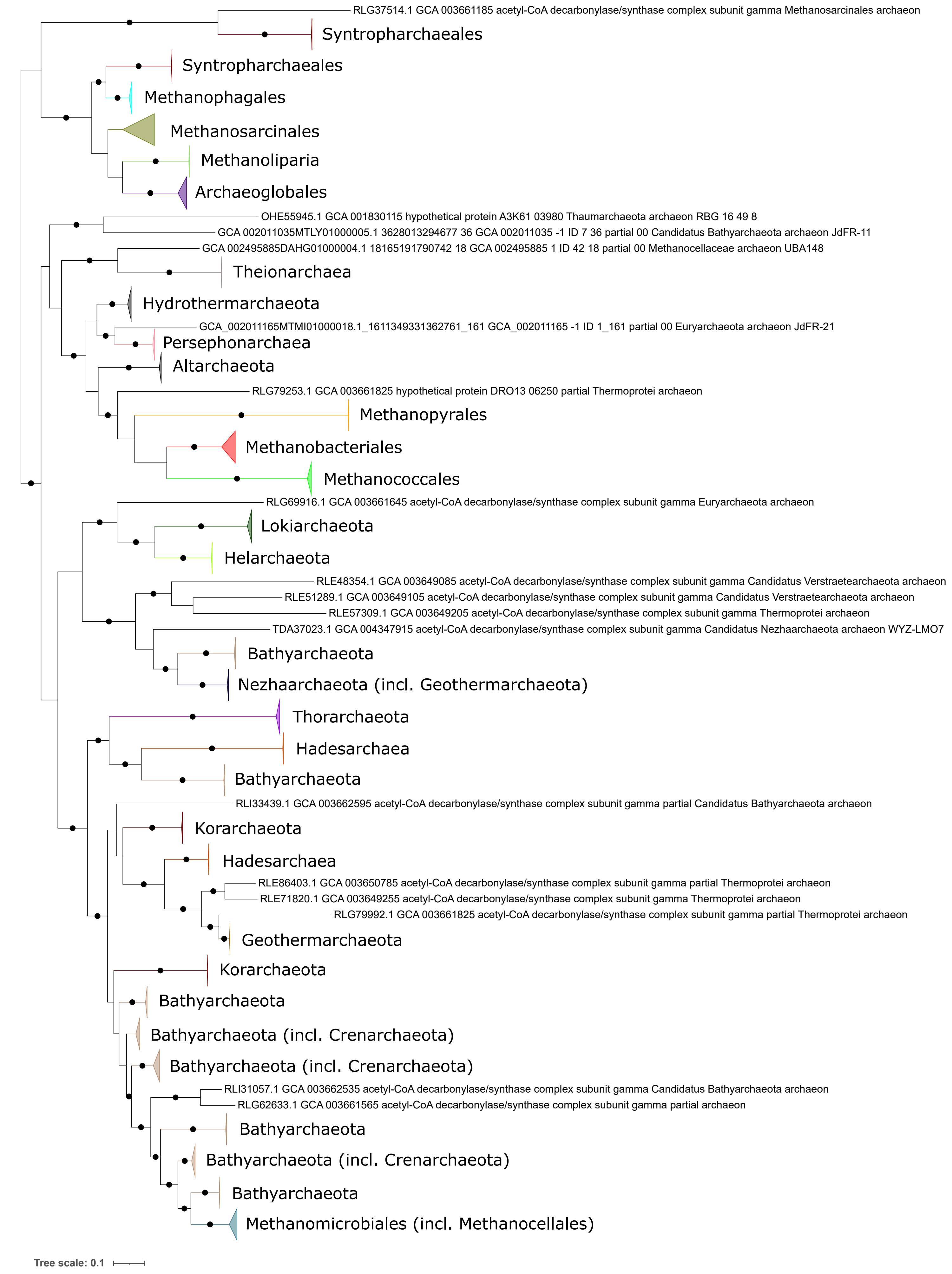
RLG37514.1 GCA 003661185 acetyl-CoA decarbonylase/synthase complex subunit gamma Methanosarcinales archaeon
Syntropharchaeales
Syntropharchaeales
Methanophagales
Methanosarcinales
Methanoliparia
Archaeoglobales
OHE55945.1 GCA 001830115 hypothetical protein A3K61 03980 Thaumarchaeota archaeon RBG 16 49 8
GCA 002011035MTLY01000005.1 3628013294677 36 GCA 002011035 -1 ID 7 36 partial 00 Candidatus Bathyarchaeota archaeon JdFR-11
GCA 002495885DAHG01000004.1 18165191790742 18 GCA 002495885 1 ID 42 18 partial 00 Methanocellaceae archaeon UBA148
Theionarchaea
Hydrothermarchaeota
GCA_002011165MTMI01000018.1_1611349331362761_161 GCA_002011165 -1 ID 1_161 partial 00 Euryarchaeota archaeon JdFR-21
Persephonarchaea
Altarchaeota
RLG79253.1 GCA 003661825 hypothetical protein DRO13 06250 partial Thermoprotei archaeon
Methanopyrales
Methanobacteriales
Methanococcales
RLG69916.1 GCA 003661645 acetyl-CoA decarbonylase/synthase complex subunit gamma Euryarchaeota archaeon
Lokiarchaeota
Helarchaeota
RLE48354.1 GCA 003649085 acetyl-CoA decarbonylase/synthase complex subunit gamma Candidatus Verstraetearchaeota archaeon
RLE51289.1 GCA 003649105 acetyl-CoA decarbonylase/synthase complex subunit gamma Candidatus Verstraetearchaeota archaeon
RLE57309.1 GCA 003649205 acetyl-CoA decarbonylase/synthase complex subunit gamma Thermoprotei archaeon
TDA37023.1 GCA 004347915 acetyl-CoA decarbonylase/synthase complex subunit gamma Candidatus Nezhaarchaeota archaeon WYZ-LMO7
Bathyarchaeota
Nezhaarchaeota (incl. Geothermarchaeota)
Thorarchaeota
Hadesarchaea
Bathyarchaeota
RLI33439.1 GCA 003662595 acetyl-CoA decarbonylase/synthase complex subunit gamma partial Candidatus Bathyarchaeota archaeon
Korarchaeota
Hadesarchaea
RLE86403.1 GCA 003650785 acetyl-CoA decarbonylase/synthase complex subunit gamma partial Thermoprotei archaeon
RLE71820.1 GCA 003649255 acetyl-CoA decarbonylase/synthase complex subunit gamma Thermoprotei archaeon
RLG79992.1 GCA 003661825 acetyl-CoA decarbonylase/synthase complex subunit gamma partial Thermoprotei archaeon
Geothermarchaeota
Korarchaeota
Bathyarchaeota
Bathyarchaeota (incl. Crenarchaeota)
Bathyarchaeota (incl. Crenarchaeota)
RLI31057.1 GCA 003662535 acetyl-CoA decarbonylase/synthase complex subunit gamma Candidatus Bathyarchaeota archaeon
RLG62633.1 GCA 003661565 acetyl-CoA decarbonylase/synthase complex subunit gamma partial archaeon
Bathyarchaeota
Bathyarchaeota (incl. Crenarchaeota)
Bathyarchaeota
Methanomicrobiales (incl. Methanocellales)
Tree scale: 0.1

**Supplementary Figure 41.** ML phylogeny of CdhE. Black circles indicate strongly supported branches (ultrafast bootstrap >=95, aLRT SH-like >=80).

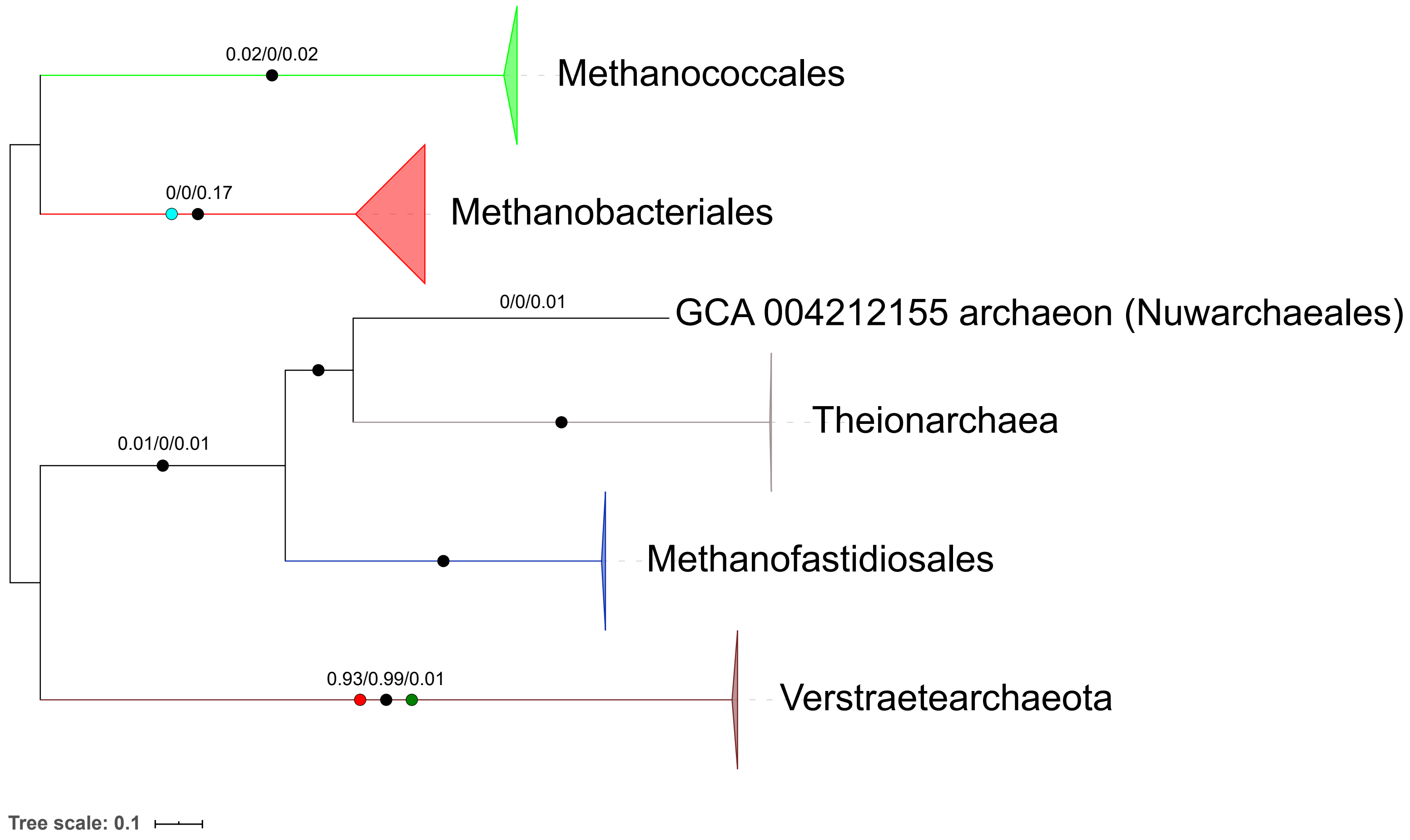
0.02/0/0.02
Methanococcales
0/0/0.17
Methanobacteriales
0/0/0.01
GCA 004212155 archaeon (Nuwarchaeales)
Theionarchaea
0.01/0/0.01
Methanofastidiosales
0.93/0.99/0.01
Verstraetearchaeota
Tree scale: 0.1

**Supplementary Figure 42.** ML phylogeny of EhbEFGHIKLMO (1914 aa positions). Black circles indicate strongly supported branches (ultrafast bootstrap >=95, aLRT SH-like >=80), red circle corresponds to the MAD root, green to MinVar, light blue to NONREV. Branch values correspond to rootstrap supports for MAD, MinVar, and NONREV respectively.

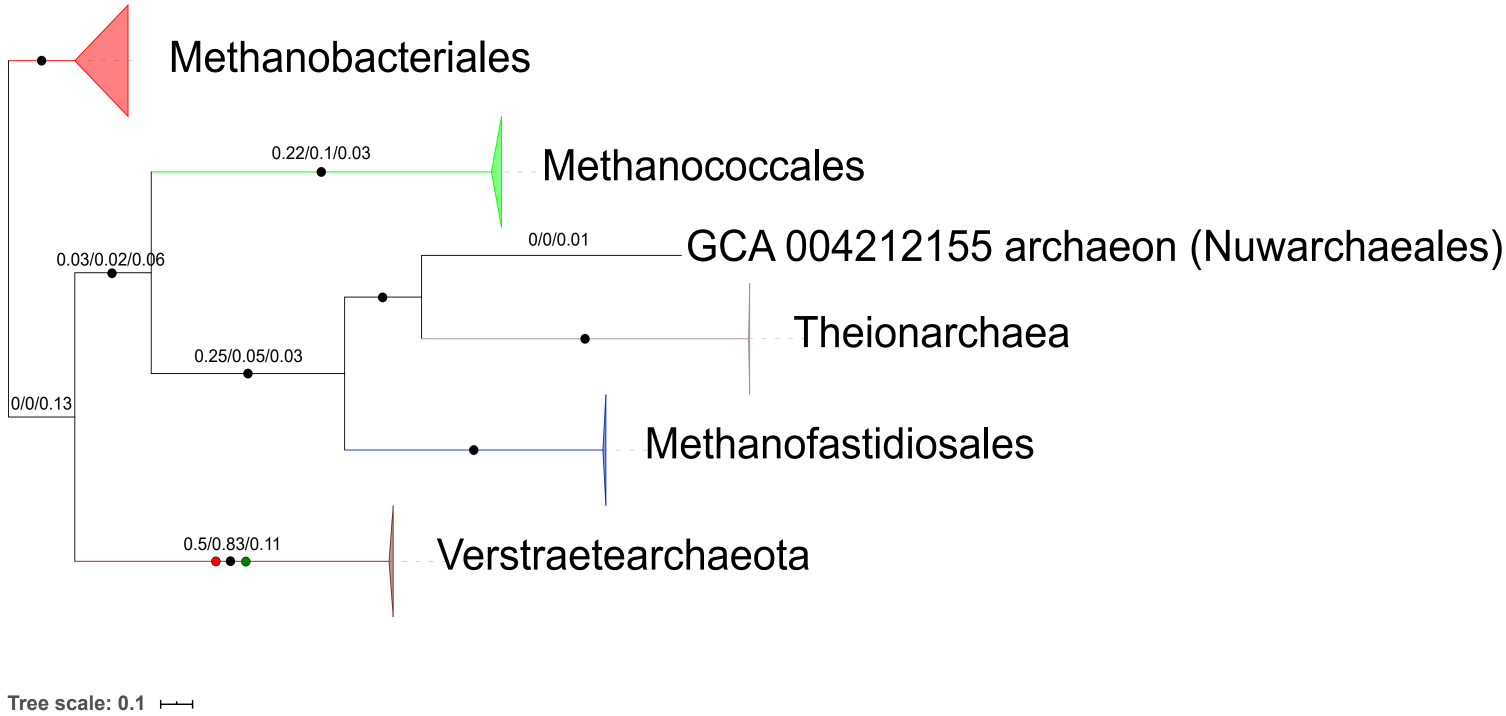
Methanobacteriales
0.22/0.1/0.03
Methanococcales
0/0/0.01
GCA 004212155 archaeon (Nuwarchaeales)
0.03/0.02/0.06
Theionarchaea
0.25/0.05/0.03
0/0/0.13
Methanofastidiosales
0.5/0.83/0.11
Verstraetearchaeota
Tree scale: 0.1

**Supplementary Figure 43.** ML phylogeny of EhbEGHIKLM (1135 aa positions). Black circles indicate strongly supported branches (ultrafast bootstrap >=95, aLRT SH-like >=80), red circle corresponds to the MAD root, green to MinVar. Branch values correspond to rootstrap supports for MAD, MinVar, and NONREV respectively. The NONREV root is within a collapsed clade.

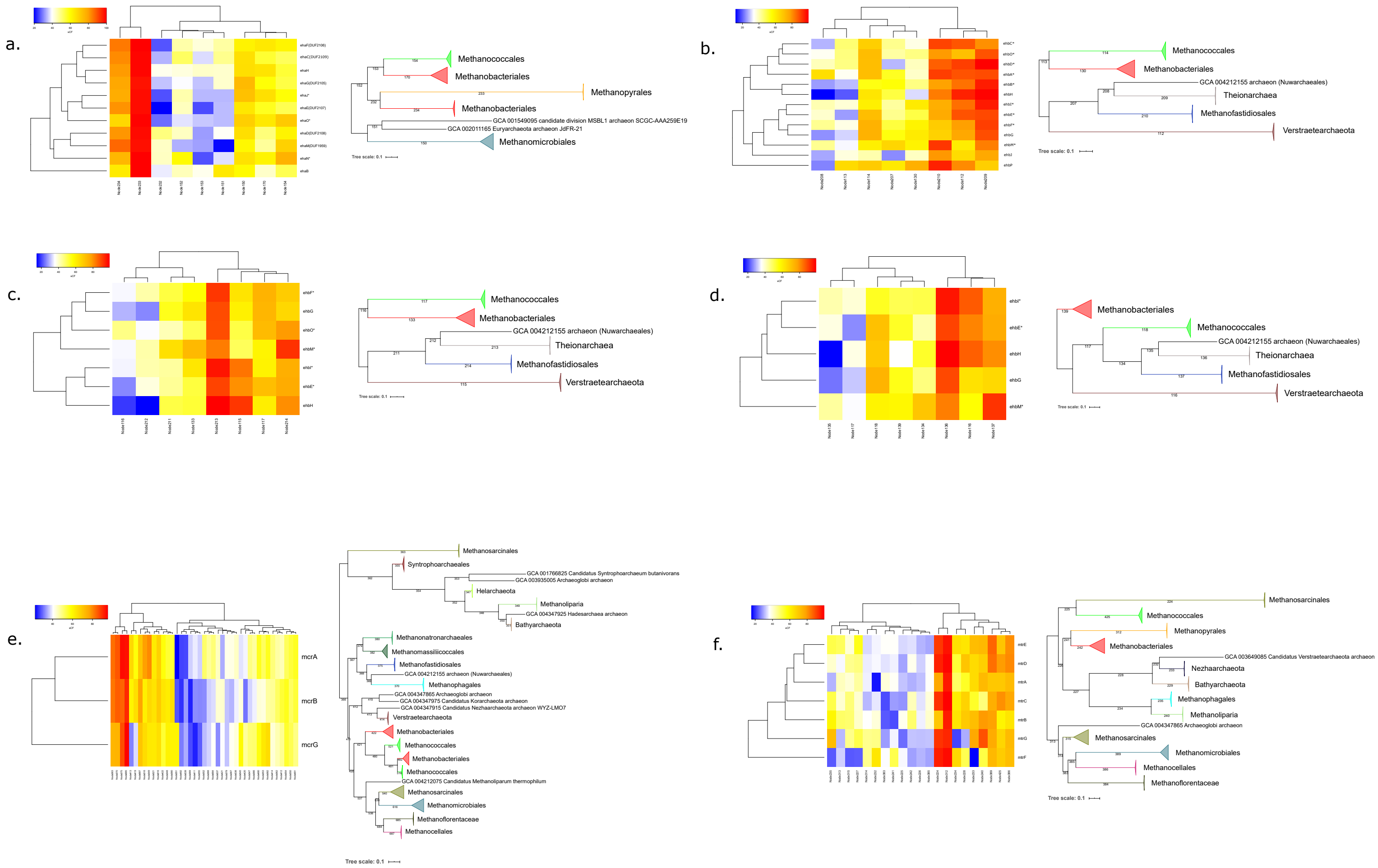

a.
Methanococcales
Methanobacteriales
Methanopyrales
Methanobacteriales
GCA 001549095 candidate division MSBL1 archaeon SCGC-AAA259E19
GCA 002011165 Euryarchaeota archaeon JdFR-21
Methanomicrobiales
Tree scale: 0.1
b.
Methanococcales
Methanobacteriales
GCA 004212155 archaeon (Nuwarchaeales)
Theionarchaea
Methanofastidiosales
Verstraetearchaeota
Tree scale: 0.1
c.
Methanococcales
Methanobacteriales
GCA 004212155 archaeon (Nuwarchaeales)
Theionarchaea
Methanofastidiosales
Verstraetearchaeota
Tree scale: 0.1
d.
Methanobacteriales
Methanococcales
GCA 004212155 archaeon (Nuwarchaeales)
Theionarchaea
Methanofastidiosales
Verstraetearchaeota
Tree scale: 0.1
e.
mcrA
mcrB
mcrG
Methanosarcinales
Syntrophoarchaeales
GCA 001766825 Candidatus Syntrophoarchaeum butanivorans
GCA 003935005 Archaeoglobi archaeon
Helarchaeota
Methanoliparia
GCA 004347925 Hadesarchaea archaeon
Bathyarchaeota
Methanonatronarchaeales
Methanomassiliicoccales
Methanofastidiosales
GCA 004212155 archaeon (Nuwarchaeales)
Methanophagales
GCA 004347865 Archaeoglobi archaeon
GCA 004347975 Candidatus Korarchaeota archaeon
GCA 004347915 Candidatus Nezharchaeota archaeon WYZ-LMO7
Verstraetearchaeota
Methanobacteriales
Methanococcales
Methanobacteriales
Methanococcales
GCA 004212075 Candidatus Methanoliparum thermophilum
Methanosarcinales
Methanomicrobiales
Methanoflorentaceae
Methanocellales
Tree scale: 0.1
f.
Methanosarcinales
Methanococcales
Methanopyrales
Methanobacteriales
GCA 003649085 Candidatus Verstraetearchaeota archaeon
Nezharchaeota
Bathyarchaeota
Methanophagales
Methanoliparia
GCA 004347865 Archaeoglobi archaeon
Methanosarcinales
Methanomicrobiales
Methanocellales
Methanoflorentaceae
Tree scale: 0.1

**Supplementary Figure 44.** Heatmap graphs of subunit-specific site concordance factors for the main clades in the phylogenies of (a) EhaBCDEFGHJMNO, (b) EhbABCDEFGHIJMOP, (c) EhbEFGHIMO, (d) EhbEGHIM, (e) McrABG, (f) MtrABCDEFG. Cladograms were derived from Euclidean distance clustering. Subunits noted with asterisks have known homologs in other hydrogenases (generic subunits). Some subunits are missing from the sCF analyses, as the concatenation dataset needs to, in terms of distribution, perfectly match the datasets of the single subunits. In case major clades of the supermatrix phylogenies were absent in the single subunit phylogenies, those subunits were omitted from the analysis.

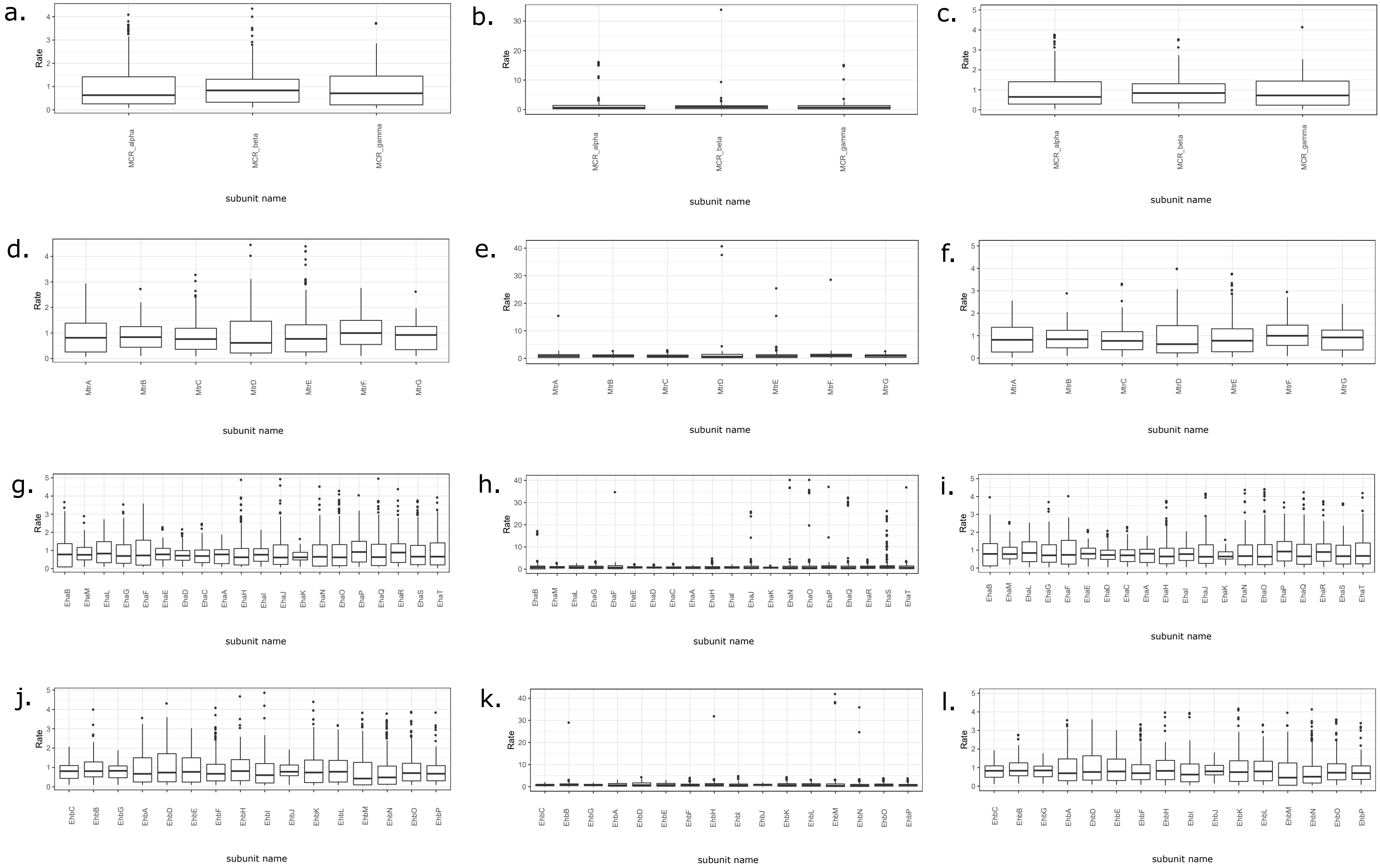

a.
b.
c.
MCR_alpha
MCR_beta
MCR_gamma
d.
e.
f.
MtrA
MtrB
MtrC
MtrD
MtrE
MtrF.
MtrG
g.
h.
i.
EhaB
EhaM
EhaL
EhaG
EhaF
EhaE
EhaD
EhaC
EhaA
EhaH
EhaI
EhaJ
EhaK
EhaN
EhaO
EhaP
EhaQ
EhaR
EhaS
EhaT
j.
k.
l.
EhbC
EhbB
EhbG
EhbA
EhbD
EhbE
EhbF
EhbH
EhbI
EhbJ
EhbK
EhbL
EhbM
EhbN
EhbO
EhbP
Rate
subunit name

**Supplementary Figure 45.** Boxplots for site-specific evolutionary rates for each subunit of (a, b, c) Mcr, (d, e, f) Mtr, (g, h, i) Eha, (j, k, l) Ehb. The plots for each complex correspond to Poisson ML, Poisson+G16 ML, Poisson+G16 empirical Bayesian respectively.

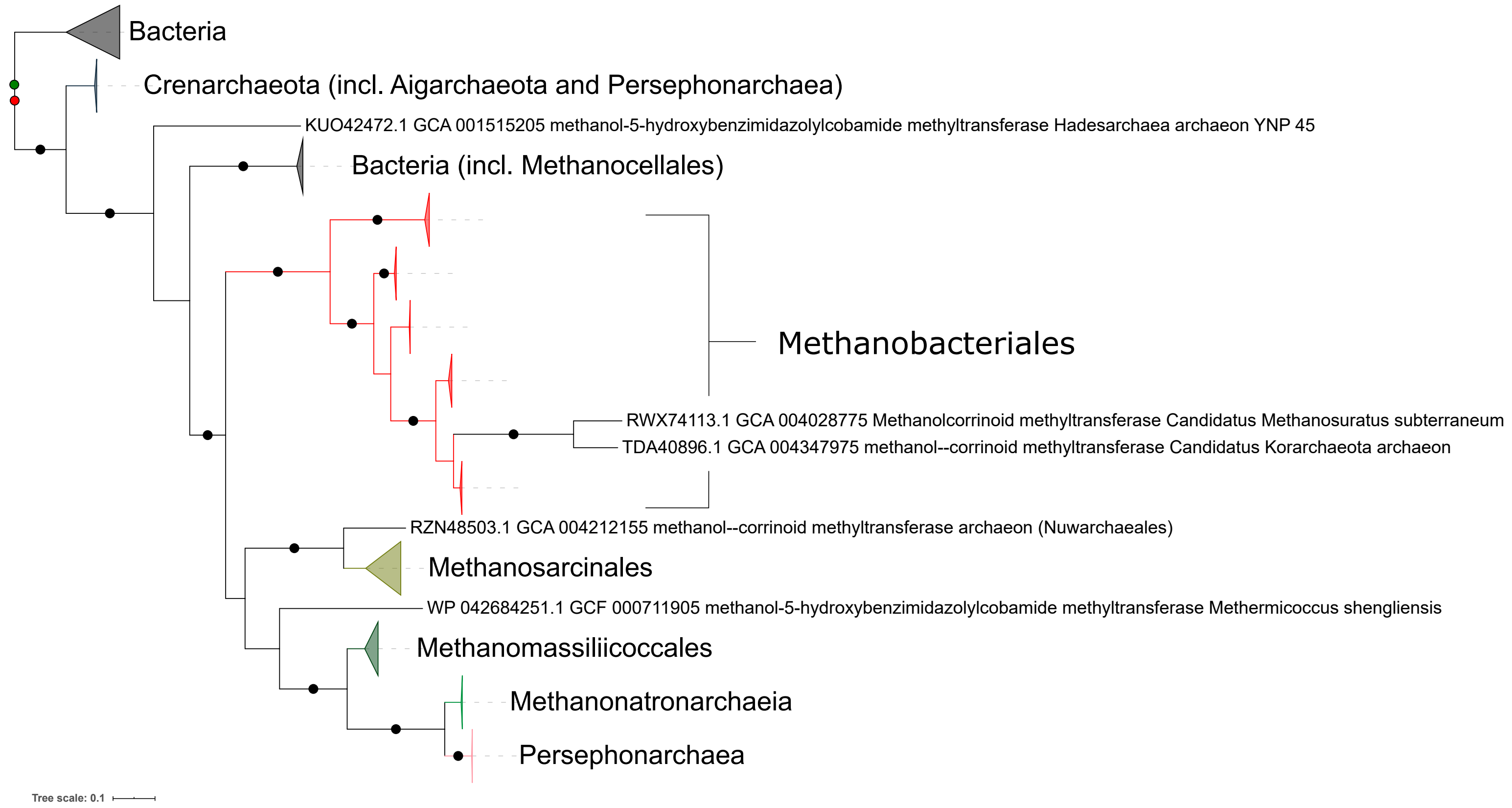
Bacteria
Crenarchaeota (incl. Aigarchaeota and Persephonarchaea)
KUO42472.1 GCA 001515205 methanol-5-hydroxybenzimidazolylcobamide methyltransferase Hadesarchaea archaeon YNP 45
Bacteria (incl. Methanocellales)
Methanobacteriales
RWX74113.1 GCA 004028775 Methanolcorrinoid methyltransferase Candidatus Methanosuratus subterraneum
TDA40896.1 GCA 004347975 methanol--corrinoid methyltransferase Candidatus Korarchaeota archaeon
RZN48503.1 GCA 004212155 methanol--corrinoid methyltransferase archaeon (Nuwarchaeales)
Methanosarcinales
WP 042684251.1 GCF 000711905 methanol-5-hydroxybenzimidazolylcobamide methyltransferase Methermicoccus shengliensis
Methanomassiliicoccales
Methanonatronarchaeia
Persephonarchaea
Tree scale: 0.1

**Supplementary Figure 46.** ML phylogeny of MtaB. Black circles indicate strongly supported branches (ultrafast bootstrap >=95, aLRT SH-like >=80), red circle corresponds to the MAD root, green to MinVar.

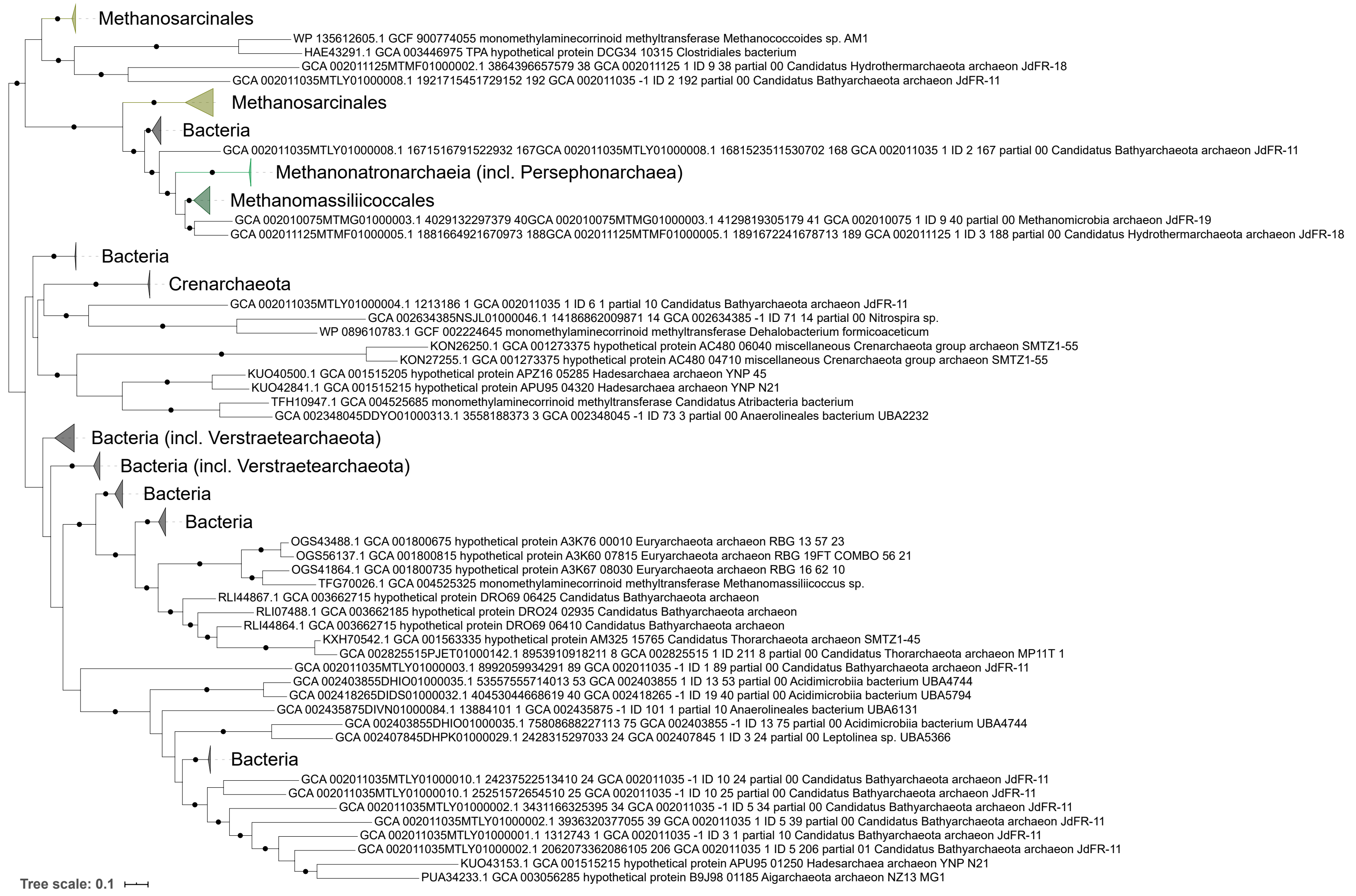

Methanosarcinales
WP 135612605.1 GCF 900774055 monomethylaminecorrinoid methyltransferase Methanococcoides sp. AM1
HAE43291.1 GCA 003446975 TPA hypothetical protein DCG34 10315 Clostridiales bacterium
GCA 002011125MTMF01000002.1 3864396657579 38 GCA 002011125 1 ID 9 38 partial 00 Candidatus Hydrothermarchaeota archaeon JdFR-18
GCA 002011035MTLY01000008.1 1921715451729152 192 GCA 002011035 -1 ID 2 192 partial 00 Candidatus Bathyarchaeota archaeon JdFR-11
Methanosarcinales
Bacteria
GCA 002011035MTLY01000008.1 1671516791522932 167GCA 002011035MTLY01000008.1 1681523511530702 168 GCA 002011035 1 ID 2 167 partial 00 Candidatus Bathyarchaeota archaeon JdFR-11
Methanonatronarchaeia (incl. Persephonarchaea)
Methanomassiliicoccales
GCA 002010075MTMG01000003.1 4029132297379 40GCA 002010075MTMG01000003.1 4129819305179 41 GCA 002010075 1 ID 9 40 partial 00 Methanomicrobia archaeon JdFR-19
GCA 002011125MTMF01000005.1 1881664921670973 188GCA 002011125MTMF01000005.1 1891672241678713 189 GCA 002011125 1 ID 3 188 partial 00 Candidatus Hydrothermarchaeota archaeon JdFR-18
Bacteria
Crenarchaeota
GCA 002011035MTLY01000004.1 1213186 1 GCA 002011035 1 ID 6 1 partial 10 Candidatus Bathyarchaeota archaeon JdFR-11
GCA 002634385NSJL01000046.1 14186862009871 14 GCA 002634385 -1 ID 71 14 partial 00 Nitrospira sp.
WP 089610783.1 GCF 002224645 monomethylaminecorrinoid methyltransferase Dehalobacterium formicoaceticum
KON26250.1 GCA 001273375 hypothetical protein AC480 06040 miscellaneous Crenarchaeota group archaeon SMTZ1-55
KON27255.1 GCA 001273375 hypothetical protein AC480 04710 miscellaneous Crenarchaeota group archaeon SMTZ1-55
KUO40500.1 GCA 001515205 hypothetical protein APZ16 05285 Hadesarchaea archaeon YNP 45
KUO42841.1 GCA 001515215 hypothetical protein APU95 04320 Hadesarchaea archaeon YNP N21
TFH10947.1 GCA 004525685 monomethylaminecorrinoid methyltransferase Candidatus Atribacteria bacterium
GCA 002348045DDYO01000313.1 3558188373 3 GCA 002348045 -1 ID 73 3 partial 00 Anaerolineales bacterium UBA2232
Bacteria (incl. Verstraetearchaeota)
Bacteria (incl. Verstraetearchaeota)
Bacteria
Bacteria
OGS43488.1 GCA 001800675 hypothetical protein A3K76 00010 Euryarchaeota archaeon RBG 13 57 23
OGS56137.1 GCA 001800815 hypothetical protein A3K60 07815 Euryarchaeota archaeon RBG 19FT COMBO 56 21
OGS41864.1 GCA 001800735 hypothetical protein A3K67 08030 Euryarchaeota archaeon RBG 16 62 10
TFG70026.1 GCA 004525325 monomethylaminecorrinoid methyltransferase Methanomassiliicoccus sp.
RLI44867.1 GCA 003662715 hypothetical protein DRO69 06425 Candidatus Bathyarchaeota archaeon
RLI07488.1 GCA 003662185 hypothetical protein DRO24 02935 Candidatus Bathyarchaeota archaeon
RLI44864.1 GCA 003662715 hypothetical protein DRO69 06410 Candidatus Bathyarchaeota archaeon
KXH70542.1 GCA 001563335 hypothetical protein AM325 15765 Candidatus Thorarchaeota archaeon SMTZ1-45
GCA 002825515PJET01000142.1 8953910918211 8 GCA 002825515 1 ID 211 8 partial 00 Candidatus Thorarchaeota archaeon MP11T 1
GCA 002011035MTLY01000003.1 8992059934291 89 GCA 002011035 -1 ID 1 89 partial 00 Candidatus Bathyarchaeota archaeon JdFR-11
GCA 002403855DHIO01000035.1 53557555714013 53 GCA 002403855 1 ID 13 53 partial 00 Acidimicrobiia bacterium UBA4744
GCA 002418265DIDS01000032.1 40453044668619 40 GCA 002418265 -1 ID 19 40 partial 00 Acidimicrobiia bacterium UBA5794
GCA 002435875DIVN01000084.1 13884101 1 GCA 002435875 -1 ID 101 1 partial 10 Anaerolineales bacterium UBA6131
GCA 002403855DHIO01000035.1 75808688227113 75 GCA 002403855 -1 ID 13 75 partial 00 Acidimicrobiia bacterium UBA4744
GCA 002407845DHPK01000029.1 2428315297033 24 GCA 002407845 1 ID 3 24 partial 00 Leptolinea sp. UBA5366
Bacteria
GCA 002011035MTLY01000010.1 24237522513410 24 GCA 002011035 -1 ID 10 24 partial 00 Candidatus Bathyarchaeota archaeon JdFR-11
GCA 002011035MTLY01000010.1 25251572654510 25 GCA 002011035 -1 ID 10 25 partial 00 Candidatus Bathyarchaeota archaeon JdFR-11
GCA 002011035MTLY01000002.1 3431166325395 34 GCA 002011035 -1 ID 5 34 partial 00 Candidatus Bathyarchaeota archaeon JdFR-11
GCA 002011035MTLY01000002.1 3936320377055 39 GCA 002011035 1 ID 5 39 partial 00 Candidatus Bathyarchaeota archaeon JdFR-11
GCA 002011035MTLY01000001.1 1312743 1 GCA 002011035 -1 ID 3 1 partial 10 Candidatus Bathyarchaeota archaeon JdFR-11
GCA 002011035MTLY01000002.1 2062073362086105 206 GCA 002011035 1 ID 5 206 partial 01 Candidatus Bathyarchaeota archaeon JdFR-11
KUO43153.1 GCA 001515215 hypothetical protein APU95 01250 Hadesarchaea archaeon YNP N21
PUA34233.1 GCA 003056285 hypothetical protein B9J98 01185 Aigarchaeota archaeon NZ13 MG1
Tree scale: 0.1

**Supplementary Figure 47.** ML phylogeny of MtmB. Black circles indicate strongly supported branches (ultrafast bootstrap >=95, aLRT SH-like >=80), red circle corresponds to the MAD root, green to MinVar.

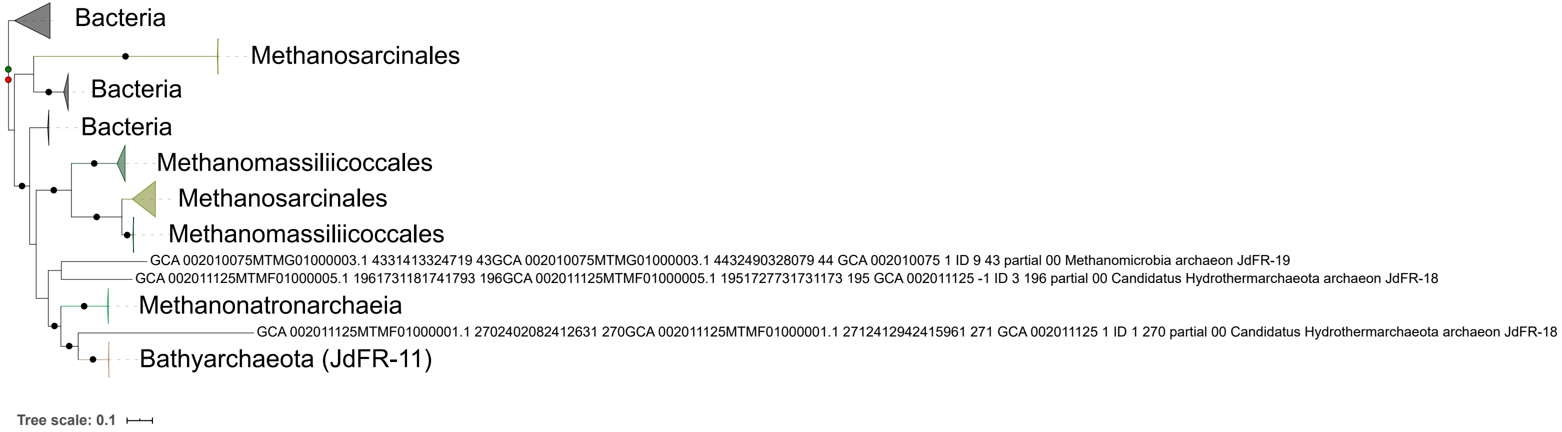

Bacteria
Methanosarcinales
Bacteria
Bacteria
Methanomassiliicoccales
Methanosarcinales
Methanomassiliicoccales
GCA 002010075MTMG01000003.1 4331413324719 43GCA 002010075MTMG01000003.1 4432490328079 44 GCA 002010075 1 ID 9 43 partial 00 Methanomicrobia archaeon JdFR-19
GCA 002011125MTMF01000005.1 1961731181741793 196GCA 002011125MTMF01000005.1 1951727731731173 195 GCA 002011125 -1 ID 3 196 partial 00 Candidatus Hydrothermarchaeota archaeon JdFR-18
Methanonatronarchaeia
GCA 002011125MTMF01000001.1 2702402082412631 270GCA 002011125MTMF01000001.1 2712412942415961 271 GCA 002011125 1 ID 1 270 partial 00 Candidatus Hydrothermarchaeota archaeon JdFR-18
Bathyarchaeota (JdFR-11)
Tree scale: 0.1

**Supplementary Figure 48.** ML phylogeny of MtbB. Black circles indicate strongly supported branches (ultrafast bootstrap >=95, aLRT SH-like >=80), red circle corresponds to the MAD root, green to MinVar.

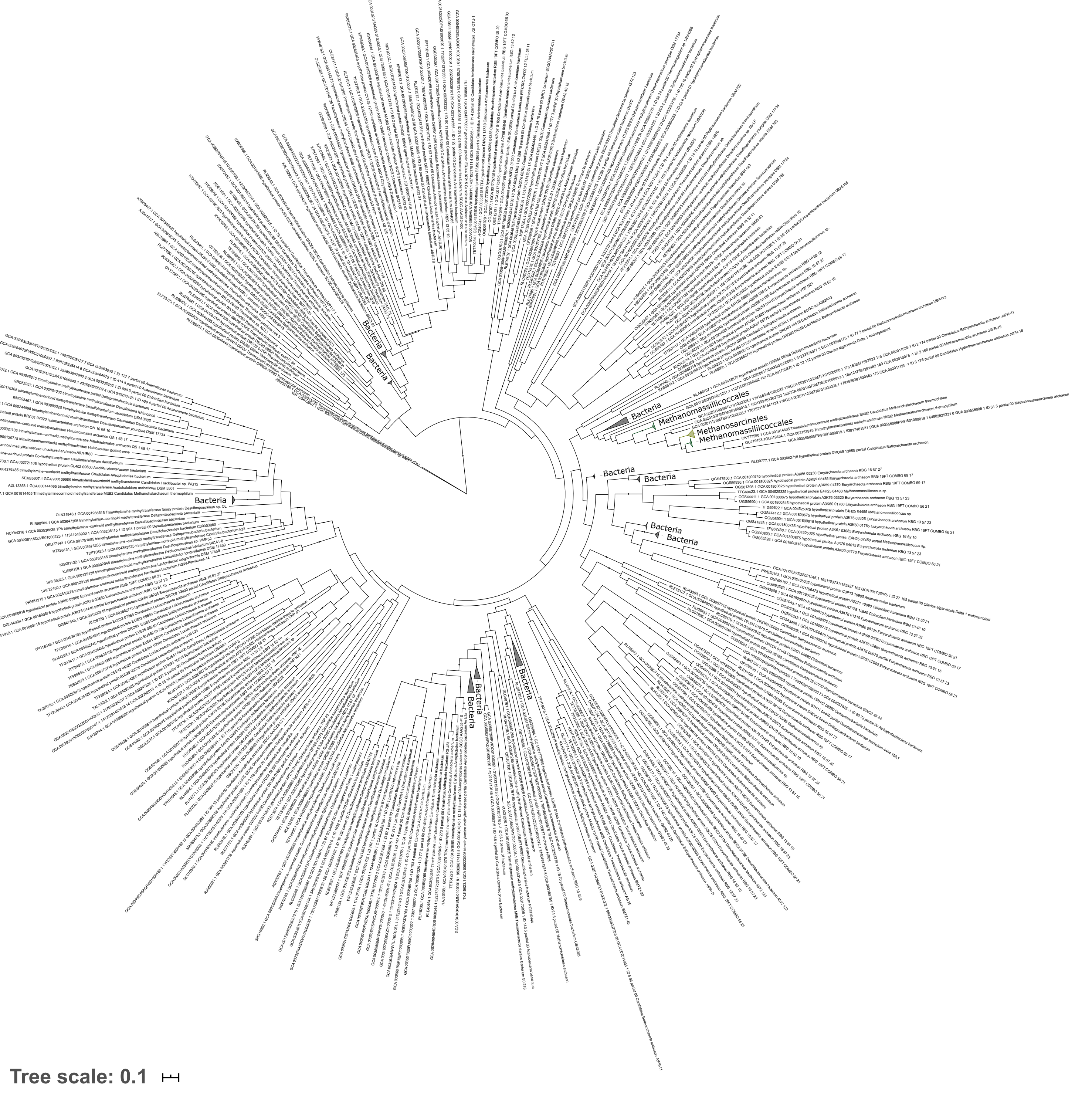
Bacteria
Methanomassiliicoccales
Methanosarcinales
Tree scale: 0.1

**Supplementary Figure 49.** ML phylogeny of MttB. Black circles indicate strongly supported branches (ultrafast bootstrap >=95, aLRT SH-like >=80).

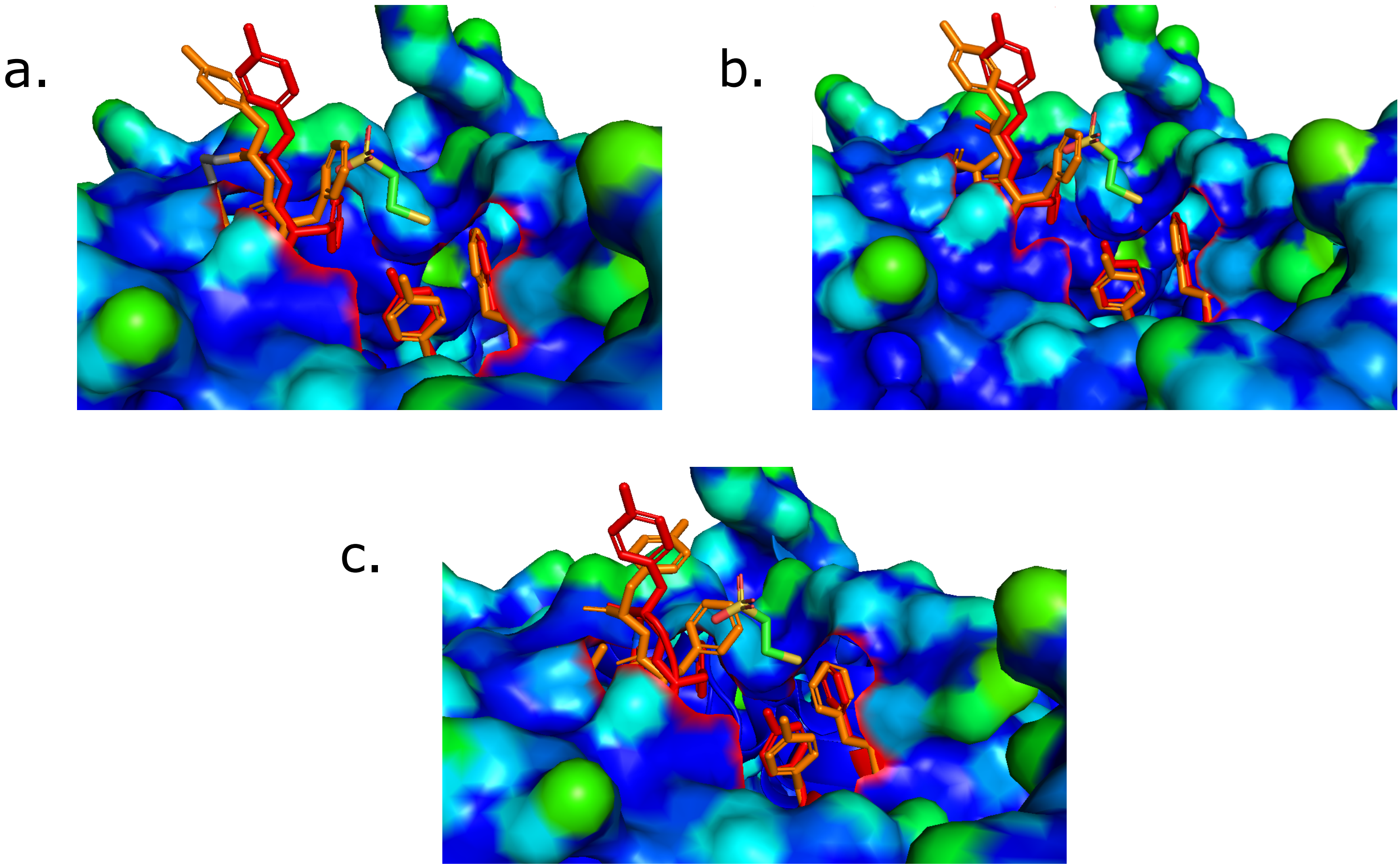
a.
b.
c.

**Supplementary Figure 50.** Homology model of the ancestral McrA sequence (orange) with the a) Mcr-like, b) Halobacterota, c) Proteoarchaeota roots aligned with the template structure of the *M. marburgensis* McrA (PDB ID: 1HBU, chain D; red). The view is focused on the methyl-CoM binding cavity.
